# CanSeer: A Method for Development and Clinical Translation of Personalized Cancer Therapeutics

**DOI:** 10.1101/2022.06.29.498138

**Authors:** Rida Nasir Butt, Bibi Amina, Muhammad Umer Sultan, Zain Bin Tanveer, Risham Hussain, Rida Akbar, Salaar Khan, Mahnoor Naseer Gondal, Muhammad Farhan Khalid, Amir Faisal, Muhammad Shoaib, Safee Ullah Chaudhary

## Abstract

Computational modeling and analysis of biomolecular network models annotated with cancer patient-specific multi-omics data can enable the development of personalized therapies. Current endeavors aimed at employing *in silico* models towards personalized cancer therapeutics remain to be fully translated. In this work, we present “*CanSeer*” a novel multi-stage methodology for developing *in silico* models towards clinical translation of personalized cancer therapeutics. The proposed methodology integrates state-of-the-art dynamical analysis of biomolecular network models with patient-specific genomic and transcriptomic data to assess the individualized therapeutic responses to targeted drugs and their combinations. *CanSeer’s* translational approach employs transcriptomic data (RNA-seq based gene expressions) with genomic profile (CNVs, SMs, and SVs). Specifically, patient-specific cancer driver genes are identified, followed by the selection of druggable and/or clinically actionable targets for therapeutic interventions. To exemplify *CanSeer*, we have designed three case studies including (i) lung squamous cell carcinoma, (ii) breast invasive carcinoma, and (iii) ovarian serous cystadenocarcinoma. The case study on lung squamous cell carcinoma concluded that restoration of Tp53 activity together with an inhibition of EGFR as an efficacious combinatorial treatment for patients with Tp53 and EGFR cancer driver genes. The findings from the cancer case study helped identify personalized treatments including APR-246, APR-246+palbociclib, APR-246+osimertinib, APR-246+afatinib, APR-246+osimertinib+dinaciclib, and APR-246+afatinib+dinaciclib. The second case study on breast invasive carcinoma revealed *CanSeer*’s potential to elucidate drug resistance against targeted drugs and their combinations including KU-55933, afuresertib, ipatasertib, and KU-55933+afuresertib. Lastly, the ovarian cancer case study revealed the combinatorial efficacy of APR-246+carmustine, and APR-246+dinaciclib for treating ovarian serous cystadenocarcinoma. Taken together, *CanSeer* outlines a novel method for systematic identification of optimal tailored treatments with mechanistic insights into patient-to-patient variability of therapeutic response, drug resistance mechanism, and cytotoxicity profiling towards personalized medicine.

## INTRODUCTION

Cancer remains a leading cause of death worldwide despite an ever-increasing repertoire of treatment modalities [1]–[4]. Amongst the salient impediments faced by anti-cancer regimens are drug resistance [5], patient-to-patient variability of therapeutic response [6], and cytotoxicity [7], [8]. Towards overcoming these issues, molecular targeting-based therapies can offer invaluable support and enhance therapeutic efficacy [9]–[11]. Such molecular targeting in cancer cells was first reported in 1998 with the development of imatinib (Abl tyrosine kinase inhibitor) for treating chronic myeloid leukemia [12]. FDA’s approval of imatinib in 2001 [12] established the plausibility and clinical potential of targeted therapies aimed at specific molecular targets. The ensuing research impetus led to the development of several targeted therapies including vemurafenib for targeting BRAF mutations [13]–[15], gilteritinib to target FLT3 mutations [16], [17], and temsirolimus for inhibiting mammalian target of rapamycin (mTOR) [18], [19], etc. These breakthrough drugs further stimulated efforts aimed at identification of molecular targets in wider varieties of cancer and onward employment in devising targeted therapies [20]–[24].

Advancements in high throughput sequencing [25] and multiplex mutational screening [26] have been instrumental in elucidating novel molecular signatures for use in designing targeted therapies [25]–[29]. Furthermore, information on genetic aberrations in individual patients is now informing targeted therapies and has given rise to the domain of personalized therapeutics [30]–[33]. In an early-harvest project, MD Anderson Cancer Center embarked on a program to screen patient-specific genetic alterations with “matched targeted therapies” [34]–[36]. Follow-up clinical trials gave momentum to genome-informed precision medicine and highlighted its potential for clinical adoption [37]–[40]. Notably, most of such trials remained limited to monotherapies (a single target - a single drug combination) [13]–[18], [29], [37] which eventually also started to lose efficacy and develop resistance. As an example, about 65% of FLT3 mutated refractory acute myeloid leukemia (AML) patients showed resistance to gilteritinib [17]. Similarly, BRAF targeting single-agent vemurafenib failed to offer benefits during phase-II clinical trials [37]. This was partially attributable to drug resistance caused by underlying genetic heterogeneity [41] and molecular cross-talk between interconnected signaling pathways [42], thereby escaping treatment [43], [44]. Latest endeavors at overcoming these undesired outcomes are now employing combinations of monotherapies such as trametinib+fluvastatin for treating lung cancer [45], and trametinib+zoledronate for KRAS-mutated patients of metastatic colorectal cancer [46], etc. Results from such combinatorial regimens have demonstrated lowered rates of treatment escape and resistance to therapy [47], [48].

Towards predicting effective combinatorial therapies, as well as to better understand their mechanistic underpinnings, biomolecular network models and their dynamical simulations have gained prominence [49]–[55]. Over the last two decades, several applications of mathematical modeling and computational dynamical analysis have helped elucidate the dynamics of human biomolecular signaling network models [56]–[63]. In recent years, incorporation of patient-specific multi-omics data into biomolecular network models has further provided key insights into patient tumors as well as enabled the orchestration of patient-centric interventions. Taking this approach, Hofree *et al*. analyzed patient mutations to cluster within lung, ovarian, and uterine cancer cohorts and predicted patient survival and clinical outcomes [64]. In a large-scale study, Hidalgo *et al* annotated 60 KEGG pathways with patient-specific genomic and transcriptomic data from 12 cancer types and decomposed the pathways into circuits for predicting cell activity and survival rate [55]. Likewise, Fey *et al* reported a c-Jun N-terminal kinase (JNK) activation switch in neuroblastoma cells and performed patient-specific simulations to predict survival [53].

Recently, the biomolecular modeling approach was tailored to patient tumor samples capitalizing on omics profiles including expression data (transcriptomics or proteomics), copy number alterations, and mutation data to stratify patients into clinical sub-groups [52]. Although these patient-specific models were geared toward precision medicine, the studies focused on clinical sub-grouping and individualized survival analyses. In a recent study, patient-specific dynamical models were constructed by coupling the continuous logic model of extrinsic and intrinsic apoptosis with microfluidic-based screenings of patients’ cancer biopsies [65]. The consequent personalized models managed to uncover the heterogeneity amongst pancreatic cancer patients, but remained limited to only a single “apoptotic” readout. Later, Ianevski *et al* implemented a machine learning approach to identify patient-specific synergistic combinatorial regimens directly from online drug databases, and reported reduced toxicity during co-inhibition of leukemic cells [66]. The study proposed coupling of *ex vivo* monotherapy screening profiles with single-cell RNA sequencing data of patient samples. However, the machine learning approach did not provide mechanistic or interpretable insights into patient tumors and treatment [67]. Here, it is important to note that, the direct employment and consumption of drug databases such as PanDrugs [68], Drug Target Commons [69], and DGIdb [70]–[72] etc. can also produce off-label drug targeting. The off-label drug use is due to the presence of clinically rejected drugs (e.g. AZD-5438 (CDK inhibitor [73]) etc.) in online drug databases. In a recent study, a personalized *in silico* “*drosophila patient model*” employed patient-specific somatic mutations, and identified personalized combinatorial treatment for colorectal cancer [74]. The proposed annotation of network models with somatic mutations remained deficient in construing a comprehensive molecular signature of individual patients. Moreover, the coverage of druggable and clinically actionable targets remained limited during exhaustive screening for predicting efficacious targets [74], [75]. To date, a multi-factorial patient data-integrative approach to simulate drug effects using drug scores, followed by quantification of therapy response remains unassessed. In addition, the lack of a systematic approach that facilitates the identification of optimal treatment, drug resistance mechanism, and cytotoxicity evaluation impedes devising of personalized therapies. As a result, the advantages offered by *in silico* studies toward personalized cancer therapeutics remain to be fully translated [76], thereby, necessitating a systematic method that elucidates the clinical translation of modeling-based personalized therapeutics.

In this work, we propose a method “*CanSeer*” to develop *in silico* models for clinical translation of personalized treatments. The proposed method initiates with the development of literature-derived biomolecular regulatory network models followed by their validation. Next, each model is evaluated against random and systematic perturbations to assess the robustness and biological plausibility of the abstraction. The validated model is employed to integrate patient-specific genomic and transcriptomic data. For that, omics data is acquired, pre-processed, normalized, and then subjected to the model. Subsequently, the dynamical analysis of the patient-centric model is performed. To establish the clinical relevance of personalized treatment, cancer genetic drivers in patients are identified. Next, prioritizing therapeutic screening to be informative for clinical translation, druggable and clinically actionable targets are selected. With this, the drugs targeting these molecular signatures, either directly or indirectly, are manually curated for molecularly and clinically informed use of drug databases. Next, drug activity score is computed to divulge the effect of different drugs targeting the same gene. Moreover, to decode the varying effect of a drug from patient to patient, *CanSeer* computes “drug scores for patient (DS)” that employs the potential activity of candidate drugs and normalized gene expressions of a patient in cancer. Patient-specific drug scores are then used to annotate the personalized models, and the models are re-analyzed.

Employing the resultant cell fate outcomes, patient-specific therapeutic response is computed together with cytotoxicity evaluation. Patient-specific therapeutic response, and cytotoxicity combination cues the optimal treatment options for cancer patient. Lastly, the cell fates of predicted optimal tailored treatment are compared with the cell fates of the patient’s cancer model to evaluate whether the treatment is “beneficial” or “resistant”. Thereby, increasing the chance to be clinically informative. Case studies performed on (i) lung squamous cell carcinoma, (ii) breast invasive carcinoma, and (iii) ovarian serous cystadenocarcinoma, elucidated patient-specific response to therapies and their combinations. The exemplars helped identify optimal personalized treatment including novel drug combinations such as, APR-246+palbociclib, APR-246+osimertinib, APR-246+afatinib, APR-246+carmustine, APR-246+dinaciclib, APR-246+osimertinib+dinaciclib, and APR-246+afatinib+dinaciclib. Moreover, the cases studies also assisted in identifying drug resistance against targeted drugs and their combinations in individual cancer patients.

Taken together, *CanSeer* provides a clinical translation framework for personalized cancer therapeutics by combining mechanistic insights from logical modeling with comprehensive “multi-omics” patient data. The systematic approach provides assessment of individualized drug effects through drug scores, as well as quantify patient-specific therapeutic response to targeted drugs and their combinations. As a result, the method helps elicit (i) optimal tailored treatments, (ii) mechanistic insights into patient-to-patient variability of therapeutic response, (iii) drug resistance, and (iv) cytotoxicity. In conclusion, the work provides a translational approach to precision oncology for formulating and assessing clinically deployable personalized treatment plans.

## MATERIALS and METHODS

### A. Biomolecular Network Model Construction

To develop the biomolecular regulatory network architecture, pathways and interactions were retrieved from existing databases including Kyoto Encyclopedia of Genes and Genomes (KEGG) [77], PathBank [78], Pathway Interaction Database (PID) [79], [80], and Reactome [81]. Pathway integration and cross-talk were then incorporated into the network architecture. Rules [82]–[84] and weight-based [85], [86] formalisms were adopted for translating the network topology towards carrying out network analyses. Case study exemplars were developed around a literature-based human signaling network comprising of 197 nodes and their interactions was obtained [59]. Boolean logic rules were defined to translate the adopted network into a rules-based model. For developing a weight-based version of rules-based biomolecular regulations, the basal value of each network node was set at 1. Next, interaction weights were computed based on the number of adjacent nodes such that the output of weight-based model becomes comparable to the rules-based model.

### B. Input and Output Node Setup using Fixed Node States and Cell Fate Programming

To analyze the human signaling network under normal ambient conditions, values for input nodes were abstracted from published literature (Supplementary Table 1). Each input value was “fixed” to cater for inputs such that the nodes’ state remains unchanged during dynamical analysis. The normalized patient gene expression for each input node was selected to personalize the network model. In the personalized cancer model, patient’s genetic alterations were also fixed as “fixed node states”. To program the cell fates, cell fate determining biomarkers and the associated network nodes’ states were then acquired from the published literature (Supplementary Table 2). Output nodes and their associated nodes’ states defining cellular functions such as apoptosis, senescence, and cell cycle arrest, etc. were abstracted from literature to further expand the set of cell fates (Supplementary Table 3). The cell fate classification program was adjusted in the light of normalized RNA-seq based gene expression data of patients to personalize the models. Combinations of nodes’ states were created to provide an exhaustive coverage for the cell fate classification scenario (Supplementary Table 4).

### C. Dynamical Deterministic Analysis of Biomolecular Networks

To analyze all the network models, Deterministic Analysis (DA) was performed using a web-based network analysis tool, Theatre for *in silico* Systems Oncology (TISON) [87]. Two files including fixed node states file, and cell fate classification file were input in addition to the network, to execute DA. The initial input conditions were introduced in the fixed node states file to begin DA of a network. The maximum iterations to search for the steady state were set to 500. Subsequently, a bootstrap of 256 network states was applied. Outputs from DA were used to map the cell fates associated with emergent network states. Drug data was integrated, and DA was performed using TISON to undertake therapeutic evaluation.

### D. Robustness Analysis and Parameter Sensitivity Analyses

To determine the susceptibility of biomolecular network to minor variations, the network inputs were perturbed by up to 10%. The input conditions were taken in combination for network analyses. A random sample of 256 network states was selected along with 10% of fixed node combinations. Statistical mechanics [61], [88], [89] was applied to compute average node propensities, standard deviation and standard error of mean (SEM) for each mapped cell fate. The average propensities, and SEMs of emergent cellular phenotypes were plotted to evaluate the stability of biologically plausible network.

To analyze the network behavior under large input perturbations, the adopted network with cell fate expansion was exposed to all possible input values (parameters), sequentially. Each input was assessed at uniform increment of 0.1 from 0 to 1. In order to see the influence of each parameter, all other inputs were kept at normal levels. For that, DA for each input was performed with bootstrap of 256 network states. Results from each input node perturbation were compiled to see the overall effect of each input on network. The interpretation of each input-cell fate relationship was validated against literature. Additionally, multiple input perturbations were performed to investigate the synergistic or antagonistic relationship of stimuli in human signaling network. The levels of co-occurring inputs were varied, and the effect on associated cell fates was stored. For combinatorial parameter sensitivity assessment, DA was performed using TISON [87].

### E. Genomic and Transcriptomic Data Assembly, Pre-processing, Normalization and Model Personalization

To acquire patients’ transcriptomic profiles (RNA-seq based gene expression data), The Cancer Genome Atlas (TCGA) PanCancer Atlas [90], [91] studies were accessed using Genomic Data Commons (GDC) [92] portal. For obtaining the genomic data including copy number variations (CNVs), somatic mutations (SMs), and genomic structural variants (SVs), the same TCGA project was queried in cBioPortal [93], [94]. Patient data matching was ensured by comparing sample IDs between GDC and cBioPortal. Patients’ genomic and transcriptomic data were obtained for the following projects: TCGA-LUSC, TCGA-BRCA, and TCGA-OV to demonstrate the proposed method “*CanSeer*” on 3 different case studies. MATLAB 2020b [95] was used to implement the pre-processing and normalization of genomic, and transcriptomic data. The normalized RNA-seq based gene expression data was used to annotate the network model.

### F. Target Identification using Drug Databases

Drugs targeting each node (biomolecule) in the network were obtained from published literature, and drug-target databases such as PanDrugs [68], DrugBank [96], and The Drug Gene Interaction Database (DGIdb) [70]–[72]. Only druggable and clinically actionable nodes were selected to enable the clinical translation of resultant models. The drugs identified to target druggable or clinically actionable nodes were then queried in Genomics of Drug Sensitivity in Cancer (GDSC2) [97] to obtain an inhibitory concentration of drugs (IC_50_). The IC_50_ values of candidate drugs were acquired for A549 cell line, BT-474 cell line, and OVCAR-3 cell line. Next, IC_50_ values were utilized to formulate the drug activity score (DAS). DAS was then normalized, and utilized to compute a “drug score for patient” (DS) which was then employed to annotate the personalized model. The cell fate outcomes from resultant model’s DA were then used to calculate patient-specific therapeutic response index (TRI), and cytotoxicity.

## RESULTS

“*CanSeer*” is a novel multi-stage methodology for developing *in silico* models towards designing personalized cancer therapeutics. The proposed framework comprises of four salient stages, which include: (i) development of literature-derived Boolean model of biomolecular regulatory network and its validation from literature, (ii) genomic and transcriptomic data collection, pre-processing, and normalization, (iii) network annotation with patient-specific omics data comprising of genomic and transcriptomic profiles followed by the dynamical analyses of personalized models, and (iv) therapeutic screening and identification of optimal regimens for cancer patients.

The overall framework of *CanSeer* along with the stage-wise details (Figure 1) are provided below.

**Figure 1:**
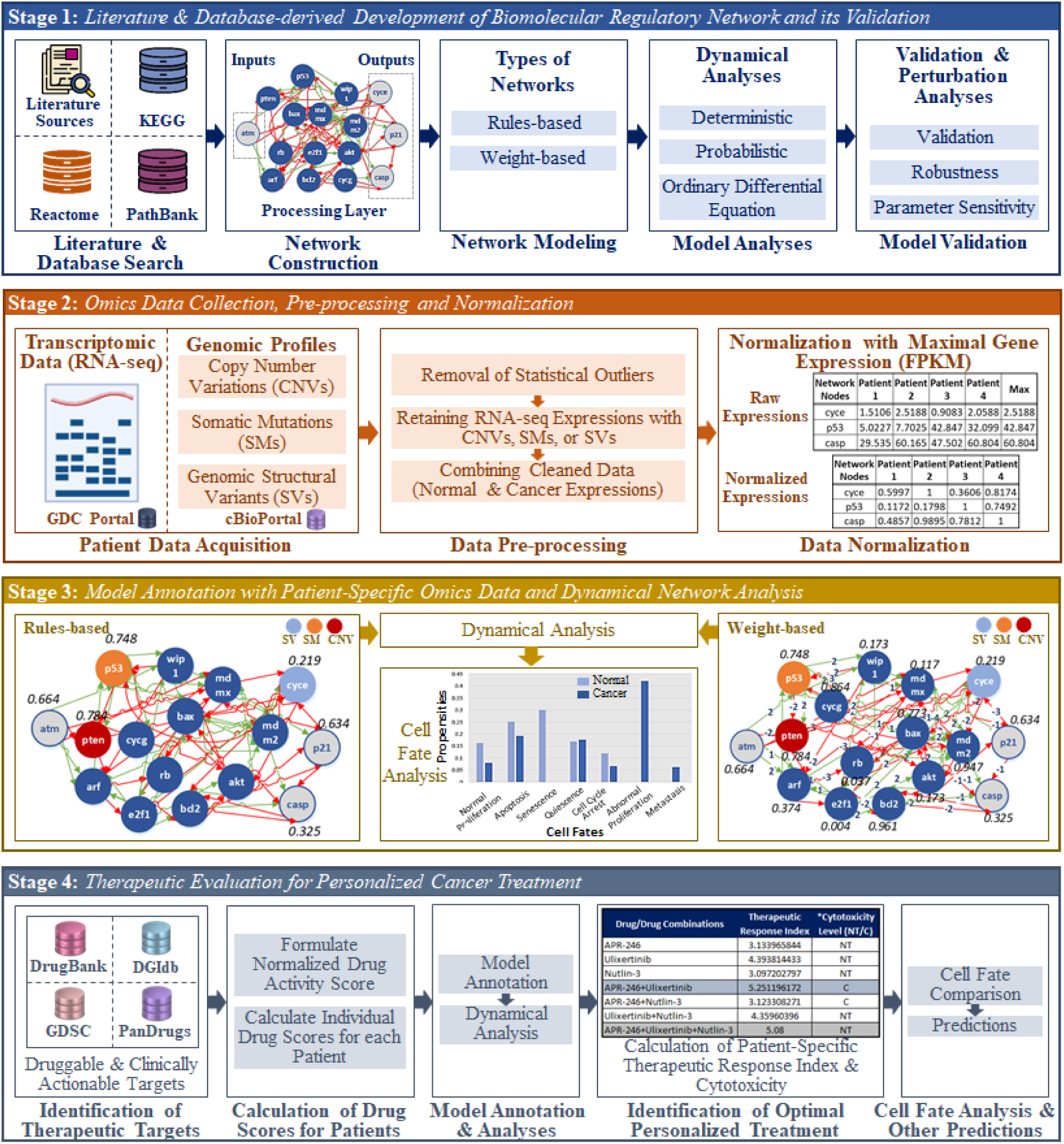
Framework of *CanSeer* for Personalized Cancer Therapeutics. The salient stages of *CanSeer* includes network construction, omics data pre-processing and normalization, model annotation, and therapeutic evaluation. Stage-1 includes the abstraction of biomolecular regulations from literature and databases to construct a network. The constructed network is translated into the rules-based or weight-based model for onwards dynamical (deterministic, probabilistic, or ordinary differential equations) analysis. The model output is validated against literature under normal conditions. Next, the model is tested by perturbing the input conditions for robustness and parameter sensitivity analyses; Stage-2 involves the patient-specific genomic and transcriptomic data acquisition, pre-processing and normalization; Stage-3 undertakes the annotation of rules-based and weight-based network models with patient-specific omics data followed by dynamical analysis and cell fates comparison, and Stage-4 includes the identification of druggable and clinically actionable targets followed by calculating the drug scores for patients. Next, the patient-centric model is annotated with the individualized drug scores followed by the dynamical analysis. The resulting cell fate propensities are further utilized to calculate patient-specific therapeutic response index (TRI) and cytotoxicity to identify the optimal individualized treatment. Lastly, the cell fate outcome of optimal treatment is compared with its normal and cancer counterparts, and predictions are made.

### A. Stage 1 - Literature and Database-derived Development of Biomolecular Regulatory Network and its Validation

To develop cancer-type specific network architectures, published literature as well as online databases including Kyoto Encyclopedia of Genes and Genomes (KEGG) [77], PathBank [78], Pathway Interaction Database (PID) [79], [80], and Reactome [81] are mined for abstracting biomolecular regulations. The topology of resulting networks include input, output, and processing nodes along with their interactions. The input nodes in the network cue the downstream processing nodes which then crosstalk, and signal output nodes. To analyze the model, the network topology is then translated into the rules-based [82]–[84] and weight-based [85], [86] network models. For the dynamical analysis of rules-based or weight-based models, normal ambient conditions are abstracted from the literature, and are assigned to the input nodes. Next, temporal dynamics of the network are performed using deterministic [61], [88], [89], probabilistic [98], [99], or ordinary differential equation [100] modalities under influence of input conditions. Results from dynamical analyses provide output node propensities which are then used to program cell fate outcomes such as quiescence, proliferation, cell cycle arrest, apoptosis, etc. To validate the resulting cell fate outcomes, the trend of cell fate propensities is tallied with the published literature. To further evaluate the biological plausibility and sensitivity of the validated model, random and systematic perturbations are introduced into the model by varying input conditions termed as robustness [101], and parameter sensitivity [102], [103] analyses, respectively. Here again, the resulting cell fate outcomes are matched, and validated against the published literature.

### B. Stage 2 - Omics Data Collection, Pre-processing, and Normalization

In *CanSeer’s* second stage, patient-specific omics data is normalized for annotating the network models. For that, patient-specific transcriptomic (RNA-seq based gene expressions) and genomic (copy number variations (CNVs), somatic mutations (SMs), and genomic structural variants (SVs)) profiles are acquired from The Cancer Genome Atlas (TCGA) PanCancer Atlas [90], [91] studies using Genomic Data Commons (GDC) [92], and cBioPortal [93], [94], respectively. Note that, patient information can be employed from other programs and projects as well such as TARGET (Therapeutically Applicable Research to Generate Effective Treatments) program and GENIE-MSK (Genomics, Evidence, Neoplasia, Information and Exchange at Memorial Sloan Kettering Cancer Center) project which are available at GDC and cBioPortal. In addition to the transcriptomic profiles (RNA-seq based gene expressions), patients’ clinical information and sample sheet are obtained from the TCGA program using GDC. Patient dataset matching is ensured by comparing sample IDs between GDC and cBioPortal. Based on the availability of RNA-seq based gene expression profiles in TCGA, the patient samples are selected and categorized into three cases: *Case 1* – *paired samples* which comprise normal (N) and cancer (C) gene expression data from the same patient, *Case 2* – *unpaired samples* which contain N and C gene expression data from different patients, and *Case 3* – *cancer samples only* that consists of gene expression data from cancer patients. The detailed workflow of each case is provided below.

#### Case 1: Paired Normal and Cancer Samples

Case 1 provides for the situation when data from both normal and cancer samples are available from the same patient(s). It employs RNA-seq based gene expression data for normal and cancer samples stored in FPKM (Fragments Per Kilobase of transcript per Million mapped reads) format. Next, the downloaded gene expressions and associated Ensembl Gene IDs are mapped onto the gene symbols using BioTools.fr [104] toolbox. Then, the gene symbols list is filtered based on the nodes (biomolecules) present in the network model. Aliases for biomolecules that do not tally with the list of gene symbols are obtained from GeneCards [105] or HUGO Gene Nomenclature Committee (HGNC) [106]. Onwards, the network nodes list is converted into the gene symbols to enable fetching of respective RNA-seq based gene expressions.

As a result, patient’s gene expression data is then mapped with the network’s gene symbols for both normal and cancer paired samples, separately. In the case of normal samples, outliers from RNA-seq based gene expression data are removed using statistical outlier removal methods i.e., 3 x MAD (Median Absolute Deviation)[107] or IQR (interquartile range) [108]. While, for cancer samples, the method detects outliers using MAD or IQR followed by splitting the gene expressions into outliers (COL), and retained (CR) gene expression data. For each cancer sample, information on CNVs, SMs, or SVs of aberrant gene expressions is obtained. If the highly altered gene expression is due to CNVs, SMs, or SVs, then the gene expression is retained. Otherwise, the alteration is considered a statistical outlier and removed. The transcriptomic data from normal and cancer samples after the removal of outliers is combined and normalized using the maximal gene expression. Figure 2 illustrates the workflow for pre-processing and normalization of patient’s omics data.

**Figure 2.**
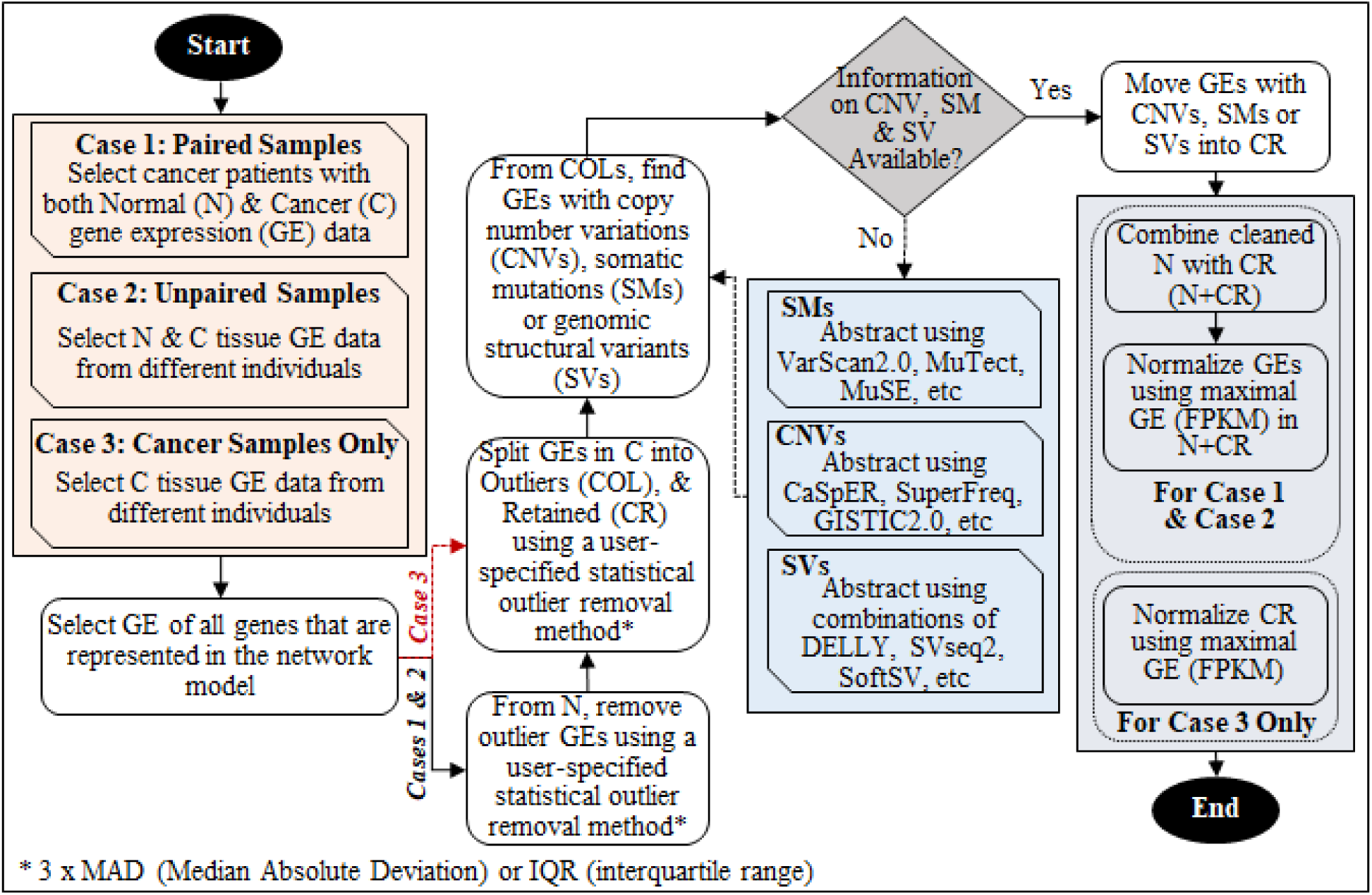
Workflow for Omics Data Pre-processing and RNA-seq based Gene Expression Normalization. The three salient cases in *CanSeer* include (i) Case - 1: paired normal and cancer samples, (ii) Case - 2: unpaired normal and cancer samples, or (iii) Case - 3: cancer samples only. Next, RNA-seq based gene expressions are extracted for all network nodes (biomolecules). Statistical outliers are removed from RNA-seq gene expressions using 3 x MAD (Median Absolute Deviation)[107] or IQR (interquartile range) in normal samples, whereas the expressions are split into Outlier (COL) and retained (CR) in cancer samples. For each cancer sample, the highly altered gene expression due to CNVs, SMs, or genomic SVs, is retained, and remaining outliers are removed. The normal and cancer samples after removal of outliers in Case 1 and 2 are combined and normalized between 0 and 1 using maximal gene expression. In Case 3, cancer samples only are normalized with the highest gene expression after the removal of outliers. In case of missing information on CNVs, SMs, or SVs, various algorithms are referred for abstracting CNVs, SMs, and SVs from whole-exome sequencing.

In case that CNVs, SMs, and SVs are missing for a particular patient, *CanSeer* integrates this information from whole-exome sequence data. For that, algorithms such as VarScan2 [109], MuTect [110], and MuSE [111] can be employed for abstracting SMs. For CNVs, CaSpER [112], SuperFreq [113], and GISTIC2.0 [114] algorithms can be used whereas, a combination of DELLY [115], SVseq2 [116], SoftSV [117], etc can be utilized for obtaining SVs.

#### Case 2: Unpaired Normal and Cancer Samples

In case of availability of normal and cancer expression data for different patients in a particular cancer type and project, Case – 2 is employed. The data pre-processing and normalization strategy in *CanSeer* requires samples containing normal and cancer expression data from different patients while the rest of the process remains the same as in Case 1.

#### Case 3: Cancer Samples Only

In the last case, wherein the availability of RNA-seq based gene expression quantification data is limited to cancer samples only, Case – 3 is employed. Data pre-processing and normalization in *CanSeer* limits the strategy employed for Cases 1 and 2 to cancer samples only.

The sample cases are provided in Supplementary Data 1.

### C. Stage 3 - Model Annotation with Patient-Specific Omics Data and Dynamical Analysis

At the third stage, *CanSeer* employs the normalized RNA-seq based gene expressions of patients along with their CNVs, SMs, and SVs, and incorporates it into the Boolean logic model for personalizing both rules-based and weight-based biomolecular network models. A step-by-step description of model annotation with normalized omics data in case of paired (Case 1), unpaired (Case 2) and cancer samples only (Case 3) is given below.

#### Case 1: Paired Normal and Cancer Samples

In Case 1, patient-specific normalized RNA-seq based gene expressions from paired normal (N) and retained cancer expressions (CR) are used to annotate the “Normal (N)” and “Cancer (C)” network models, respectively. The normalized gene expressions are assigned to the input nodes in rules-based networks while in the case of weight-based networks all network nodes are set with normalized gene expression data. For weight-based networks, basal values are computed for all network nodes in the light of normalized gene expressions of patients. Next, networks are annotated with their genomic alterations including CNVs, SMs, and SVs to develop models of personalized cancers. In the case of rules-based networks, CNVs, SMs, and SVs are incorporated in the model by setting the corresponding nodes to normalized gene expressions such that the node activity is maintained at each time step of dynamical analysis. For weight-based networks, CNVs, SMs, and SVs are integrated into the model by fixing the nodes’ activity with the respective normalized gene expressions at each time-step. Dynamical analysis [87], [118] is carried out for each personalized normal and cancer network model followed by comparing the average node activity of output nodes with the patient-specific normalized gene expressions of respective output genes. A “match” is declared if the average node activity of output nodes is comparable to the normalized gene expressions of corresponding genes. If the result does not compare, then the network is iteratively fine-tuned for making the model representative of the patient. The approach for tuning the network model includes modifying the logical rules and changing the edge weights in rules-based and weight-based networks, respectively. Cell fates from the tuned personalized models are compared for normal and cancer case.

#### Case 2: Unpaired Normal and Cancer Samples

For Case 2, *CanSeer* computes the median gene expression of each gene in normalized N from N+CR, which is used to annotate the network model. Normalized median gene expressions are integrated into the network model to represent a normal network. The normalized median gene expressions are assigned to the input nodes in the rules-based networks. On the other hand, for weight-based networks, all the network nodes are annotated with normalized gene expression data. Basal values are computed in the light of normalized median gene expressions in the weight-based networks. Next, dynamical analysis is performed for the normal network, and the average node activity of output nodes is compared with the normalized median gene expressions of respective genes. Next, the network is annotated with the patient’s cancer gene expressions along with CNVs, SMs, and SVs to model individual tumors. The method of integrating patient-specific transcriptomic (normalized RNA-seq based gene expressions) and genomic (CNVs, SMs, and SVs) data is similar to the personalization of cancer models in Case 1 (see above). The dynamical analysis is performed for personalized cancer network models and the average node activity of output nodes is tallied with the patient-specific normalized cancer gene expressions of corresponding genes. The network personalization step is completed if the average node activity and normalized gene expressions of output nodes are comparable. Otherwise, the network is fine-tuned by modifying the rules and edge weights in rules-based and weight-based networks, respectively. Lastly, the comparison is made between cell fate outcomes from DA of median assigned normal and patient-specific cancer models.

#### Case 3: Cancer Samples Only

*CanSeer* utilizes the normalized patient-specific cancer gene expressions together with the CNVs, SMs, and SVs for the personalization of network model. The integration of omics data into the network model is carried out as described in Case 1. The personalized cancer model is validated by comparing the average node activity of output nodes with the patient’s gene expressions for corresponding genes in cancer samples. Network fine-tuning is performed as described for Cases 1 and 2. A schematic representation of model annotation with omics data to customize the network models to patient-centric models is shown in Figure 3.

**Figure 3.**
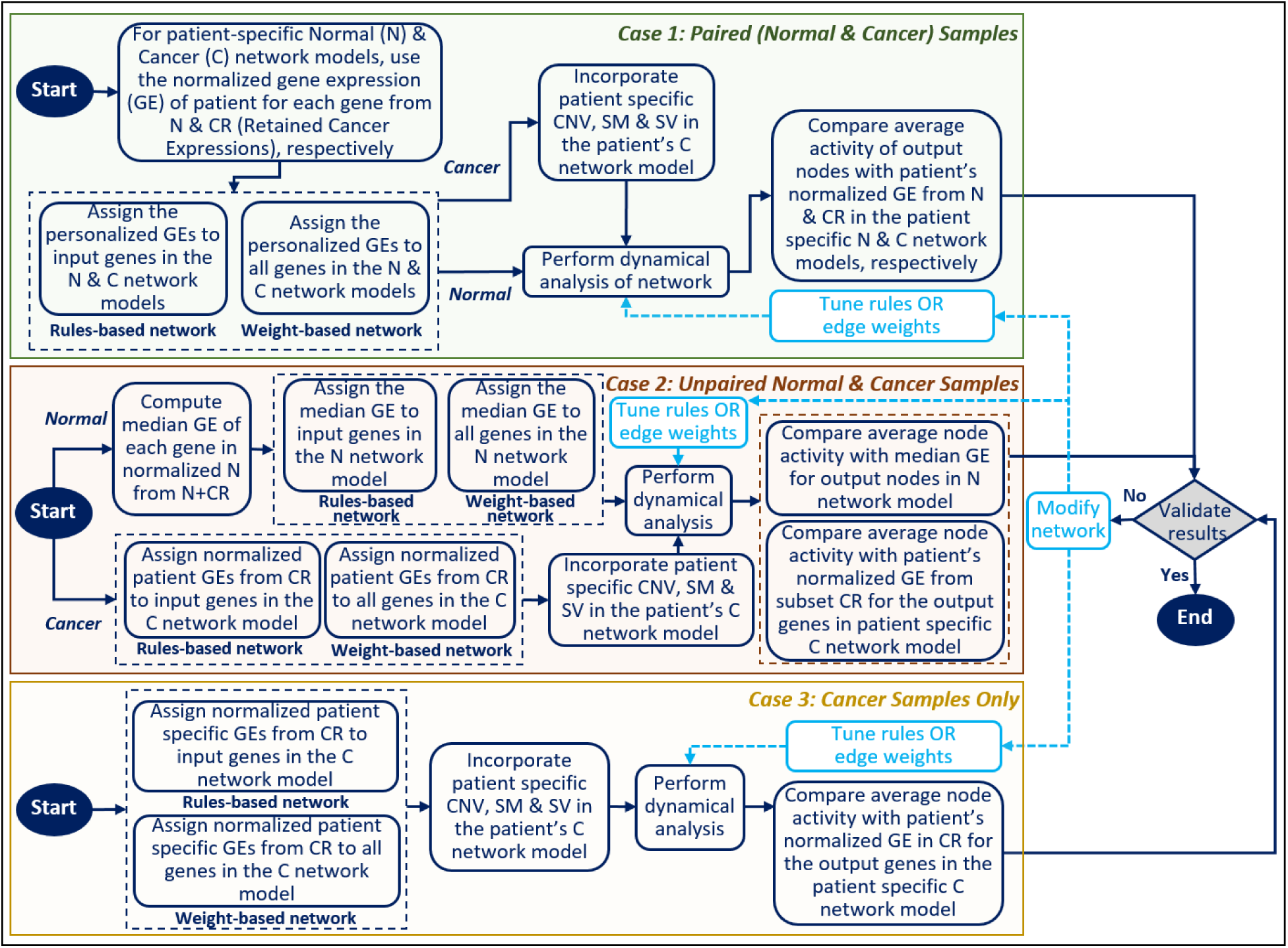
Schematic Representation of Integrating Omics Data into Network Models. In Case 1, patient-specific normalized gene expressions are utilized to annotate the network for making it a patient-centric Normal (N) and Cancer (C) model. Patient-associated Copy Number Variations (CNVs), Somatic Mutations (SMs), and Genomic Structural Variations (SVs) are incorporated in the C model. Dynamical analysis of personalized N and C models is performed and the average node activity of output nodes is compared with the patient-specific normalized gene expression of respective genes. In Case 2, the median gene expression of each gene in normalized N from N+CR (normal and retained cancer expressions) is computed to annotate the network model. The normalized median gene expressions are integrated into the network model representing the N network, whereas patient-specific expressions are assigned to the genes for constructing the individualized C models. Moreover, patients’ CNVs, SMs, and SVs are incorporated in the C model. Then, dynamical analysis of N and personalized C model is performed. The average node activities of output nodes are then tallied with the median and patient-specific expression of respective genes in the N and C models, respectively. In Case 3, the patient’s cancer gene expressions are assigned along with CNVs, SMs, and SVs data incorporation into the model. Subsequently, dynamical analysis is carried out following output nodes validation. In all three cases, the average node activity of output nodes is matched with the respective normalized gene expression. A match is declared if the expression values are comparable, otherwise, the network is iteratively fine-tuned.

### D. Stage 4 - Therapeutic Evaluation for Personalized Cancer Treatment

The last stage of *CanSeer* involves the personalization of cancer therapeutics. For that, personalized *in silico* cancer models are screened under the effect of different drugs and their combinations. Drugs targeting each node (biomolecule) in the network are obtained from published literature, and drug-target databases such as PanDrugs [68], DrugBank [96], and The Drug Gene Interaction Database (DGIdb) [70]–[72]. For enabling the clinical translation of the resultant models, only targetable nodes (either druggable or clinically actionable) are selected along with the corresponding drugs. To systematically prioritize druggability and repurposing opportunities, cancer driver genes in the selected TCGA project are acquired from OncoVar [120], and IntOGen [121]. The drugs identified to target druggable or clinically actionable nodes are then queried in Genomics of Drug Sensitivity in Cancer (GDSC2) [97] to obtain an inhibitory concentration of drugs (IC_50_). The IC_50_ values of candidate drugs are utilized to formulate the potential drug activity. To quantify the potential activity of each drug for a specific tissue type, drug activity scores (DAS) are computed by taking the inversed log IC_50_ value of the drug, as given in the equation below.

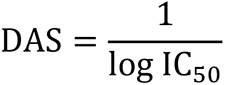

Drug activity scores are then normalized (NDAS) using the highest drug activity score, as shown in the equation below.

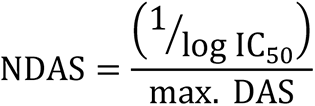

NDAS less than 0 and greater than 1 are approximated to 0 and 1, respectively. Next, the quantified patient-specific drug response termed as “drug score for patient (DS)” is calculated. For that, the maximal efficacy gain induced by a drug (MEGID) using the normalized RNA-seq based gene expression of a patient in cancer (NRGEPC) for the case of a tumor suppressor is computed as follows.

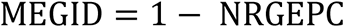

On the other hand, for drug targeting of an oncogene or proto-oncogene, MEGID is set as:

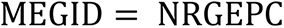

DS is then defined for tumor suppressors as:

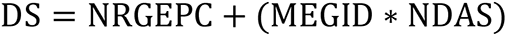

For the case of oncogene and proto-oncogenes:

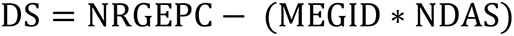

Next, patient-specific drug scores are used to annotate nodes in the personalized network models and the network models are re-analyzed. The cell fate outcomes from dynamical analysis of network models are further employed to calculate the patient-specific therapeutic response index (TRI) against each drug and drug combination. To calculate TRI, the ratio of anti-cancer cell fate propensities (excluding normal proliferation) to the cancer promoting cell fate propensities is taken. Next, the cytotoxicity level of each drug and drug combinations for individual patient is evaluated. For that, 50% of the propensity of normal proliferating cells under cancerous conditions is used as a “cytotoxicity threshold”. The cell fate propensity of normal proliferation under therapy is compared with the measured cytotoxicity threshold for each patient. As a result, the drug or drug combination is declared “non-toxic” for a patient if the propensity of the normal proliferating cell is greater than the computed threshold. Conversely, if the propensity of normal proliferation is less than the threshold, the drug or drug combination is considered “cytotoxic”. *CanSeer* classifies drugs or drug combinations with the highest TRI and “non-toxic” status against cytotoxicity measure, as an optimal personalized treatment option. Lastly, the cell fates under patient-specific optimal treatment conditions are compared with the corresponding normal and cancer model cell fate outcomes followed by interpretations.

### E. Exemplars Demonstrating the Development of Clinically Translational Personalized Cancer Therapeutics using *CanSeer*

To exemplify the employment of “*CanSeer*”, we have constructed three case studies aimed at developing personalized therapeutics for (i) lung squamous cell carcinoma, (ii) breast invasive carcinoma, and (iii) ovarian serous cystadenocarcinoma. As a first step, towards obtaining a biomolecular signaling network for the aforementioned case studies, a literature survey was initiated for a large-scale human signaling network. Consequent to that, a network comprising of 197 nodes including 13 inputs, 8 outputs, and 176 processing nodes along with their interactions [59] was selected for onward employment in the case studies. The network’s 13 input nodes represent different external stimuli and propagate incoming signals to the 176 downstream processing nodes which cue the 8 output nodes. The adopted network was then translated into a rules-based model for dynamical analysis. To determine the dynamical behavior of the reconstructed model, Deterministic Analysis (DA) [61], [87]–[89], [118] was performed under the normal conditions which were assigned to the input nodes (Supplementary Table 1). To program cell fates using output node values obtained from DA, each output node was associated with representative cellular functions, such as normal proliferation, abnormal proliferation, metastasis and quiescence, in light of literature (Supplementary Table 2) [59]. These cell fates and their propensities were then reproduced (within ±10%) using DA in Theatre for *in silico* Systems Oncology (TISON) [87] (Supplementary Figure 1 and Supplementary Data 2). Next, cell fates involved in oncogenesis such as apoptosis, senescence, and cell cycle arrest were also programmed (Supplementary Table 3) and mapped by updating the network rules (Supplementary Information A). Outcomes from DA of the updated network, under normal input conditions, corroborated with the published literature [59] (Supplementary Figure 2 and Supplementary Data 3). Next, to evaluate robustness of the network, perturbations were introduced as input signals and the network response was scrutinized using DA pipeline [87], [118]. The results exhibited the highest variation in normal proliferation (SEM 0.0025) followed by apoptosis (SEM 0.0021) in the network model (Supplementary Figure 3 and Supplementary Data 4). Further, to inspect the distinct input-output relationship, parameter sensitivity analysis was carried out by screening input nodes individually and in combinations. The outcomes indicated that the adopted network model is sensitive to levels of alphailig, DNAdamage, egf, ilonetnf, tgfb, and wnt stimuli. The associated variations in cell fate outcomes are provided in Supplementary Data 5.

Below, we employ this validated network model to develop three different case studies on lung squamous cell carcinoma, breast invasive carcinoma, and ovarian serous cystadenocarcinoma. The case studies demonstrate the development of *in silico* personalized cancer models for precision therapies and manifest an interpretable method to uncover regulatory mechanisms involved in treatment resistance using multi-omics data.

#### Case Study 1: Employing *CanSeer* to Evaluate Personalized Therapeutics for Lung Squamous Cell Carcinoma

A cohort of 551 lung cancer samples was selected from TCGA “Lung Squamous Cell Carcinoma (TCGA-LUSC)” project and associated RNA-seq based gene expression data in FPKM format was extracted for both normal and cancer samples (Supplementary Video 1, and Step-by-Step Guide 1). The downloaded gene expressions associated Ensembl Gene IDs were mapped onto the gene symbols using BioTools.fr (Supplementary Data 6). Next, gene symbols were filtered based on the network nodes representing a variety of biomolecules; since some of the network nodes were not gene names, not all nodes tallied with the gene symbols. For such cases, gene aliases were obtained from GeneCards [61] or HUGO Gene Nomenclature Committee (HGNC) [51] (Supplementary Data 7). In addition to RNA-seq based gene expressions, CNVs, SMs, and SVs of the selected cohort were also obtained from TCGA PanCancer Atlas study using cBioPortal (Supplementary Video 2, and Step-by-Step Guide 2). After data collection, paired normal and cancer samples (Case 1) were obtained from a 551 patient cohort. In the selected cohort, 49 paired samples were retained. Converting the network node list into gene symbols enabled automated fetching of respective RNA-seq based gene expressions for the selected sub-cohort. As a result, patient’s gene expression data was mapped with the network’s gene symbols for both normal and cancer paired samples, separately (Supplementary Data 8 and Supplementary Information B).

In the next step, genomic and transcriptomic data were pre-processed and normalized (Supplementary Data 9 and Supplementary Information B). In the case of normal samples, outliers from gene expression data were removed using IQR. While, for cancer samples, the highly altered gene expressions detected as outliers by IQR method were super-imposed to the patient-specific information on CNVs, SMs, and SVs. The altered expressions due to CNVs, SMs, or SVs were retained while others are removed as outliers. The transcriptomic data from normal and cancer samples after the removal of outliers were combined and normalized using the maximal gene expression (Supplementary Data 10). The normalized gene expressions were then used for network annotation.

To annotate rules-based network models with patient-specific omics data, only those patients were selected for which no gene expression was removed as an outlier for input or output nodes. The removal of outliers reduced the sample size (no. of patients) from 49 to 7 for onward use in the rules-based network model. Then, normalized RNA-seq based gene expressions were assigned to their respective input nodes in rules-based networks. Notably, some input nodes are not directly associated with the gene symbols, hence are not assigned gene expressions directly. Moreover, the input nodes representing biomolecules that contain multiple sub-units were assigned abstracted values computed from downstream nodes’ associated genes. The criteria to assign representative values to network nodes in light of patient’s gene expressions were based on the following: patient mutation (PM), network connectivity (NC), frequency (F), exact match (EM), etc. (Supplementary Table 5 and Supplementary Video 3). In addition to the assignment of normalized patient-specific values to input nodes in normal and cancer models, patient’s CNVs, SMs, and SVs were also incorporated into the cancer model (Supplementary Data 11). The CNVs, SMs, and SVs were integrated by setting the corresponding nodes to normalized gene expressions such that the nodes’ activities remain constant throughout each time step of DA. To ensure that the model is representative of a patient, DA was carried out for each personalized normal and cancer network model with updated cell fates (Supplementary Table 4). The average node activity of output nodes attained from DA was made comparable to the patient-specific normalized gene expressions of respective output genes by iterative fine-tuning (Supplementary Data 11). Next, to compare the patient-centric normal vs cancer model, cell fate outcomes were analyzed. The result generally exhibited a decreasing trend in normal cell fates along with an increase in oncogenic fates in patients’ cancer models (Supplementary Data 11).

In the next step, the personalized *in silico* cancer models were screened under the effect of different drugs and their combinations for evaluating optimal treatment options for each individual patient. For that, an extensive literature, and database search was performed to identify druggable or clinically targetable nodes in the network model along with the targeting drugs (Supplementary Data 12). The drugs reported to target network nodes in clinical settings included APR-246, Nutlin-3, Afatinib, Dinaciclib, Osimertinib, Palbociclib, and Ulixertinib, etc (Supplementary Data 13). Moreover, patient-specific cancer driver genes belonging to the network were identified to facilitate decision-making regarding treatment options. For that, cancer driver genes in the TCGA-LUSC project were acquired from OncoVar [84] and IntOGen [85] (Supplementary Table 6). These cancer drivers were manually searched in the patient-specific genetic alterations to identify patient’s cancer driver genes (Supplementary Data 11).

Next, the short-listed candidate drugs were looked up in GDSC2 to extract IC_50_ values for lineage and site-specific cell line (A549 cell line) of lung squamous cell carcinoma. Utilizing the IC_50_ value of each drug, DAS and NDAS were computed for A549 cell line (Supplementary Data 13). Next, to compute the drug score for each patient, MEGID was calculated using NRGEPC. MEDGID, NRGEPC, and NDAS were used in concert to calculate patient-specific drug scores (DS) (Supplementary Data 11). Next, DS were used to annotate nodes in the personalized network models and the patient models were re-analyzed. The cell fate outcomes from DA were employed to compute patient-specific TRI against each selected drug and drug combinations for individual patients (Supplementary Data 11). To calculate TRI, the cumulative cell fate propensities of apoptosis, quiescence, senescence, and cell cycle arrest were divided by the total propensities of abnormal proliferation and metastasis. The resulting patient-specific TRIs are provided in the respective patient files (Supplementary Data 11). Next, to evaluate the cytotoxicity level of each drug and drug combinations for individual patient, cytotoxicity threshold for each individual was measured and classified as “non-toxic” or “cytotoxic”. The cytotoxicity status of each drug and its combinations for individual patients are provided along with the TRIs in patient files to identify optimal personalized treatment (Supplementary Data 11). Lastly, the cell fates obtained from DA under patient-specific normal, cancer, and optimal treatment conditions were compared, and the predicted outcomes are reported in individual patient files (Supplementary Data 11). Broadly, the cell fate comparisons exhibited an increase in apoptosis, quiescence, and cell cycle arrest along with a decrease in propensities of abnormal proliferation, metastasis, and normal proliferation. However, no general trend was observed in cellular fate “senescence”. Moreover, the results revealed that targeting patient-specific cancer driver genes is an optimal treatment option. Together with this, novel combinations were devised against patient-specific mutations amongst which APR-246+palbociclib, APR-246+afatinib, and APR-246+osimertinib showed optimal therapeutic outcomes.

Towards personalization, a similar approach to rules-based network models was applied for customizing weight-based networks (Supplementary Information C), however, in this case, gene expressions for all network nodes were required. Moreover, only those patients were selected for which no gene expression was removed as an outlier, hence reducing the sample size to just 1 patient (Supplementary Data 14). For weight-based networks, basal values were computed for all network nodes in the light of normalized gene expressions of patients for both normal and cancer models [73]. Next, patient cancer models were further annotated with the patient-specific genetic alterations including CNVs, SMs, and SVs. The remaining steps of the workflow remained the same as in the case of rules-based network models (Supplementary Data 14). The resulting cell fate outcome of the weight-based personalized cancer model was in accordance with its rules-based variant. However, trends of cellular fates in patient-centric normal model showed variation from its rules-based model. Overall, the therapeutic outcome of the same patient through rules-based and weight-based models exhibited that restoration of Tp53 activity together with inhibition of EGFR could be an efficacious combinatorial treatment. Moreover, weight-based analysis further revealed a novel efficacious combination i.e. APR-246+Afatinib+dinaciclib.

#### Case Study 2: Developing Personalized Therapeutics for Breast Invasive Carcinoma using *CanSeer*

Towards demonstrating the development of personalized therapeutics for breast cancer, a cohort of 1222 samples was selected from TCGA’s “Breast Invasive Carcinoma (TCGA-BRCA)” project. The RNA-seq based gene expression data of these patients were downloaded in FPKM format for the following sample types: “solid tissue normal”, “metastatic” and “primary tumor” (Supplementary Video 4, and Step-by-Step Guide 3). RNA-seq based gene expressions associated Ensembl Gene IDs were converted into the gene symbols before onward processing (Supplementary Data 6). The obtained gene symbols list was short-listed for the network nodes and gene aliases were acquired to match the node names (Supplementary Data 7). Additionally, CNVs, SMs, and SVs of the selected cohort were also attained using cBioPortal (Supplementary Video 5, and Step-by-Step Guide 4).

After gathering all the required omics data, sub-cohorts of 50 normal samples and 70 cancer samples were randomly selected from 1222 samples for unpaired normal and cancer samples (Case 2). Note that, unpaired samples were taken only to demonstrate *CanSeer* for Case 2, however, the samples from the same patients for normal and cancer are available in “TCGA-BRCA” project. The selected network’s gene symbols for both normal and cancer samples, separately (Supplementary Data 15). Next, to normalize the RNA-seq based gene expressions, the normal samples were first pre-processed using IQR, and outliers were removed. While, for cancer samples, the highly altered gene expressions detected as outliers (by IQR method) were removed except for the expressions altered due to CNVs, SMs, or SVs. The transcriptomic data from unpaired normal and cancer samples after the removal of outliers were combined and normalized by the highest expression of that gene in a selected dataset (Supplementary Data 9, 16, and Supplementary Information 2).

Following the normalization of omics data, the median gene expression of each gene in normalized N from N+CR was computed (Supplementary Data 17). Normalized median gene expressions were then assigned to the input nodes of the rules-based network to represent a normal network model. Importantly, some input nodes (biomolecules) can’t be directly linked to gene symbols or contain multiple sub-units. To assign a specific representative value to such nodes, the input values were computed from adjacent downstream network nodes (genes). The criteria designed to compute representative values include the following: exact match (EM), patient mutation (PM), matching the downstream node’s average activity with the normalized gene expression by varying the input value from 0 to 1 and selecting that input value at which downstream node’s activity matches its respective normalized gene expression (S_DS), etc. (Supplementary Table 5, Supplementary Video 3). Next, DA was performed for the normal network, and the average node activity of output nodes was compared with the normalized median gene expressions of respective genes (Supplementary Data 17). For developing individualized cancer models, 13 samples left after the removal of outliers, were considered. Among these 13 samples, rules-based network models were developed for only 6 patients. Others were found to have either genetic alterations beyond network architecture or insufficient drug information to target the patient mutations. For developing the rules-based models, the normalized patient-specific cancer expressions and abstracted values were assigned to the input nodes (Supplementary Data 17). In addition, CNVs, SMs, and SVs were also integrated into the patient cancer model (Supplementary Data 17) by setting the corresponding nodes to normalized gene expressions such that the nodes’ activities remained constant throughout DA. Subsequently, DA was executed for each personalized cancer model and the average node activity of output nodes was compared to the corresponding normalized gene expressions of that patient. To make average node activity and normalized gene expression comparable for each output node, the patient models were fine-tuned iteratively. Next, to compare the median assigned normal vs patient-centric cancer model, cell fate outcomes attained from DAs were analyzed. The cell fate comparison showed a mixed trend of variations in oncogenic fates in patients’ cancer models in comparison to the median assigned normal network (Supplementary Data 17). Hence, indicating that median expressions are not reflective of absolute normal expressions of patients.

For the weight-based normal network (Supplementary Information 3), all the network nodes were annotated with normalized median gene expressions and the basal values were computed. Next, DA was performed and the average node activity of output nodes was compared with the normalized median gene expressions of respective genes (Supplementary Data 18). Further, the model was iteratively fine-tuned for ensuring to be the patient’s representative. For developing weight-based individualized cancer models, the method of network annotation with normalized RNA-seq based gene expressions, CNVs, SMs, and SVs is similar to the personalization of cancer models as in Case Study 1. Here, only 1 weight-based personalized cancer model was developed, as only 1 patient was left with no gene expression removed as an outlier (Supplementary Data 18). The cell fate outcomes of the weight-based median assigned normal network revealed that the median gene expressions of normal samples do not depict normal condition as gene expressions vary widely amongst patients, even in normal case. Therefore, the cell fate comparison between median assigned normal model and patient-centric cancer model did not exhibit meaningful trends.

Each personalized *in silico* cancer model was screened under the effect of different drugs and their combinations. In order to find efficacious treatment for each individual patient of breast invasive carcinoma, druggable and clinically targetable nodes in the network model were identified (Supplementary Data 12). Alongside targetable network nodes, the drugs reported to target these nodes were also collected. The selected drugs included Olaparib, KU-55933, Afuresertib, APR-246, Nutlin-3, Afatinib, Osimertinib, Palbociclib, and Carmustine, etc (Supplementary Data 13). Additionally, cancer driver genes in personalized models were identified by mapping cancer drivers of TCGA-BRCA project to patient-specific genetic alterations (Supplementary Table 6, and Supplementary Data 17 and 18). The short-listed candidate drugs for patient driver and passenger mutations were queried in GDSC2 to extract IC_50_ values for BT-474 cell line. Employing the IC_50_ value of each drug, DAS and NDAS were computed for BT-474 cell line (Supplementary Data 13). Next, to compute individual DS, the formula given in Stage 4 were employed (Supplementary Data 17 and 18) followed by personalized model annotation and analyses. The cell fate outcomes from DAs were used to compute TRI against each selected drug and drug combinations for each patient (Supplementary Data 17 and 18). For computing TRIs, the ratio of cumulative cell fate propensities of anti-cancer fates (apoptosis, quiescence, senescence, and cell cycle arrest) to total propensities of cancer promoting fates (abnormal proliferation and metastasis) was taken. The resulting patient-specific TRIs are provided in their respective patient files (Supplementary Data 17 and 18). Next, to categorize the drugs and drugs’ combinations as “non-toxic” or “cytotoxic” for individual patient, cytotoxicity threshold of drugs and drugs’ combinations was measured for each patient. The cytotoxicity status of each drug and its combinations for individual patients together with TRIs reveal efficacious tailored treatment options (Supplementary Data 17 and 18). Lastly, for comparison of cell fates between normal, cancer and predicted efficacious treatment, only patient-specific cancer and therapeutic cell fates were taken into account. The cell fate outcomes of median assigned normal network were excluded. As a result, no general increasing or decreasing trend was observed in cellular fates. In general, treatment resistance was observed against Afuresertib (AKT3 inhibitor) in BRCA patients (Supplementary Data 17 and 18). Likewise, patients with PTEN mutation showed no benefit upon treatment with KU-55933 (an ATM inhibitor indirectly targeting PTEN). These results are in line with DGIdb that reports ATM and PTEN genes to be “drug resistant” [70]–[72]. Despite the differences in cellular fates, the therapeutic outcomes (i.e., resistance against Afuresertib) was found to be consistent.

#### Case Study 3: Identifying optimal individualized treatment for Ovarian Serous Cystadenocarcinoma

In case study 3, a cohort of 379 ovarian cancer patients was selected from TCGA “Ovarian Serous Cystadenocarcinoma (TCGA-OV)” project. The project contained only cancer samples (Case 3) and the associated RNA-seq based gene expression files (FPKM format) were downloaded from GDC portal (Supplementary Video 6, and Step-by-Step Guide 5). The extracted files contain RNA-seq based gene expressions against Ensembl Gene IDs. To map the gene expressions to gene symbols, Ensembl Gene IDs were converted to gene symbols using BioTools.fr (Supplementary Data 6). The obtained gene symbols list was then short-listed for network nodes and gene aliases were acquired to complement the node names (Supplementary Data 7). Besides RNA-seq based gene expressions, CNVs, SMs, and SVs of the selected cohort were also acquired from cBioPortal (Supplementary Video 7, and Step-by-Step Guide 6).

Next, a sub-cohort of 10 cancer patients was randomly selected for mapping network nodes with gene symbols, which enabled automated fetching of RNA-seq based gene expressions (Supplementary Data 19). After obtaining the patients’ gene expressions, the statistical outliers were detected. To remove outliers from cancer samples, patient-specific information on CNVs, SMs, and SVs was over-laid with the RNA-seq based gene expressions. The highly altered gene expressions detected as statistical outliers by IQR method were retained if due to CNVs, SMs, and SVs, otherwise they are removed. Then, RNA-seq based gene expressions were normalized by the highest gene expression against all samples (Supplementary Data 20). The normalized RNA-seq based gene expressions were then used for network annotation.

To annotate rules-based models with patient-specific omics data, only those patients were selected for which no gene expression was removed as an outlier for both input or output nodes. While, for weight-based models, only those samples were considered for which normalized gene expressions were retained for all network nodes. The removal of outliers reduced the sample size (no. of patients) from 10 to 4 for onward use in rules-based models and further to 1 for weight-based model. Then, normalized gene expressions were assigned to their respective input nodes in rules-based models and to all network nodes in weight-based model. The input nodes representing biomolecules with multiple sub-units or not directly linked to gene symbols, were assigned abstracted values (Supplementary Table 5, Supplementary Video 3). In case of rules-based models, patient’s CNVs, SMs, and SVs were incorporated in the network in addition to assigning values to input nodes (Supplementary Data 21). In case of weight-based model, basal values were computed in light of normalized gene expressions of patient and CNVs, SMs, and SVs were integrated into the model. To ensure that the model is representative of a patient, DA was carried out for each rules-based and weight-based personalized models. The average node activity of output nodes obtained from DA were iteratively tuned to the corresponding patient-specific normalized gene expressions (Supplementary Data 21 and 22).

In the next step, the personalized *in silico* cancer models were screened for various drugs and their combinations. To find optimal treatment option for individual ovarian cancer patient, the druggable and clinically actionable targets accompanied by the drugs were selected from literature and databases (Supplementary Data 12). The selected drugs included APR-246, Nutlin-3, Dinaciclib, Ipatasertib, and Ulixertinib, etc (Supplementary Data 13). Next, to quantify the potential activity of these drugs (DAS) for ovarian serous cystadenocarcinoma, IC_50_ values for well-established ovarian cancer cell line (OVCAR-3 cell line) were acquired from GDSC2. The computed DAS together with its normalization (NDAS) for OVCAR-3 cell line are provided in (Supplementary Data 13). Next, individualized DS was computed using the mathematical relationship formulated in Stage 4 (Supplementary Data 21 and 22). With the computed DS, the personalized network models were again annotated, and re-analyzed. The cell fate outcomes from DA were employed to compute TRI against each selected drug and drug combinations for individual patients along with cytotoxicity status. The treatment option with highest TRI and “non-toxic” status was considered best among all drugs and drugs’ combinations tested (Supplementary Data 21 and 22). Further, the cell fates of predicted best tailored treatment were compared with the cell fates of cancer counterparts. Consequently, no general trend was observed in cell fate outcomes. Moreover, for making better treatment decision, cancer driver genes of patients were identified by mapping cancer drivers of TCGA-OV project to patient-specific genetic alterations (Supplementary Table 6, and Supplementary Data 21 and 22). Consequently, targeting patient cancer driver genes were found to be more viable. However, patients with genetic alteration of CCNE1 showed resistance against dinaciclib, irrespective of being a cancer driver gene. This result is in accordance with DGIdb that classifies CCNE1 gene to be “drug resistant” [70]–[72]. Lastly, the case study also revealed the combinatorial efficacy of APR-246+Carmustine, and APR-246+Dinaciclib for treatment of ovarian cancer.

## DISCUSSION

Computational analysis of biomolecular regulatory networks in cancer has become an indispensable tool for elucidating the mechanistic underpinnings of tumorigenesis and progression [56]–[58], [60]. Dynamical simulations of such models have helped classify patients into clinical sub-groups [55], [64], [122] as well as predict survival [52], [53], [55], [122]. Further, network modeling has also been used to evaluate efficacious drug targets by perturbing the network nodes - as singletons as well as in combinations [63], [74]. However, the lack of incorporation of information on druggable, and clinically actionable targets in the perturbation-based therapeutic screening, impedes the translation of *in silico* network models in personalized cancer therapeutics. Towards enhancing the clinical relevance of computational models, recent studies have incorporated RNA-seq based gene expressions, copy number variations (CNVs), and somatic mutations (SMs) in the network models to map patients’ molecular profiles. However, qualitative translation remains inadequate in capturing a patient’s molecular profile, which then further hinders the development of clinically-translational synergistic combinatorial treatments. Additionally, the employment of online drug databases which also contain clinically disapproved drugs can lead to off-label *in silico* drug targeting. As a consequence, the assistance offered by *in silico* studies towards personalized cancer care remains rudimentary in the context of clinical adoption [50]. Moreover, the simulation of drug effects using drug scores coupled with quantitative patient’s omics data to assess therapeutic response remains unevaluated. Lastly, integrative modeling frameworks that facilitate computational investigations into optimal treatments, drug resistance mechanisms, and cytotoxicity towards designing personalized therapies remain lacking. Taken together, there is a need for a systematic method to develop *in silico* models that provide comprehensive coverage of patient’s molecular profile, as well as facilitate clinical translation of modeling-based personalized therapeutics.

For addressing this need, we propose “*CanSeer*”, a method to develop *in silico* models for clinical translation of personalized therapeutics. *CanSeer* couples state-of-the-art dynamical analysis of biomolecular network models with patient-specific genomic and transcriptomic data to assess the individualized therapeutic responses to targeted drugs and their combinations. Towards a quantitative evaluation of personalized *in silico* models, *CanSeer* employs RNA-seq based gene expression data to annotate the network models. Moreover, the patient’s CNVs, SMs, and genomic structural variants (SVs) are incorporated to enable the elucidation of molecular signatures. *CanSeer* envisages quantitative personalized models in comparison to recent approaches that encoded CNVs (amplified genes and homozygous deletions), and SMs (oncogenes and tumor suppressors) as 0 or 1. For an integrative investigation into patient tumor, statistical outliers detected in RNA-seq based gene expression data having CNVs, SMs, and SVs are retained. Considering the heterogeneous impact of genetic alterations, network nodes are annotated with normalized RNA-seq based gene expressions such that the node activity remains conserved throughout the analysis for genetically altered genes. For enabling translational pertinence and prioritizing repurposing opportunities, cancer driver genes in patients are identified, and druggable and/or clinically actionable target nodes are selected for therapeutic interventions. As a result, *CanSeer* is capable of eliciting optimal tailored treatment by employing therapeutic screening of druggable and clinically actionable targets, hence enabling clinical translation. The proposed formulation of drug scores further assisted in capturing the effect of drug to drug variability, and facilitated the evaluation of specific drugs on different cell lines. Computing patient-specific drug scores allowed identifying patient responses to targeted drugs and their combinations. Finally, the patient-specific therapeutic response index together with evaluating cytotoxicity helped identify optimal tailored treatment.

*CanSeer* takes Kauffman’s logical modeling approach [64] and employs deterministic analysis pipeline [87], [118], to enable customization and evaluation of large personalized networks. Earlier, Beal *et al.* devised personalized models by integrating CNVs, and SMs as discrete data, and expressions (gene/protein) as continuous data into Boolean models [52]. However, resultant model’s integrative investigation remained limited as CNVs, and SMs remained unmapped with expression data. *CanSeer* addresses this by integrating RNA-seq based gene expressions with CNVs, SMs, and SVs for a consolidative investigation into patient’s tumor. The mapping between gene expression and genetic alterations is based on positive correlation of CNVs, SMs, and SVs with gene expression levels [125]–[131]. In 2022, Montagud *et al.* [132] applied Beal *et al’s* approach on Fumia’s Boolean model [133] to personalize 488 prostate cancer patients, and eight cell lines. Despite having the multiple readouts including proliferation, invasion, migration, metastasis, DNA repair, and apoptosis, the study remained limited to eliciting proliferation and apoptosis in patient-centric models. Further, to identify potential treatments, knock-outs were performed to increase apoptosis, and deplete proliferation. A similar approach was employed by Mahnoor *et al*. [74] to predict efficacious combinatorial treatments for colorectal cancer. However, the discrete perturbations only show the maximal effect, and hence cannot be employed in either wet-lab experiments or clinical trials. In contrast, *CanSeer* takes into account druggable and clinical actionable targets, and investigates the quantitative patient treatment response by incorporating drugs’ IC_50_ values to establish clinical relevancy and applicability. Moreover, in the Mahnoor *et al*’s colorectal cancer study, cell-type specific RNA-seq based gene expressions of fly gut was integrated with patient-specific mutations to identify personalized drug combinations [74].

In contrast, *CanSeer* incorporates a comprehensive molecular signature comprising of patient-specific RNA-seq based gene expressions, CNVs, SMs, and SVs. Towards improving clinical translation of computational precision oncology approaches, Ianevski *et al*. executed machine learning approach to select patient-specific drug combinations with synergy-efficacy-toxicity balance [66]. However, some of the identified combinations were not clinically applicable due to the utilization of public drug databases that contain incomplete information. *CanSeer* employs extensive literature review in addition to drug database utilization to avoid devising inoperable combinations. In addition, *CanSeer* applies computational modeling approach which provides mechanistic insights into patient tumors [67], and resistance mechanism in addition to cytotoxicity evaluation, and optimal tailored treatment identification.

*CanSeer’s* methodology has been exemplified through three different case studies to identify personalized optimal treatment by tailoring targeted drugs as well as their combinations. In case study 1, the restoration of Tp53 activity together with an inhibition of EGFR was found to be an efficacious combinatorial treatment for patients with Tp53 and EGFR cancer driver genes. Novel therapeutic combinations including APR-246+palbociclib, APR-246+afatinib, APR-246+Osimertinib, and APR-246+afatinib+dinaciclib were identified for treatment of lung squamous cell carcinoma. Case study 2 exemplified *CanSeer*’s potential to elucidate drug resistance against targeted drugs and their combinations including KU-55933, afuresertib, ipatasertib, and KU-55933+afuresertib. Case study 3 revealed the combinatorial efficacy of APR-246+Carmustine, and APR-246+Dinaciclib for treating ovarian serous cystadenocarcinoma.

In terms of limitations and future extensions, *CanSeer* can cue the development of a multi-scale modeling pipeline around the framework [134] by integrating regulations from different spatiotemporal scales. Bootstrapping and applying *CanSeer* to a large cohort of patient samples can help elucidate the bigger landscape of association between patient-specific omics data and therapeutic combinations. Furthermore, wet lab validation of resulting novel combinations, and repositioned drugs can enhance the method’s reliability towards clinical acceptability. The framework can be extended further as a web-based application to provide high-throughput performance by leveraging on graphics processing units (GPUs) using CUDA-toolkit to enable faster analyses for predicting personalized treatments. Additionally, novel pathway, and drug databases, accompanied by cancer omics and clinical datasets can be integrated to enhance applicability of *CanSeer*.

## Supporting information

Step by Step Guide

## CONCLUSION

In Conclusion, *CanSeer* is a novel translational precision oncology framework that assists in leveraging clinically actionable information to formulate deployable personalized treatment plans.

## DATA AVAILABILITY STATEMENT

All the data, and its interpretation needed to evaluate the results of the study are available along with the manuscript, supplementary materials, and supplementary data. The supplementary videos are available at https://www.youtube.com/playlist?list=PLaNVq-kFOn0a6HWmZq_KxeSbY6UpnvvKB

## AUTHOR CONTRIBUTIONS

SU supervised the study. SU and RN designed the personalized therapeutics framework. RN carried out the literature review, developed the case studies, undertook the analyses, performed validations, and interpreted the results. BA, MU, ZT and MF implemented the code for data pre-processing, normalization, and deterministic analysis. SU, and RN drafted the manuscript. RH, RA, SK, MN, AF and MS assisted in devising the strategy for the framework. All authors read and approved the final manuscript.

