## Supplementary material for "CanSeer: A Method for Development and Clinical Translation of Personalized Cancer Therapeutics": Captions.docx

**Supplementary Figure 1** – **Cell Fate Propensities of Human Signaling Network.** Cho *et al.*’s rules-based human signaling network was reproduced using TISON, followed by the comparison of cell fate propensities.

**Supplementary Figure 2** – **Cell Fate Propensities of Updated Network.** The comparison of cell fate propensities of updated network with the following under normal conditions: (i) model result reported by Cho *et al.*, and (ii) the deterministic analysis result of Cho *et al* reproduced using TISON.

**Supplementary Figure 3** – **Cell Fate Outcome of Robustness Analysis along with Corresponding Standard Error of Means (SEMs).** The graph shows average cell fate propensities and the corresponding SEMs obtained as a result of robustness analysis.
