## Supplementary material for "CanSeer: A Method for Development and Clinical Translation of Personalized Cancer Therapeutics": Captions.docx

**Supplementary Table 1** – **Normal Input Conditions.** The normal ambient conditions abstracted from literature, and assigned to the input nodes.

**Supplementary Table 2** – **Cell Fate Classification.** The cellular phenotypes and their associated biomarkers for determining cell fates.

**Supplementary Table 3** – **Cell Fate Expansion.** Expansion of cellular phenotypes and their associated biomarkers to map cell cycle arrest, senescence, and apoptosis in addition to normal proliferation, quiescence, abnormal proliferation, and metastasis.

**Supplementary Table 4** – **Cell Fate Classification in Light of Patient's RNA-seq Gene Expression Data.** The cell fate classification designed in light of patient's RNA-seq gene expression data normalized between 0 and 1, wherein each gene has values ranging between 0 and 1 across patients.

**Supplementary Table 5 – Gene Selection Code and Description.** The criteria for selecting genes to assign representative values to network nodes are represented by codes, followed by their description.

**Supplementary Table 6 – Cancer Driver Genes.** The cancer driver genes of “TCGA-LUSC”, “TCGA-BRCA”, and “TCGA-OV” projects.
