## Supplementary figures and images for "CanSeer: A Method for Development and Clinical Translation of Personalized Cancer Therapeutics"

### Supplementary_Figure_1.png

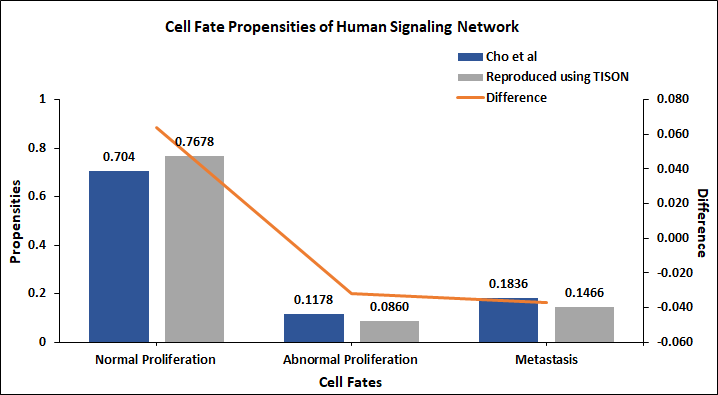

### Supplementary_Figure_2.png

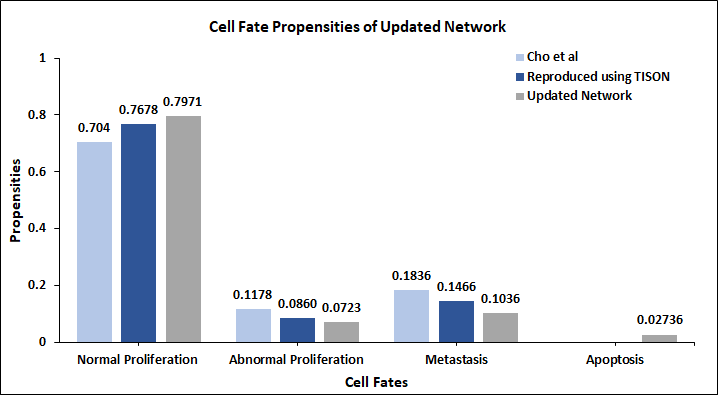

### Supplementary_Figure_3.png

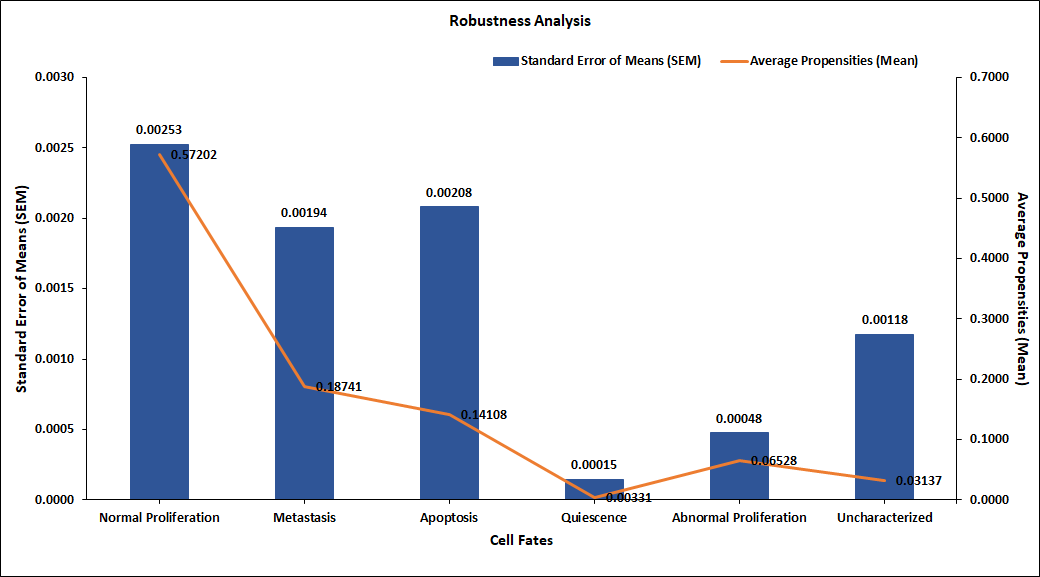
