## Supplementary Information for "CanSeer: A Method for Development and Clinical Translation of Personalized Cancer Therapeutics"

### Table of Contents

**A. Updating Network Rules to Map Cell Fates** ........ 3

**B. Implementation of Genomic and Transcriptomic Data Pre-processing and Normalization** 3

**C. Conversion of Rules-based Network to Weight-based Network** ........ 4

#### A. Updating Network Rules to Map Cell Fates

The rules for “caspthree”, “caspnine”, and “ptwoone” network nodes were updated, which enabled the mapping of following cellular fates: apoptosis, senescence, and cell cycle arrest.

#### B. Implementation of Genomic and Transcriptomic Data Pre-processing and Normalization

The *CanSeer* genomic data pre-processing and RNA-seq based gene expression data normalization algorithm has been implemented in MATLAB 2020b. The process initiates with filtering of the sample sheet obtained from the TCGA program based on the following keywords: ‘normal’, ‘tumor’, ‘primary’ and ‘metastatic’ to include all types of normal and tumor samples, except ‘recurrent tumors’. Next, specific case amongst Cases 1, 2, and 3 as (i) “paired”, (ii) “unpaired”, and (iii) “cancer samples only” is selected. For case 1, a new sample sheet is assembled for both normal and cancer samples that includes patient/sample IDs along with the RNA-seq gene expression file names mapped against normal and cancer samples. For case 2, two separate sample sheets are formed for the normal and cancer case. Each sample sheet contains patient/sample IDs with their corresponding RNA-seq gene expression file names. For case 3, the sample sheet is assembled for cancer samples only. Subsequently, the option of random sample or complete dataset of RNA-seq gene expressions can be opted. Next, the network’s nodes list, comprising of all network nodes, is employed to extract and align RNA-seq gene expressions of patients with the respective genes and patient IDs. In the next step, the outliers are detected in the dataset using MAD (Median Absolute Deviation) or IQR (interquartile range) methods. After detection of statistical outliers, Copy Number Variations (CNVs), Somatic Mutations (SMs) and Genomic Structural Variations (SVs) are selected and processed. From CNVs, deep deletions (-2) and amplification (+2) (based on GISTIC processing) are retained. CNVs having low-confidence values of –1 and +1, obtained using GISTIC processing, are removed. The MATLAB script encodes deep deletions of genes as -2, amplifications as +2, and remaining data as 0. Next, the filtered SMs (for selected patients and network genes) are transformed into logical arrays, where 0 and 1 represent the absence and presence of mutation, respectively. Similarly, patient-specific SVs are also transformed into logical arrays representing the absence (0) and presence (1) of genomic SVs. Lastly, the detected outliers are super-imposed with the CNVs, SMs, and SVs to retain the highly altered RNA-seq gene expressions (CR) resulting from CNVs, SMs, and SVs. The remaining outlier gene expressions are removed after which the normal and cancer samples are combined and normalized between 0 and 1 using the highest gene expression across patients. The normalization strategy for Case 3 is similar to Cases 1 and 2, with the exception of availability of RNA-seq gene expression data for only cancer samples, wherein, the RNA-seq gene expressions are normalized by the maximum gene expression from cancer samples only.

##### C. Conversion of Rules-based Network to Weight-based Network

To begin with weight-based network customization, the rules-based to weight-based network conversion was required. For that, the basal values of all network nodes were initially kept as 1. Subsequently, the edge weights were computed based on the number of adjacent nodes such that the deterministic analysis (DA) result of weight-based network matches with the rules-based DA results. The converted network was then utilized for developing weight-based personalized models.
