## Supplementary material for "CanSeer: A Method for Development and Clinical Translation of Personalized Cancer Therapeutics": Step by Step Guide: Step_by_Step_Guide_2.docx

### Step-by-Step Guidelines for Downloading Patient Genomic Data from cBioPortal

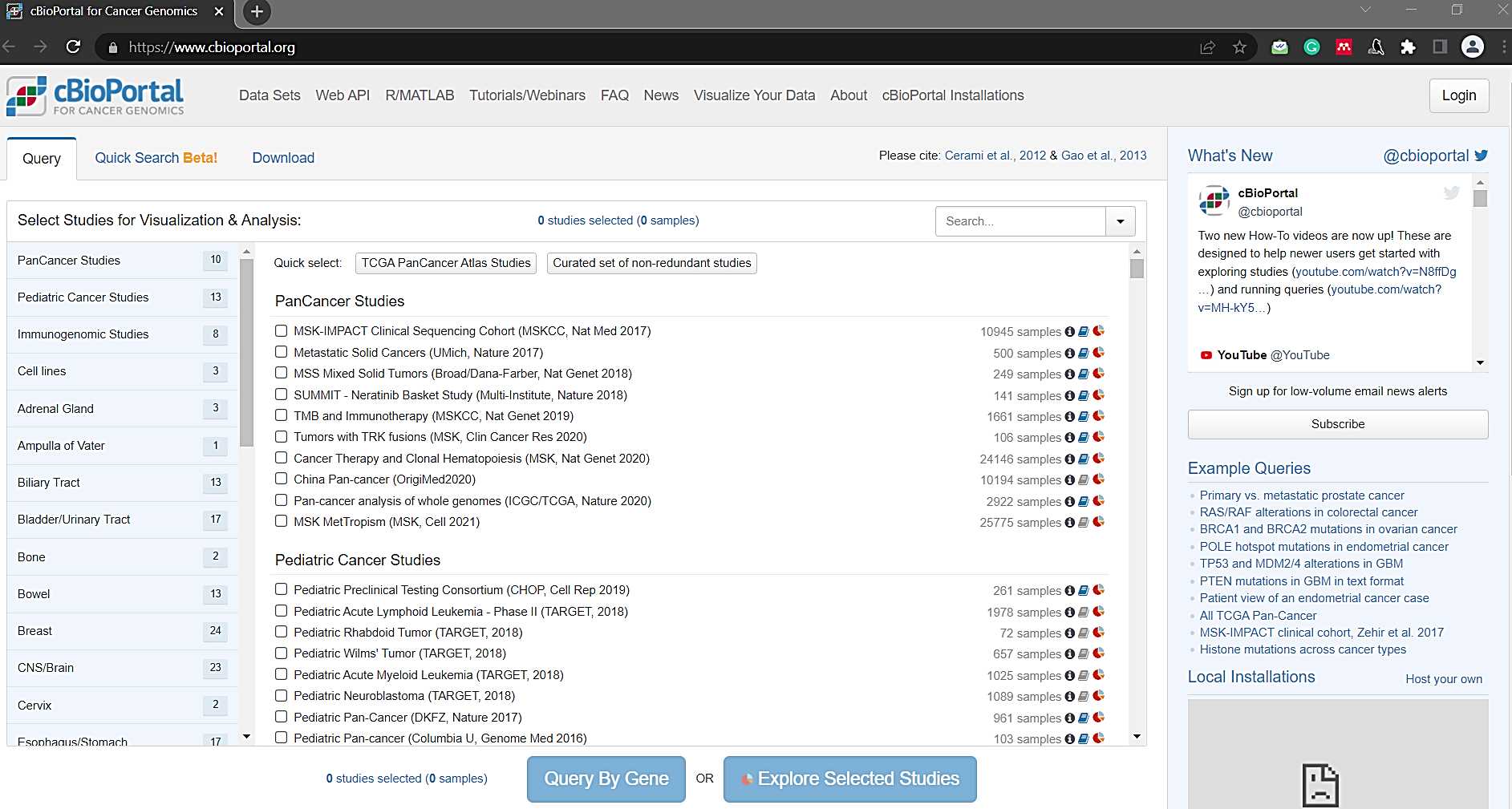

**Step1:** Go to https://www.cbioportal.org/

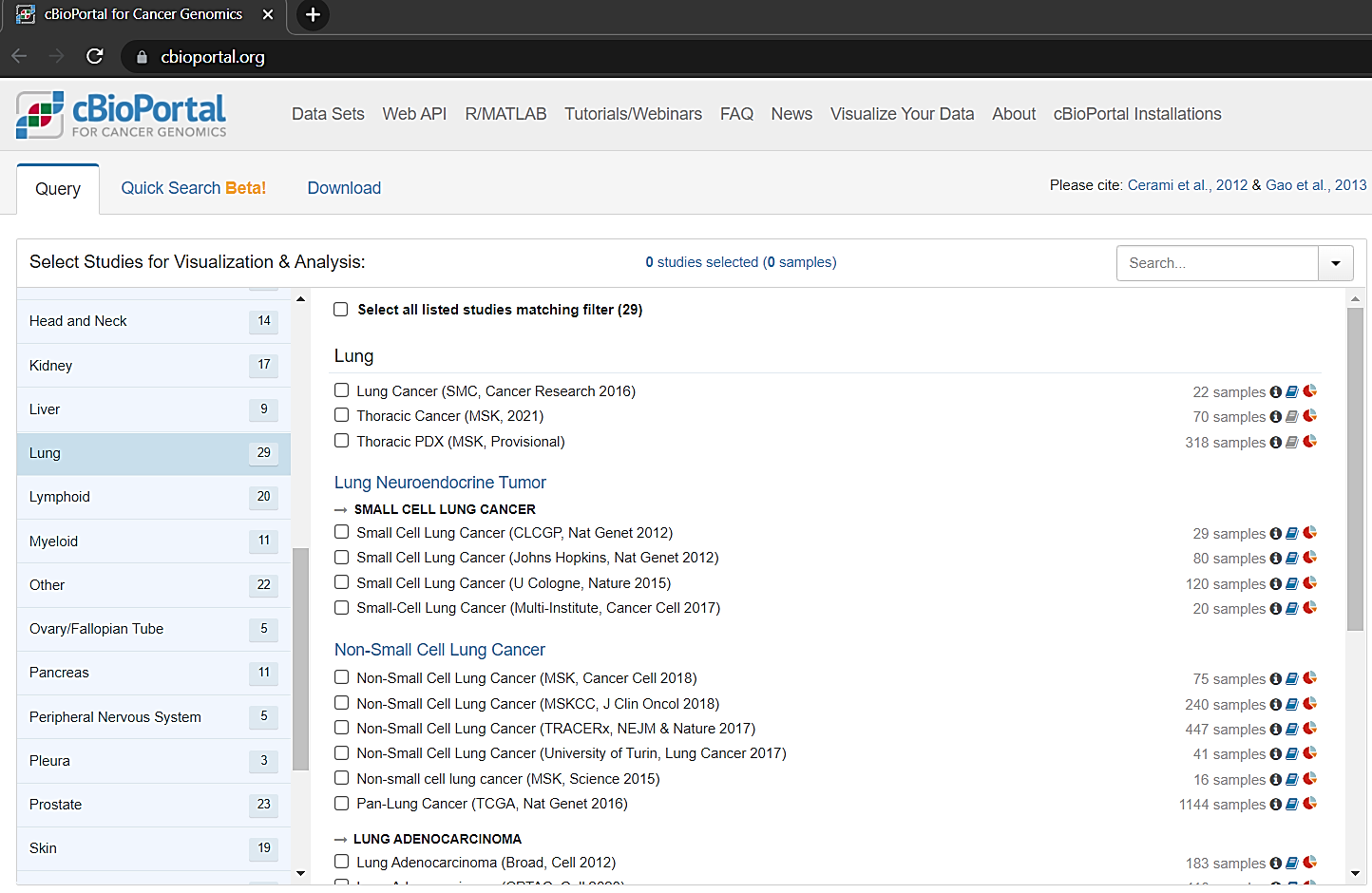

**Step2:** Select the study e.g., Lung

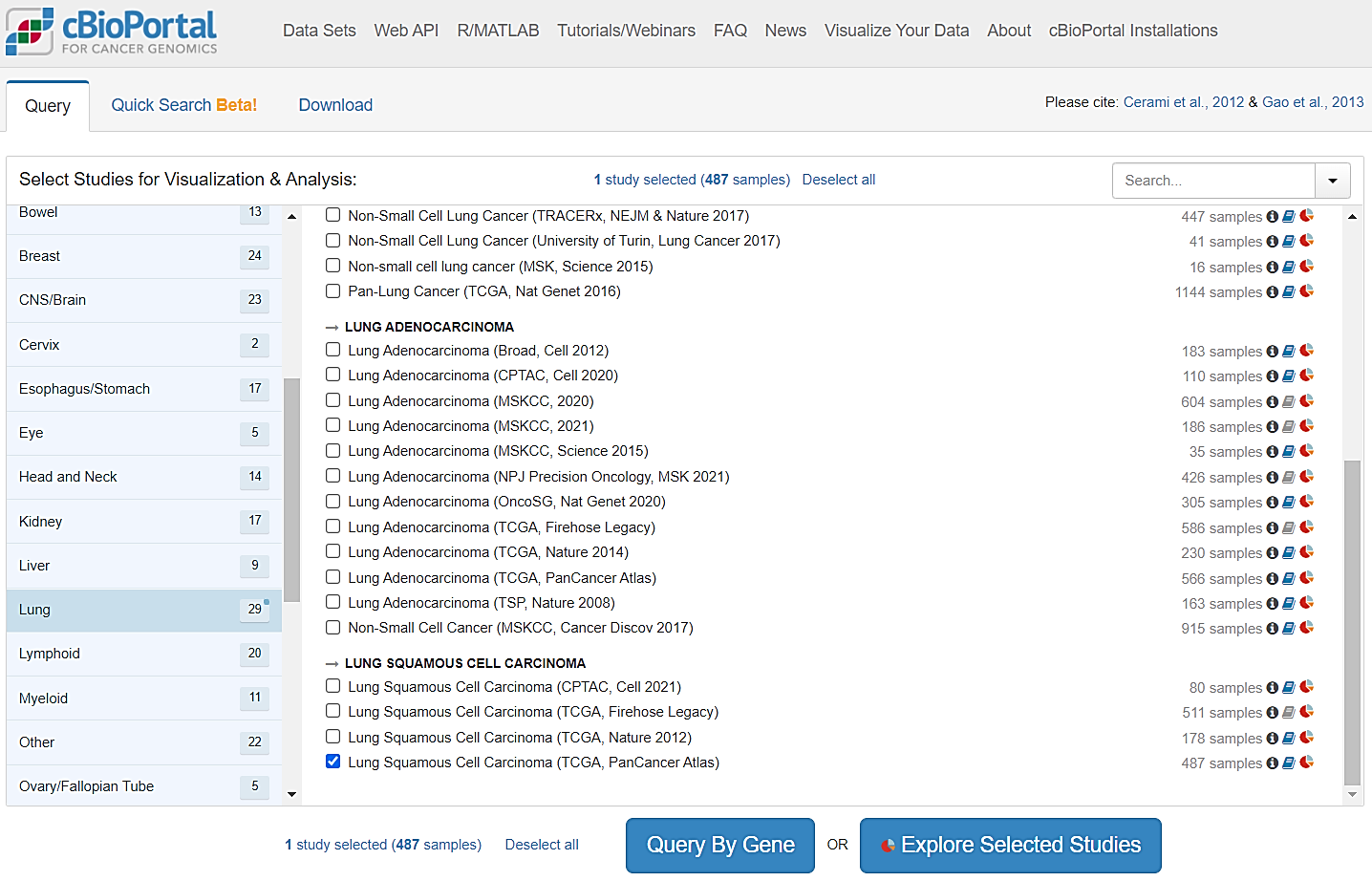

**Step3:** Select specific case study e.g., Lung Squamous Cell Carcinoma (TCGA, PanCancer Atlas)

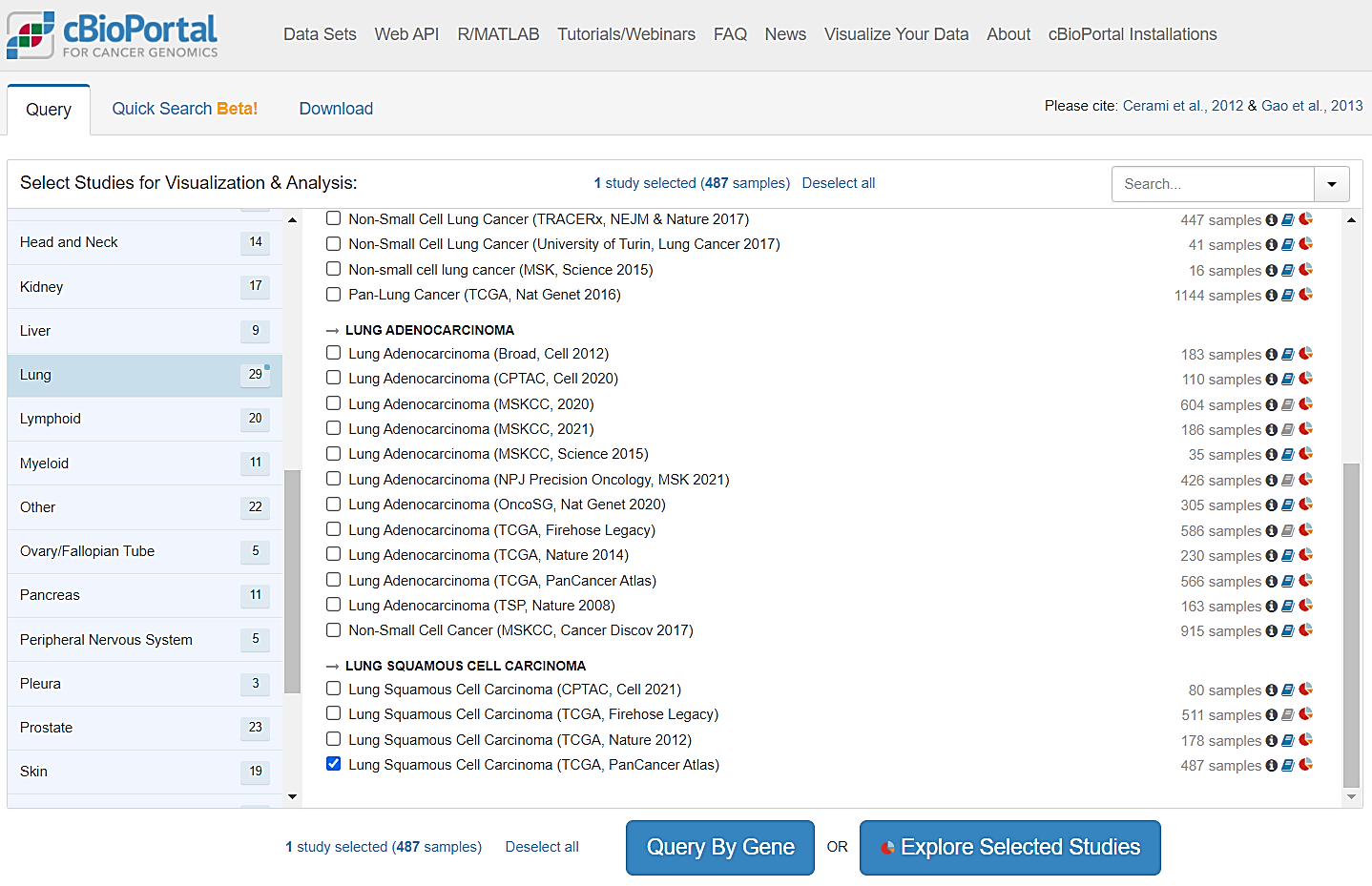

**Step4:** Click on "Explore Selected Studies" button

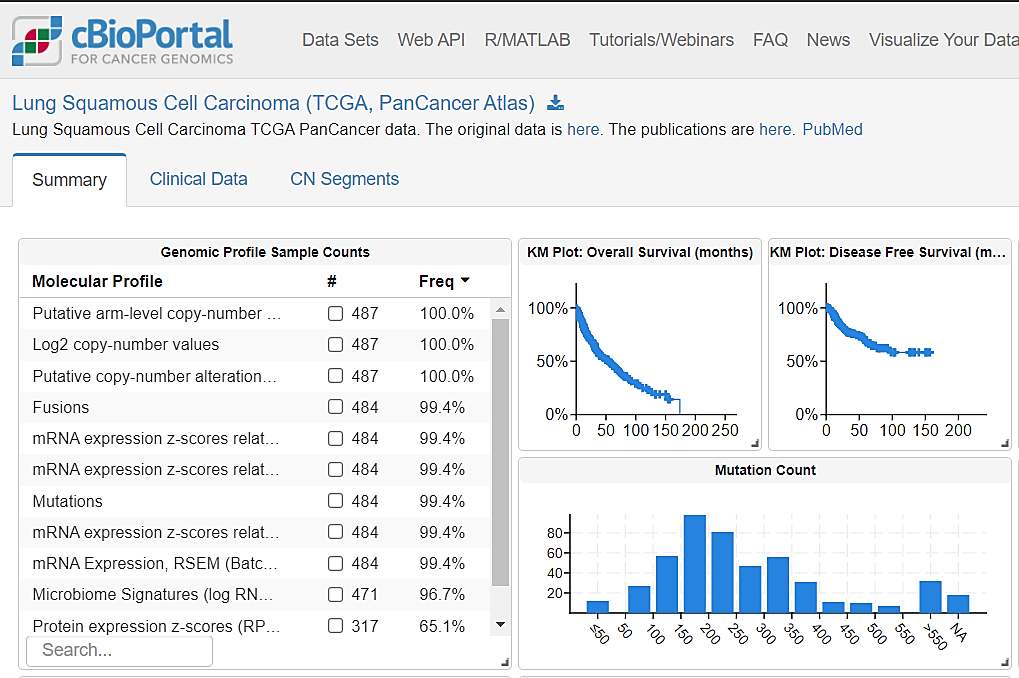

**Step5:** Click "Download Icon"

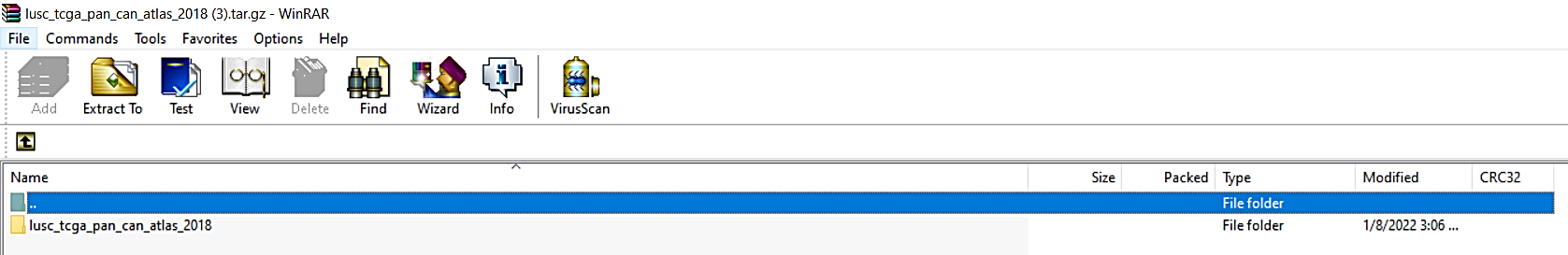

**Step6:** Open the downloaded zip folder

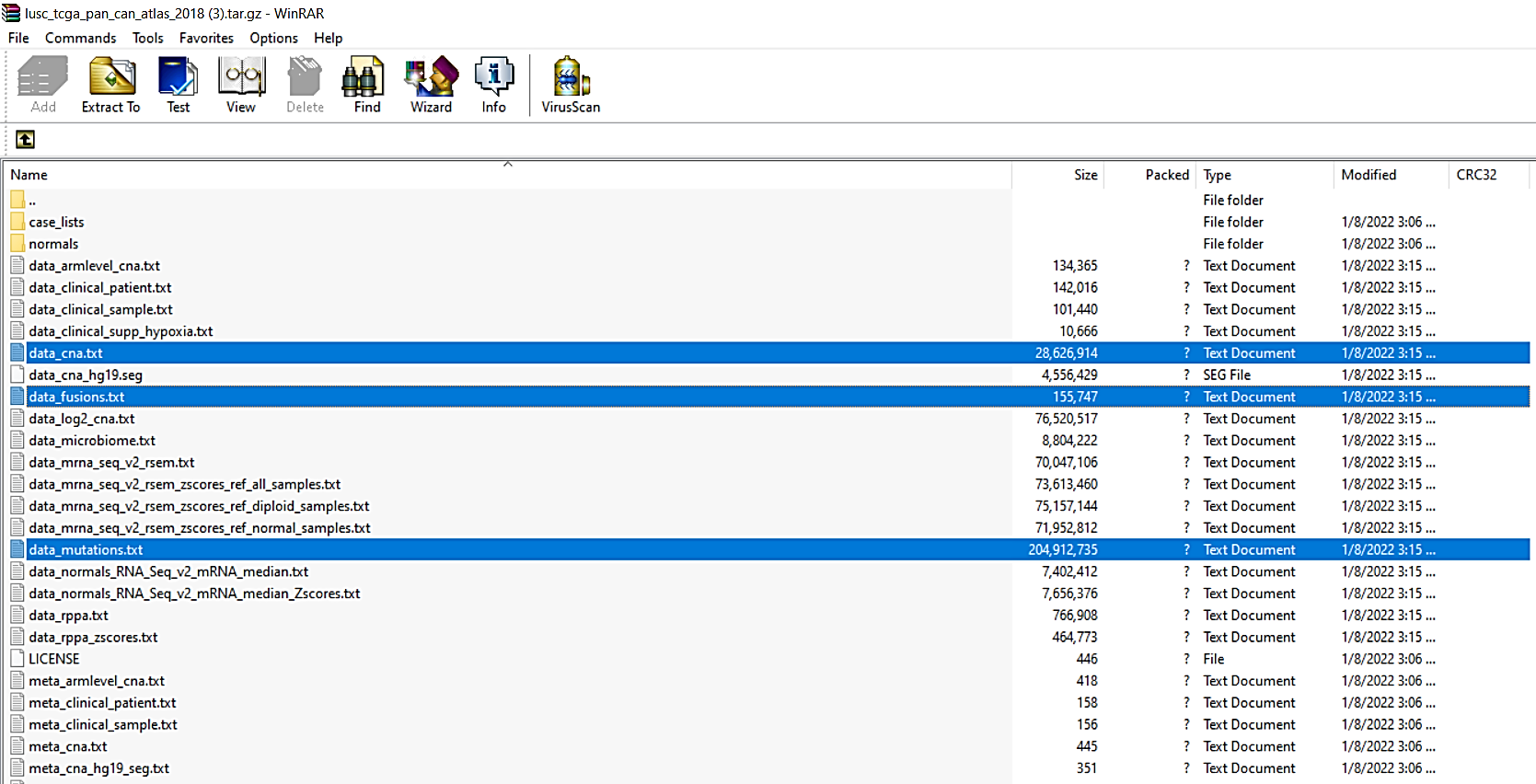

**Step7:** Select the files containing patient information on copy number variations (data_cna.txt), genomic structural variants (data_fusions.txt), and somatic mutations (data_mutations.txt)
