## Supplementary material for "CanSeer: A Method for Development and Clinical Translation of Personalized Cancer Therapeutics": Step by Step Guide: Step_by_Step_Guide_3.docx

### Step-by-Step Guidelines for Downloading RNA-seq based Gene Expression Data from GDC Data Portal

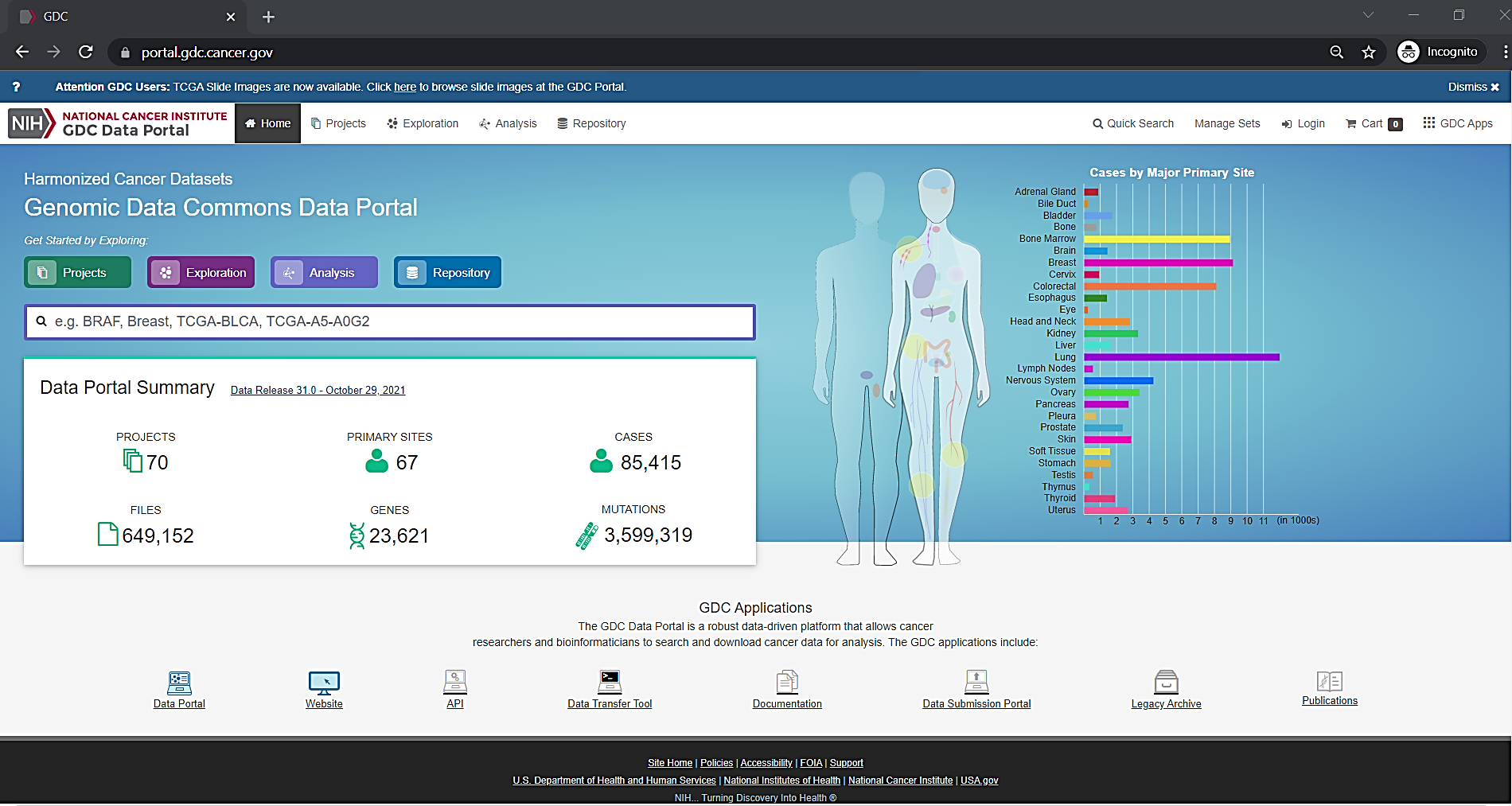

**Step1:** Go to https://portal.gdc.cancer.gov/

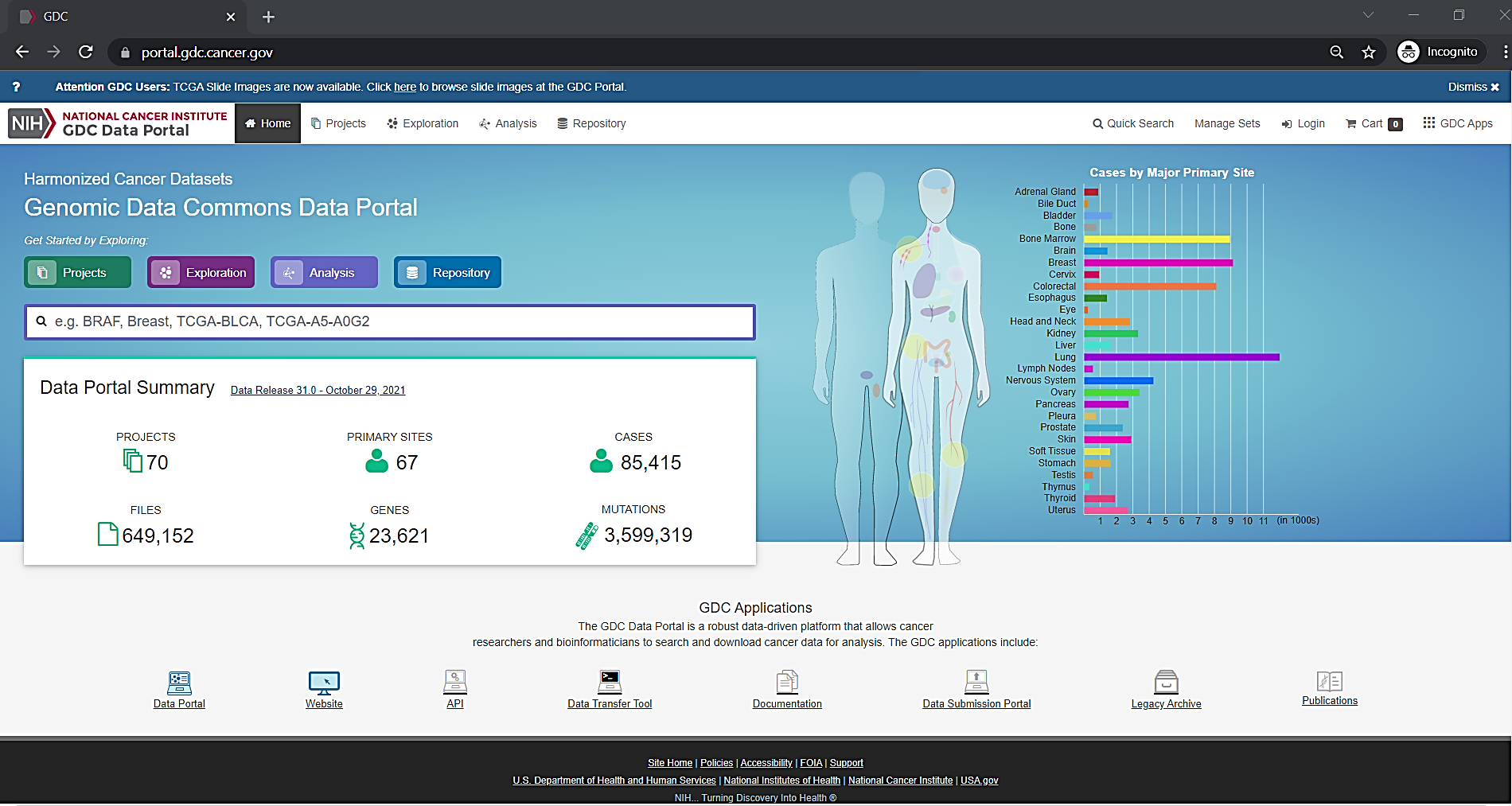

**Step2:** Click on "Repository" button

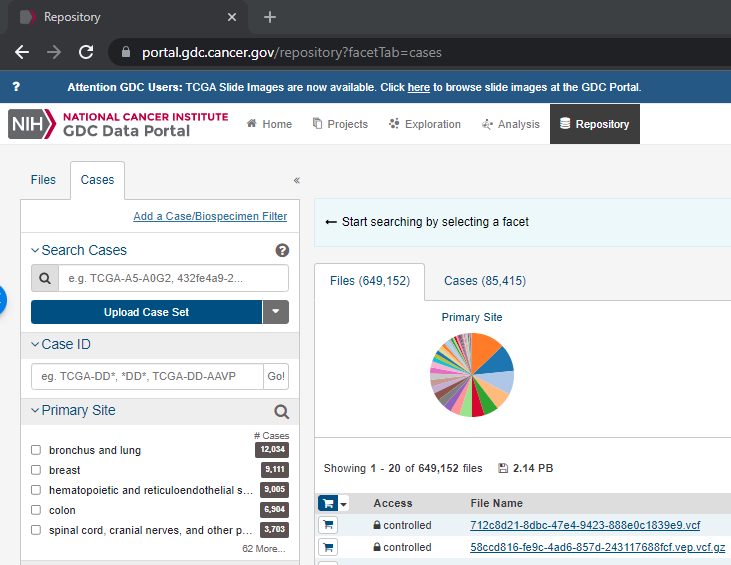

**Step3:** Select "Cases"

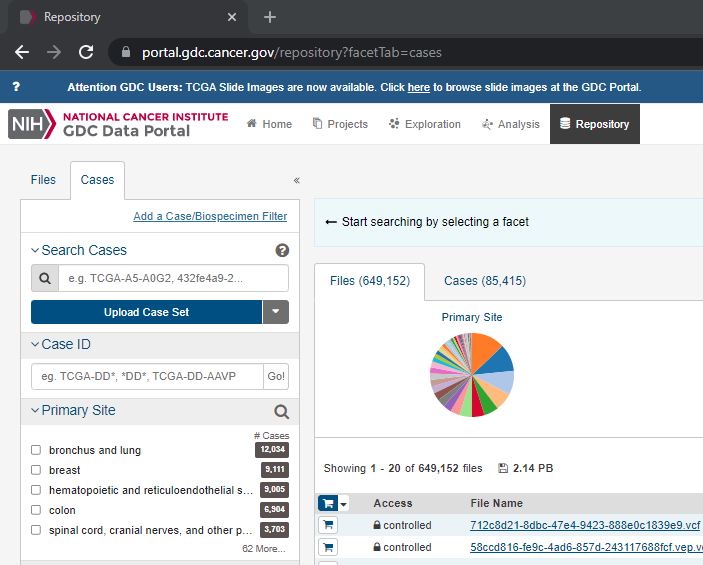

**Step4:** Click on "Add a Case/Biospecimen Filter"

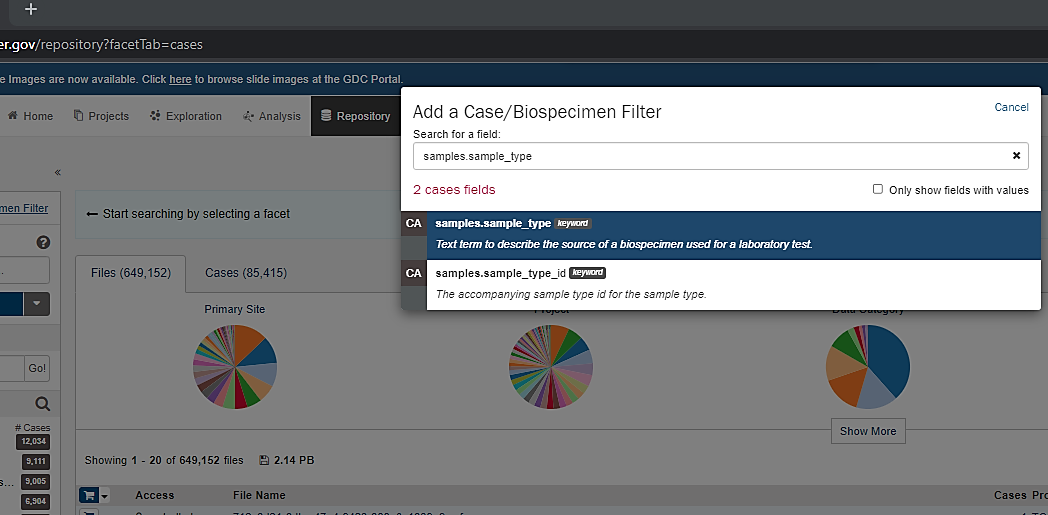

**Step5:** Type "samples.sample_type" in search field and select it

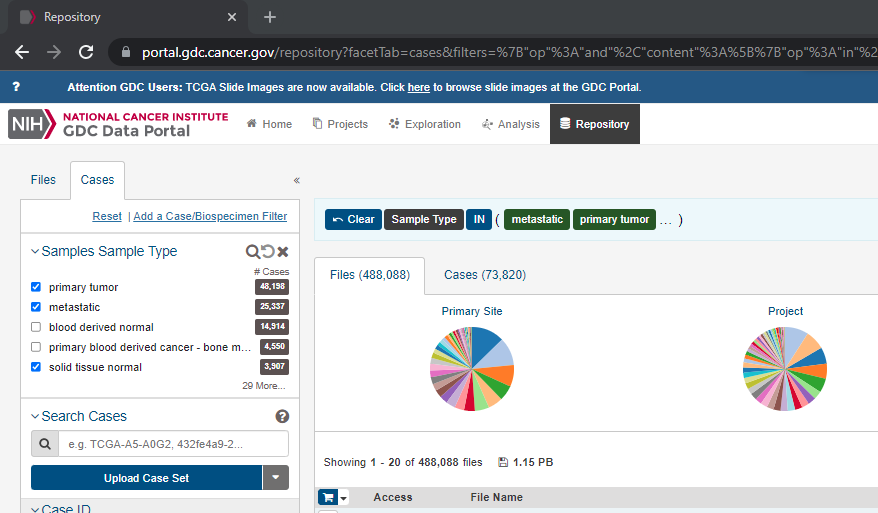

**Step6:** Select the sample types e.g. primary tumor, metastasis and solid tissue normal

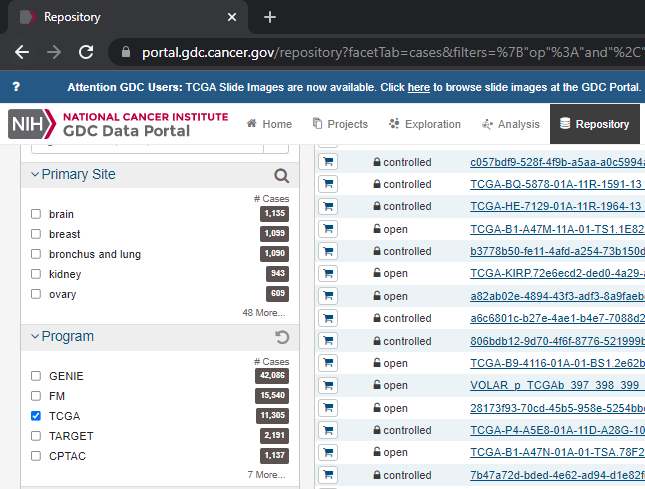

**Step7:** Select the Program e.g. TCGA

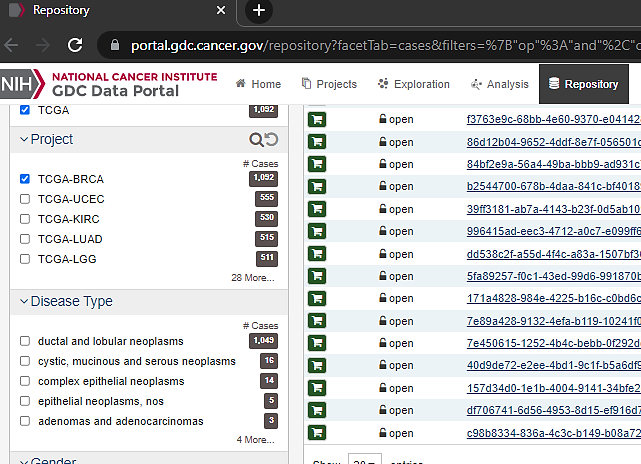

**Step8:** Select the Project e.g. TCGA-BRCA

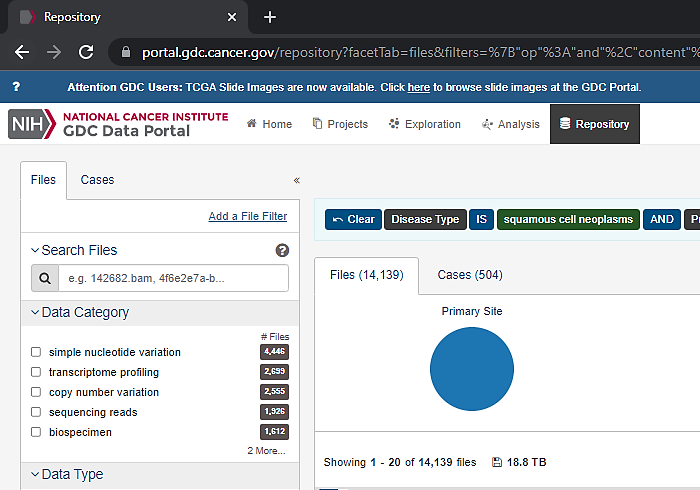

**Step9:** Select "Files"

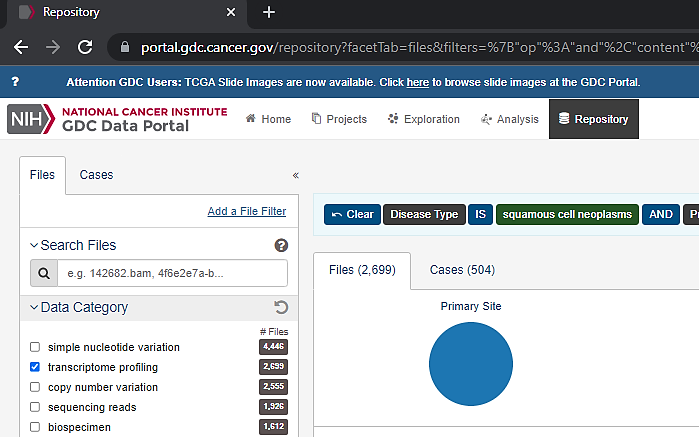

**Step10:** Select "transcriptome profiling" in Data Category

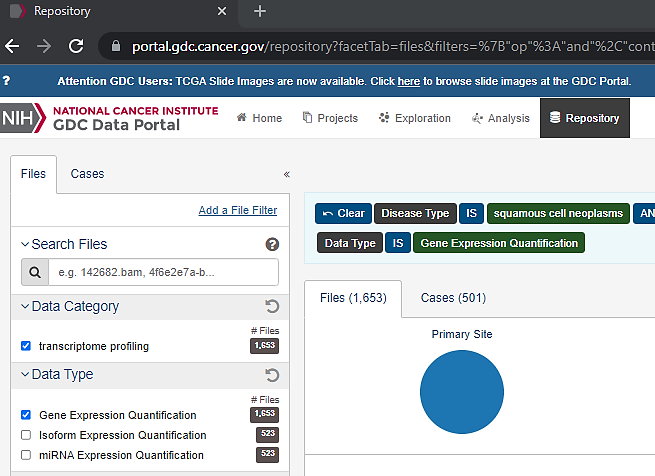

**Step11:** Select "Gene Expression Quantification" in Data Type

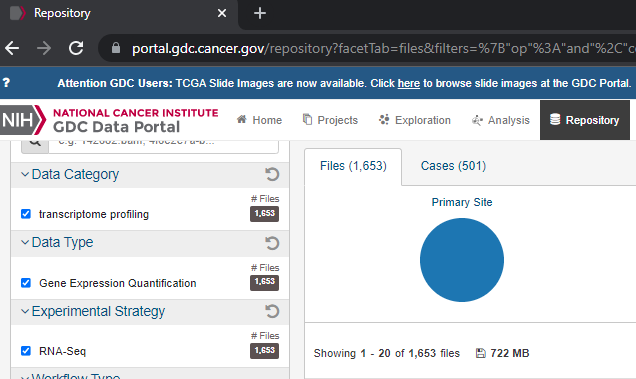

**Step12:** Select "RNA-seq" in Experimental Strategy

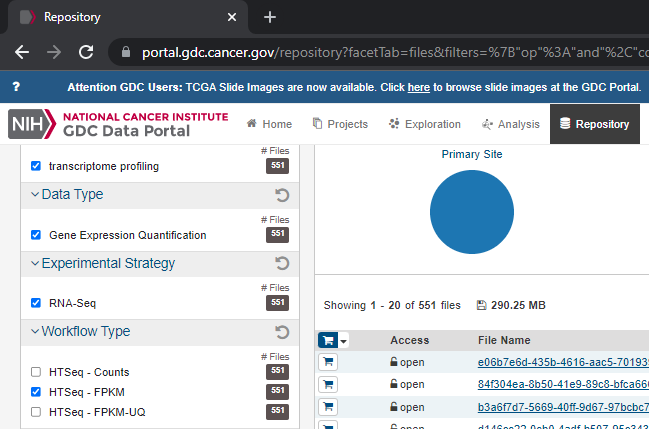

**Step13:** Select "HTSeq - FPKM" in Workflow Type

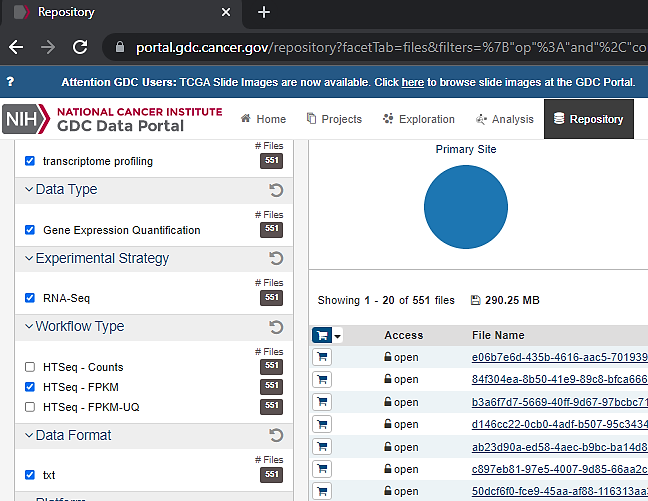

**Step14:** Select ".txt" in Data Format

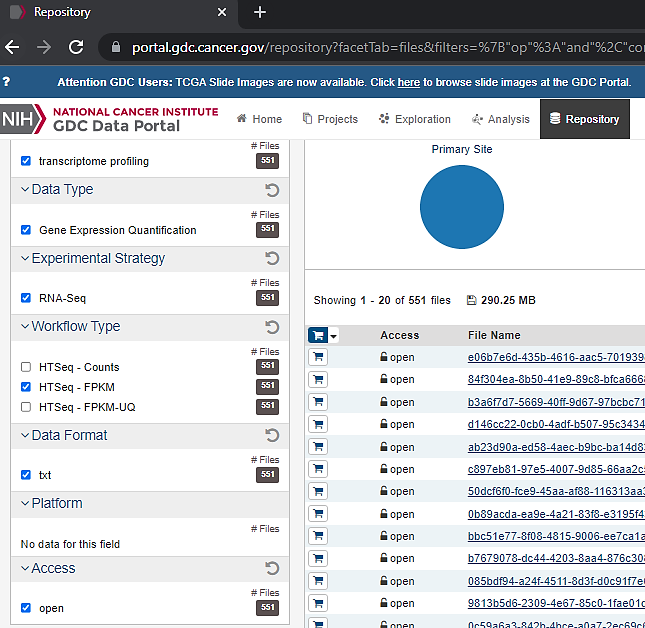

**Step15:** Select "Open" in Access

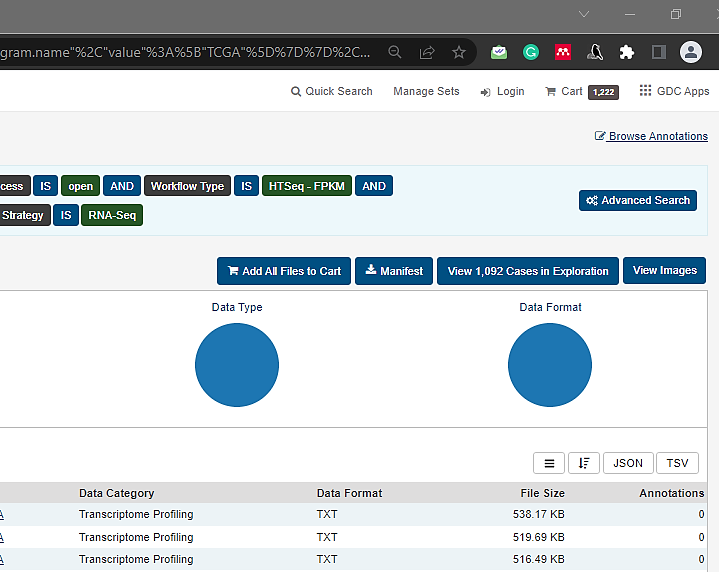

**Step16:** Click on "Add All Files to Cart"

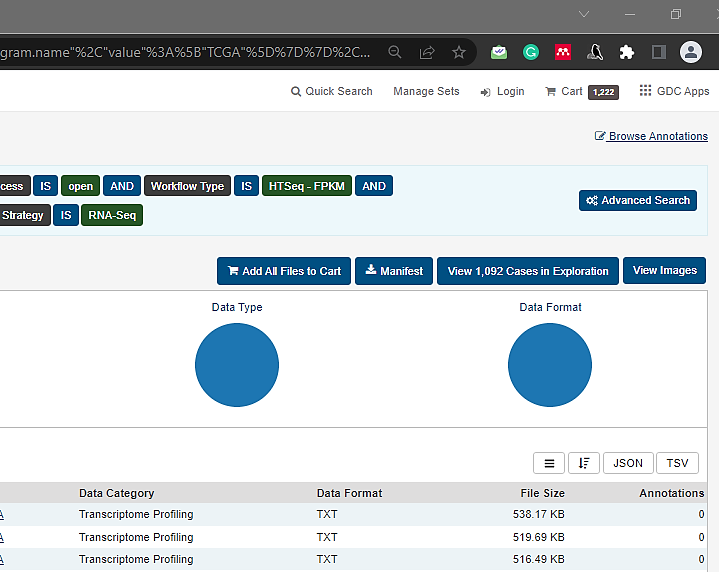

**Step17:** Click on "Cart"

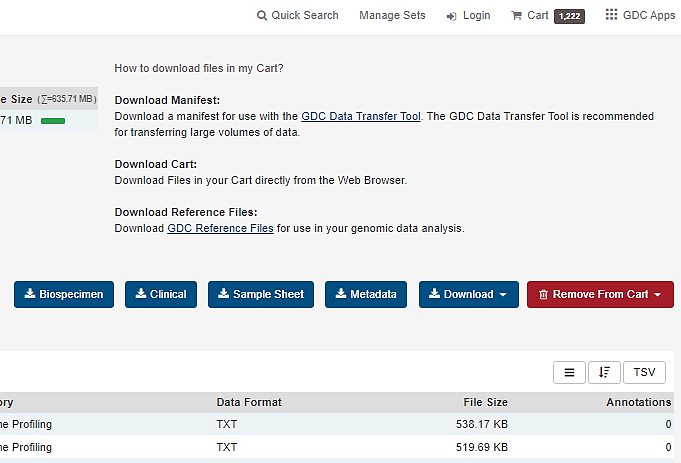

**Step18:** Select the "Sample Sheet" button to download it

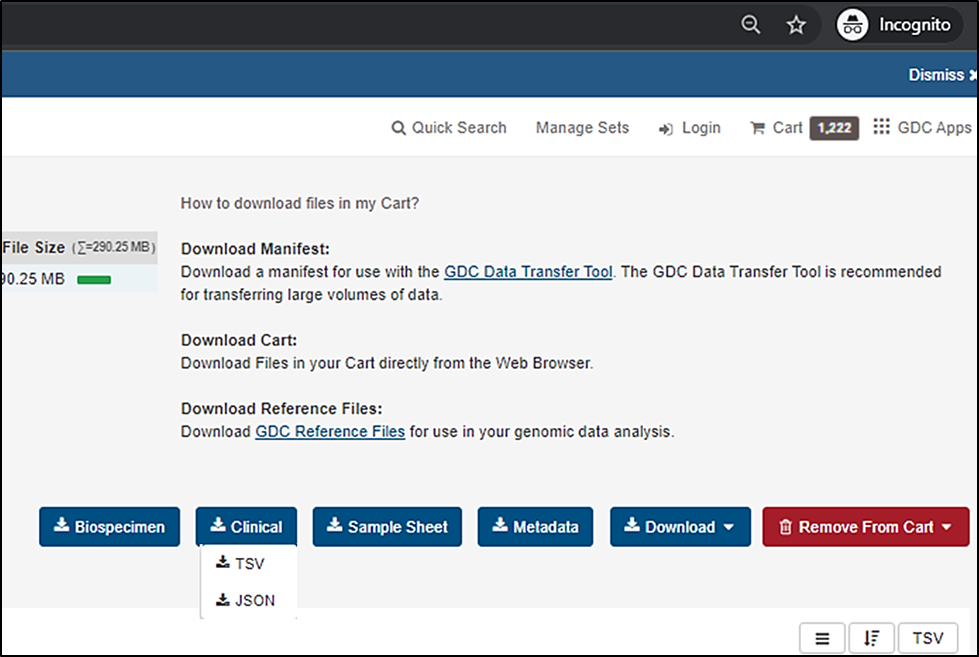

**Step19:** Click on "Clinical" button and select "TSV" to download clinical information

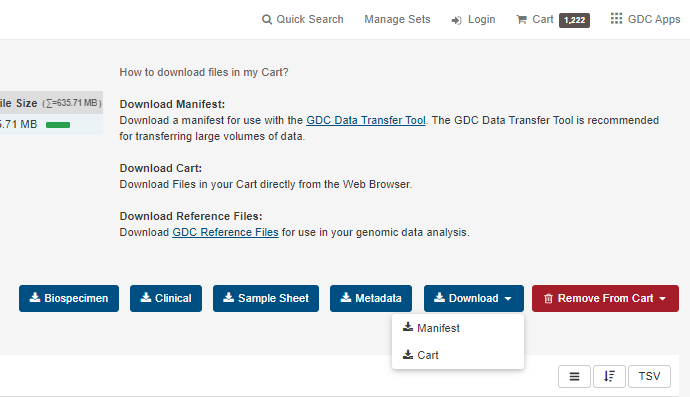

**Step20:** Click on the "Download" button and select "Cart"
