## Supplementary material for "CanSeer: A Method for Development and Clinical Translation of Personalized Cancer Therapeutics": Extraction_of_Raw_Expression_Only_Tumor.m

```
% This function takes parameters 'Network Genes list','Only Tumor CaseIDs' and 'Results_File'
% Returns the sorted Network genes array with Tumor Raw expression.

function [TumorRawExpressionFile]=Extraction_of_Raw_Expression_Only_Tumor(Path_Network_Genes,Sheet_T,Path_Expression_Data,Results_File)
tic;

progressbar('extracting gene expression');
progressbar(0.15);

% Get the path of expression data
Path_Exp_Data=Path_Expression_Data;

if("Empty"==Path_Exp_Data)
    display("Files not found; reset the path");
else
    % Read Network Genes list
    NetworkGenes=Path_Network_Genes;
    T2 = readtable(NetworkGenes);
    GenesList=table2cell(T2(:,1));
    NGenesSum=sum(T2{:,2}~="");
    NGenes=table2cell(T2(1:NGenesSum,2));
    [SortedNGenes,GenesInd]=intersect(GenesList,NGenes);
    
    % Assign Sheet_N and Sheet_T
    Sheet_Tumor=Sheet_T;
    % Get the TFNames and TCID
    TCID=Sheet_Tumor(2:end,1);
    TFNames= Sheet_Tumor(2:end,2);
    % Initialize the matrix for storing Tumor Expression
    Tumor_Exp=zeros(length(SortedNGenes),length(TCID));
    progressbar(0.3);
    % Read all the TFNames and extract expression
    for m=1:length(TCID)
        % Fetch Tumor FileNames from the Paired Sheet
        TFName= TFNames{m,1};
        TPath=char(TFName);
        newStr1 = split(TPath,'.gz');
        TF_Path=newStr1{1};
        TFPath=[char(Path_Exp_Data),char(TF_Path)];
        try
            % Read Expression files from the Paired CaseID List
            % Normal Exp stored in'NExp' and Tumor Exp stored in 'TExp'
            [EGNames1,TExp] = textread(TFPath,'%s %f');
            TExp=TExp(GenesInd);
            Tumor_Exp(:,m)=TExp;
        catch
            display('Tumor expression data file not found:');
            display(char(TF_Path));
            continue;
        end
    end
    
    progressbar(0.95);
    
        
    % Enlist NetworkGenes that are not matched with the Ensemble Gene List
    [G,I]=ismember(NGenes,SortedNGenes);
    MissingGenes=NGenes(find(~I));
    
    %For Tumor
    MissingGenesArrayTumor=zeros(length(MissingGenes),length(TCID));
    MissingGenesArrayTumor(MissingGenesArrayTumor==0) = NaN;
    MissingNetGenesArrayTumor=[MissingGenes,table2cell(array2table(MissingGenesArrayTumor))];    
    
    
    % Concatenate SortedNGeneNames and CaseIDs for 'T' with Expression
    NGenesTumorExp=[SortedNGenes,table2cell(array2table(Tumor_Exp))];
    Tumor_CID=['NetworkGenesSorted',transpose(TCID)];
    TumorRawExpressionFile=vertcat(Tumor_CID,NGenesTumorExp);
    % Vertically concatenate TumorRawExpressionFile with MissingNetGenesArray
    TumorRawExpressionFileUser=vertcat(TumorRawExpressionFile,MissingNetGenesArrayTumor);
    
    %% Write results data file %%
    Results_File_Path=cellstr(Results_File(1));
    UUID=cellstr(Results_File(2));
    % Tumor Raw Expression
    ResultsFilePath_T=strcat(char(Results_File_Path),'Only_Tumor_RawExpressionFile_',char(UUID),'.xlsx');
    xlswrite(ResultsFilePath_T,TumorRawExpressionFileUser);
     
    progressbar(1);
end
toc;
end
```
