## Supplementary material for "CanSeer: A Method for Development and Clinical Translation of Personalized Cancer Therapeutics": Extraction_of_Raw_Expression_Paired.m

```
% This function takes parameters 'Network Genes list','Paired CaseIDs' and 'Results_File'
% Returns the sorted Network genes array with NormalRawExpression and Tumor Raw expression.

function [NormalRawExpressionFile,TumorRawExpressionFile]= Extraction_of_Raw_Expression_Paired(Path_NetworkGenes,Filtered_Sheet,Path_Expression_Data,Results_File)
tic;

progressbar('extracting gene expression');
progressbar(0.15);
% Get the path of expression data
Path_Exp_Data=Path_Expression_Data;

if("Empty"==Path_Exp_Data)
    display("Files not found; reset the path");
else
    
    % Read Network Genes list
    NetworkGenes=Path_NetworkGenes;
    T2 = readtable(NetworkGenes);
    GenesList=table2cell(T2(:,1));
    NGenesSum=sum(T2{:,2}~="");
    NGenes=table2cell(T2(1:NGenesSum,2));
    [SortedNGenes,GenesInd]=intersect(GenesList,NGenes);
    
    % Get the Paired CaseIDs and FileNames
    Paired=Filtered_Sheet;
    
    % Third Column contains Normal File Names
    NFNames= Paired(2:end,3);
    NCID=Paired(2:end,2);
    Normal_Exp=zeros(length(SortedNGenes),length(NCID));
    
    % Third Column contains Tumor File Names
    TFNames= Paired(2:end,4);
    TCID=Paired(2:end,1);
    Tumor_Exp=zeros(length(SortedNGenes),length(TCID));
    progressbar(0.25);
    % Extracting the Raw Expression of Normal and Cancer Paired Samples
    for m=1:length(TCID)
        % Fetch Normal FileNames from the Paired Sheet
        NFName= NFNames{m,1};
        NPath=char(NFName);
        newStr = split(NPath,'.gz');
        NF_Path=newStr{1};
        NFPath=[char(Path_Exp_Data),char(NF_Path)];
        % Fetch Tumor FileNames from the Paired Sheet
        TFName= TFNames{m,1};
        TPath=char(TFName);
        newStr1 = split(TPath,'.gz');
        TF_Path=newStr1{1};
        TFPath=[char(Path_Exp_Data),char(TF_Path)];
        
        try
            % Read Expression files from the Paired CaseID List
            % Normal Exp stored in'NExp' and Tumor Exp stored in 'TExp'
            [EGNames,NExp] = textread(NFPath,'%s %f');
            [EGNames1,TExp] = textread(TFPath,'%s %f');
            NExp =NExp(GenesInd);
            TExp=TExp(GenesInd);
            Normal_Exp(:,m)=NExp;
            Tumor_Exp(:,m)=TExp;
        catch
            display('Expression data file not found:');           
            display(char(NF_Path));
            display(char(TF_Path));
        end
    end
    
    progressbar(0.9);
    
     % Enlist NetworkGenes that are not matched with the Ensemble Gene List
    [G,I]=ismember(NGenes,SortedNGenes);
    MissingGenes=NGenes(find(~I));
    
    % For Normal
    MissingGenesArrayNormal=zeros(length(MissingGenes),length(NCID));
    MissingGenesArrayNormal(MissingGenesArrayNormal==0) = NaN;
    MissingNetGenesArrayNormal=[MissingGenes,table2cell(array2table(MissingGenesArrayNormal))];
    
    %For Tumor
    MissingGenesArrayTumor=zeros(length(MissingGenes),length(TCID));
    MissingGenesArrayTumor(MissingGenesArrayTumor==0) = NaN;
    MissingNetGenesArrayTumor=[MissingGenes,table2cell(array2table(MissingGenesArrayTumor))];  
    
    % Concatenate SortedNGeneNames and CaseIDs for 'N' and 'T' with Expression
    NGenesNormalExp=[SortedNGenes,table2cell(array2table(Normal_Exp))];
    Normal_CID=['NetworkGenesSorted',transpose(NCID)];
    NormalRawExpressionFile=vertcat(Normal_CID,NGenesNormalExp);
    % Vertically concatenate NormalRawExpressionFile with MissingNetGenesArrayNormal
    NormalRawExpressionFileUser=vertcat(NormalRawExpressionFile,MissingNetGenesArrayNormal);
    
    NGenesTumorExp=[SortedNGenes,table2cell(array2table(Tumor_Exp))];
    Tumor_CID=['NetworkGenesSorted',transpose(TCID)];
    TumorRawExpressionFile=vertcat(Tumor_CID,NGenesTumorExp);
    % Vertically concatenate TumorRawExpressionFile with MissingNetGenesArrayTumor
    TumorRawExpressionFileUser=vertcat(TumorRawExpressionFile,MissingNetGenesArrayTumor);
    
    %% Write results data file %%
    Results_File_Path=cellstr(Results_File(1));
    UUID=cellstr(Results_File(2));
    % Normal Raw Expression
    ResultsFilePath_N=strcat(char(Results_File_Path),'Paired_NormalRawExpressionFile_',char(UUID),'.xlsx');
    xlswrite(ResultsFilePath_N,NormalRawExpressionFileUser);
    % Tumor Raw Expression
    ResultsFilePath_T=strcat(char(Results_File_Path),'Paired_TumorRawExpressionFile_',char(UUID),'.xlsx');
    xlswrite(ResultsFilePath_T,TumorRawExpressionFileUser);
    
    progressbar(1);
end
    toc;
end
```
