## Supplementary material for "CanSeer: A Method for Development and Clinical Translation of Personalized Cancer Therapeutics": Filter_Copy_Number_Variations.m

```
% Note: changem() requires mapping toolbox

% This function takes three parameters 'Path_Network_Genes','Paired_File','Path_CNV' and 'Results_File'
% Returns the logical array of CNV for Tumor samples [Rows represent sorted list of Network Genes and Columns represents the order of CID in paired sample file]

function [CNV_LA]= Filter_Copy_Number_Variations(Path_Network_Genes,Sheet_Filtered,Path_CNV,Results_File)
% Time calculator
tic;

% Intersect network genes with CNVGenesList to find the index
% Network_Genes_List=Path_Network_Genes;
% NetworkGenes=readtable(Network_Genes_List);
% NetGenesCount=sum(NetworkGenes{:,2}~= "");
% Net_GNames=table2cell(NetworkGenes(1:NetGenesCount,2));

% Read Network Genes list
NetworkGenes=Path_Network_Genes;
T2 = readtable(NetworkGenes);
GenesList=table2cell(T2(:,1));
NGenesSum=sum(T2{:,2}~="");
NGenes=table2cell(T2(1:NGenesSum,2));
[SortedNGenes,GenesInd]=intersect(GenesList,NGenes);
Net_GNames=SortedNGenes;

% Get the paired samples from the sample sheet
% Note: If we make a separate function to get the logical array of CNV for
% our network genes and paired patients then we have to call functions

Filtered_Sample_Sheet=Sheet_Filtered;
% Number of columns in sheet differs in paired,4 unpaired_T,2 and
% OnlyTumor,3
if(size(Sheet_Filtered,2)==4)
    Tumor_CaseID=Filtered_Sample_Sheet(2:end,2);
elseif(size(Sheet_Filtered,2)==2)
    Tumor_CaseID=Filtered_Sample_Sheet(2:end,1);
else
    Tumor_CaseID=Filtered_Sample_Sheet(2:end,1);
end


% Get Mutations Data ColA and ColQ, mutated genes and SampleIDs stored in  'ColA1' and 'ColQ1'
Path_CNV=Path_CNV;

if(strlength(Path_CNV)==0)
    % Return empty logical array for CNV of same length
    CNV_LA=zeros(length(Net_GNames),length(Tumor_CaseID));
    display("CNV data is missing");
else
    % Show progress bar
    progressbar('loading CNV data');
    progressbar(0.15);
    
    CNV=importdata(Path_CNV);
    
    progressbar(0.75);
    % # of Genes=Row-1 (Row1 is 'HugoSymbol') and # of CIDs= Col-2 (Col1 is'HugoSymbol'
    % and COl2 'EntrezID')
    [Rows,Cols]=size(CNV.textdata);
    [Rows1,Cols1]=size(CNV.data);
    CNV_CID= CNV.textdata(1,3:Cols);
    CNV_Genes=CNV.textdata(2:Rows,1);
    CNV_Data=CNV.data(:,2:Cols1);
    [SortedNetGenes,CNV_ALLGenesIndex]=intersect(CNV_Genes,Net_GNames);
   
    % Not all network genes have CNV 
    
    CNV_Filtered=zeros(length(Net_GNames),length(Tumor_CaseID));
    % To bring coherence
    [genes,Ind]=intersect(SortedNGenes,SortedNetGenes)
    
    for i=1:length(Tumor_CaseID)
        LogicalArray=strfind(CNV_CID,char(Tumor_CaseID(i)));
        Col_Ind=find(~cellfun(@isempty,LogicalArray));
        if Col_Ind > 0
            Temp=CNV_Data(CNV_ALLGenesIndex,Col_Ind);
            %CNV_Filtered(:,i) =Temp; 
            
            CNV_Filtered(Ind,i) =Temp;
        end
    end
    
    progressbar(0.9);
    

    % Now replace the numbers in CNV_Filtered with '1' and '0' by looking at the
    % LOOKUP TABLE [-2 -1 0 1 2]
    % -2 or 2 = 1  -1 or 0 or 1= 0
    Unique=unique(CNV_Filtered);
    Lookup_T=[-2 -1 0 1 2] ;
    New_T=[1 0 0 0 1];
    LA=ismember(Lookup_T,Unique);
    New=New_T(find(LA));
    CNV_LA=changem(CNV_Filtered,New,Unique);
    % For users to discreminate the -2 and 2
    Users_T=[-2 0 0 0 2];
    Users_New=Users_T(find(LA));
    Users_CNV_LA=changem(CNV_Filtered,Users_New,Unique);
    
    % Add Headers in the CNV logical array
    Tumor_CNV_LA=[['NetworkGenes';sort(Net_GNames)],[transpose(Tumor_CaseID);num2cell(Users_CNV_LA)]];
    
%     % Enlist NetworkGenes that are not matched with the Ensemble Gene List
%     [G,I]=ismember(SortedNGenes,SortedNetGenes);
%     MissingGenes=NGenes(find(~I));  
%     %For Tumor
%     MissingGenesArrayTumor=zeros(length(MissingGenes),length(Tumor_CaseID));
%     MissingGenesArrayTumor(MissingGenesArrayTumor==0) = NaN;
%     MissingNetGenesArrayTumor=[MissingGenes,table2cell(array2table(MissingGenesArrayTumor))];  
%     % Vertically concatenate  Users_CNV_LA with MissingNetGenesArrayTumor
%     TumorCNV_LA_User=vertcat(Tumor_CNV_LA,MissingNetGenesArrayTumor);
    %% Write results data file %%
    Results_File_Path=cellstr(Results_File(1));
    UUID=cellstr(Results_File(2));
    
    % Tumor Raw Expression
    ResultsFilePath_T=strcat(char(Results_File_Path),'Tumor_CNV_Logical_Array_',char(UUID),'.xlsx');
    xlswrite(ResultsFilePath_T,Tumor_CNV_LA);
    
    %%
    % Close progress bar
    progressbar(1);    
end
toc;

end
```
