## Supplementary material for "CanSeer: A Method for Development and Clinical Translation of Personalized Cancer Therapeutics": Filter_SomaticMutations.m

```
% This function takes input parameters 'Path_Network_Genes','Paired_File', 'Path_CNV'and 'Results_File'
% Returns the logical array of SomaticMutation for Tumor samples [Rows represent sorted list of Network Genes and Columns represents the order of CID in paired sample file]

function [SomaticMutation_LA]=Filter_SomaticMutations(Path_Network_Genes,Sheet_Filtered,Path_SomaticMutations,Results_File)
tic;
% Intersect network genes with CNVGenesList to find the index
% Network_Genes_List=Path_Network_Genes;
% NetworkGenes=readtable(Network_Genes_List);
% NetGenesCount=sum(NetworkGenes{:,2}~= "");
% Net_GNames=table2cell(NetworkGenes(1:NetGenesCount,2));

% Read Network Genes list
NetworkGenes=Path_Network_Genes;
T2 = readtable(NetworkGenes);
GenesList=table2cell(T2(:,1));
NGenesSum=sum(T2{:,2}~="");
NGenes=table2cell(T2(1:NGenesSum,2));
[SortedNGenes,GenesInd]=intersect(GenesList,NGenes);
Net_GNames=SortedNGenes;

% Get the paired samples from the sample sheet
% Note: If we make a separate function to get the logical array of CNV for
% our network genes and paired patients then we have to call functions
Filtered_Sample_Sheet=Sheet_Filtered;
% Number of columns differs in paired_Sample_Sheet,4 unpaired_Sample_Sheet,2 and
% OnlyTumor_Sample_Sheet,3
if(size(Sheet_Filtered,2)==4)
    Tumor_CaseID=Filtered_Sample_Sheet(2:end,2);
elseif(size(Sheet_Filtered,2)==2)
    Tumor_CaseID=Filtered_Sample_Sheet(2:end,1);
else
    Tumor_CaseID=Filtered_Sample_Sheet(2:end,1);
end

% Get Mutations Data ColA and ColQ, mutated genes and SampleIDs stored in  'ColA1' and 'ColQ1'
SomaticMutationsList=Path_SomaticMutations;
% To check if CNV file is empty or not
if(strlength(SomaticMutationsList)==0)
    % Return empty logical array for somatic mutations of same length
    SomaticMutation_LA=zeros(length(Net_GNames),length(Tumor_CaseID));
    display("Somatic mutations data is missing");
else
    % Show progress bar
    progressbar('Loading somatic mutations data');
    progressbar(0.1);
    progressbar(0.15);
    S_Mutations_T=readtable(SomaticMutationsList);
    fieldnames(S_Mutations_T);
    ColA1 = S_Mutations_T.('Hugo_Symbol');
    ColQ1=S_Mutations_T.('Tumor_Sample_Barcode');
    NewTable=[ColA1,ColQ1];
    progressbar(0.85);
    % Sort rows of NewTable based on 'Tumor_Sample_Barcode'
    SortedT=sortrows(NewTable,2);
    SMutGenesAll=SortedT(:,1);
    
    SomaticMutation_LA=zeros(length(Net_GNames),length(Tumor_CaseID));
    for i=1:length(Tumor_CaseID)
        indexfind=strfind(SortedT(:,2),char(Tumor_CaseID(i)));
        IndList=find(~cellfun(@isempty,indexfind));
        %All mutated genes per CID
        MutatedGenesCID=SMutGenesAll(IndList);
        [NetGenesSortedMutList,NetGenesInd]=intersect(MutatedGenesCID,Net_GNames);
        SomaticMutation_LA(:,i)=ismember(sort(Net_GNames),NetGenesSortedMutList);
    end
        progressbar(0.9);
    % Add Headers in the Somatic Mutations logical array
    Tumor_Somatic_Mutation_LA=[['NetworkGenes';sort(Net_GNames)],[transpose(Tumor_CaseID);num2cell(SomaticMutation_LA)]];
    
    %% Write results data file %%
    Results_File_Path=cellstr(Results_File(1));
    UUID=cellstr(Results_File(2));
    
    % Tumor Raw Expression
    ResultsFilePath_T=strcat(char(Results_File_Path),'Tumor_Somatic_Mutations_Logical_Array_',char(UUID),'.xlsx');
    xlswrite(ResultsFilePath_T,Tumor_Somatic_Mutation_LA);
    
    %%
    % Close progress bar
    progressbar(1);
end
toc;
end
```
