## Supplementary material for "CanSeer: A Method for Development and Clinical Translation of Personalized Cancer Therapeutics": Filtering_CIDs_Only_Tumor.m

```
% This function takes 'Path_Sample_Sheet' and 'Results_File' as the input argument
% Return the output arguments 'Filtered_Sheet' and 'Reselect_var'

function [Filtered_Sheet,Reselect_var]=Filtering_CIDs_Only_Tumor(Path_Sample_Sheet,Results_File)
% Time calculator
tic;
Path_Sheet=Path_Sample_Sheet;
if("Empty"==Path_Sheet)
    %display("File not found; reset the path");
else
    
    Sample_Sheet=readtable(Path_Sample_Sheet);
    Sample_Sheet_T=sortrows(Sample_Sheet,7);
    fieldnames(Sample_Sheet_T);
    Sample_Type=Sample_Sheet_T.('SampleType');
    Case_ID=Sample_Sheet_T.('CaseID');
    File_Name=Sample_Sheet_T.('FileName');
    Filtered_Sheet=[];
    NCID=[];
    TCID=[];
    NFName=[];
    TFName=[];
    NSType=[];
    TSType=[];
    Reselect_var=0;
    Filtered_Sheet=[];
    [Unique_Sample_Type]=unique(Sample_Type);
    % This block of code will filter the normal
    for i=1:length(Unique_Sample_Type)
        % Check which of these key words are present in the unique sample types
        % at index i
        Primary=strfind(Unique_Sample_Type(i),'Primary');
        Tumor=strfind(Unique_Sample_Type(i),'Tumor');
        Metastatic=strfind(Unique_Sample_Type(i),'Metastatic');
        % Would not cater recurrent and normal so this should be empty
        Recurrent=strfind(Unique_Sample_Type(i),'Recurrent');
        Normal=strfind(Unique_Sample_Type(i),'Normal');
        
        % This condition will skip 'Recurrent' or 'Normal' sample types
        if(logical(1)== ((~cellfun('isempty',Primary)) | (~cellfun('isempty',Tumor)) |(~cellfun('isempty',Metastatic))) & ((cellfun('isempty',Recurrent))))
            % Return logical array by comparing each sample type in the whole
            % sample types list
            LA=ismember(Sample_Type,Unique_Sample_Type(i));
            % Find and return the tumor CIDs
            Tumor_CID=Case_ID(find(LA));
            % Stored in the cell array by vertical concatenation
            TCID=[TCID;Tumor_CID];
            % Find and return the tumor filenames
            Tumor_FName=File_Name(LA);
            TFName=[TFName;Tumor_FName];
            % Find and return the tumor sample types
            T_Sample_Type=Sample_Type(find(LA));
            TSType=[TSType;T_Sample_Type];
            
        end
    end
    
    if(logical(1) == (~cellfun('isempty',{TCID})))   
        
       tryagain=1;
       while tryagain>0 
           
           
           prompt = {'Enter 1 for random sample selection and 2 for all samples from filtered only tumor samples:'};
           dlgtitle = 'Random sampling';
           dims = [1 70];
           definput = {'1 or 2'};
           answer = inputdlg(prompt,dlgtitle,dims,definput);
           Rand_All=str2num(answer{1});   
 
           
        % Giving user a choice for random or whole cohort selection  
        %Rand_All = input('Enter 1 for random sample selection and 2 for all samples from filtered only tumor samples:  ');
        
        switch  Rand_All
            case 1
                prompt = {'Enter number of random samples for tumor samples:'};
                dlgtitle = 'Random samples for only tumor';
                dims = [1 70];
                definput = {'10'};
                answer = inputdlg(prompt,dlgtitle,dims,definput);
                Random_T=str2num(answer{1});
                
                %Random_T = input('Enter number of random samples for tumor samples:');
                if(Random_T<length(TCID))
                % Random selection of tumor samples
                Tumor_T= randperm(length(TCID));
                Tumor_Idx=Tumor_T(1:Random_T);
                Rand_Selected_TCID=TCID(Tumor_Idx);
                Rand_Selected_TFName=TFName(Tumor_Idx);
                Rand_Selected_TSType=TSType(Tumor_Idx);
                Sheet_Tumor=[Rand_Selected_TCID,Rand_Selected_TFName,Rand_Selected_TSType];
                tryagain=0;
                else 
                uiwait(msgbox('Your random sample size exceeds the number of tumor patients; reselect a smaller sample'));
                tryagain=1;
                end
            case 2         
                Sheet_Tumor=[TCID,TFName,TSType];
                tryagain=0;
            case 3 
                %display('Enter correct option for random or whole sample selection! '); 
                uiwait(msgbox('Enter correct option for random or whole sample selection!'));
                tryagain=1;
        end
     end
        % ONLY TUMOR
        Filtered_Sheet=[{'Tumor_CID', 'Tumor_FName','SampleType'};Sheet_Tumor];
        
        % Write results data file %%
        Results_File_Path=cellstr(Results_File(1));
        UUID=cellstr(Results_File(2));
        % Tumor sample sheet
        ResultsFilePath_T=strcat(char(Results_File_Path),'Only_Tumor_Sample_Sheet_',char(UUID),'.xlsx');
        xlswrite(ResultsFilePath_T,Filtered_Sheet);

        Reselect_var=0;
    else
        %display("Reselect the right option");
        uiwait(msgbox('Reselect the right option'));
        Reselect_var=1;
    end
end

toc;

end
```
