## Supplementary material for "CanSeer: A Method for Development and Clinical Translation of Personalized Cancer Therapeutics": Filtering_CIDs_Paired.m

```
% This function takes 'Path_Sample_Sheet' and 'Results_File' as the input argument
% Return the output arguments 'PairedSheet' and 'Reselect_var'

function [PairedSheet,Reselect_var]=Filtering_CIDs_Paired(Path_Sample_Sheet,Results_File)
% Time calculator
tic;
Path_Sheet=Path_Sample_Sheet;
if("Empty"==Path_Sheet)
    display("File not found; reset the path");
else
    
    Sample_Sheet=readtable(Path_Sample_Sheet);
    Sample_Sheet_T=sortrows(Sample_Sheet,7);
    fieldnames(Sample_Sheet_T);
    Sample_Type=Sample_Sheet_T.('SampleType');
    Case_ID=Sample_Sheet_T.('CaseID');
    File_Name=Sample_Sheet_T.('FileName');
    PairedSheet=[];
    NCID=[];
    TCID=[];
    NFName=[];
    TFName=[];
    NSType=[];
    TSType=[];
    Reselect_var=0;
    [Unique_Sample_Type]=unique(Sample_Type);
    % This block of code will filter the normal
    for i=1:length(Unique_Sample_Type)
        % to check if any of these sample types present in the list
        Normal=strfind(Unique_Sample_Type(i),'Normal');
        Primary=strfind(Unique_Sample_Type(i),'Primary');
        Tumor=strfind(Unique_Sample_Type(i),'Tumor');
        Metastatic=strfind(Unique_Sample_Type(i),'Metastatic');
        % Would not cater Recurrent so this should be empty
        Recurrent=strfind(Unique_Sample_Type(i),'Recurrent');
        
        if(logical(1)== (((~cellfun('isempty',Normal)) | (~cellfun('isempty',Primary)) | (~cellfun('isempty',Tumor)) |(~cellfun('isempty',Metastatic))) & (cellfun('isempty',Recurrent))))
            if (logical(1)==(~cellfun('isempty',Normal)))
                LA=ismember(Sample_Type,Unique_Sample_Type(i));
                Normal_CID=Case_ID(find(LA));
                Normal_FName=File_Name(LA);
                NCID=[NCID;Normal_CID];
                NFName=[NFName;Normal_FName];
                N_Sample_Type=Sample_Type(find(LA));
                NSType=[NSType;N_Sample_Type];
            else
                LA=ismember(Sample_Type,Unique_Sample_Type(i));
                Tumor_CID=Case_ID(find(LA));
                TCID=[TCID;Tumor_CID];
                Tumor_FName=File_Name(LA);
                TFName=[TFName;Tumor_FName];
                T_Sample_Type=Sample_Type(find(LA));
                TSType=[TSType;T_Sample_Type];
            end
        end
    end
    
    [Paired_TCID,Ind_TCID]=intersect(TCID,NCID);
    T_FName=TFName(Ind_TCID);
    if(logical(1) == (~cellfun('isempty',{Paired_TCID})))
        [Paired_NCID,Ind_NCID] =intersect(NCID,TCID);
        N_FName=NFName(Ind_NCID);
         
        tryagain=1;
       while tryagain>0 
           
           prompt = {'Enter 1 for random sample selection and 2 for all samples from filtered paired samples:'};
           dlgtitle = 'Random sampling';
           dims = [1 70];
           definput = {'1 or 2'};
           answer = inputdlg(prompt,dlgtitle,dims,definput);
           Rand_All=str2num(answer{1});   
           
              % Giving user a choice for random or whole cohort selection  
        %Rand_All = input('Enter 1 for random sample selection and 2 for all samples from filtered paired samples: ');
        
        switch  Rand_All
            case 1
                prompt = {'Enter size of random samples for both normal and tumor samples:'};
                dlgtitle = 'Random Samples for Normal and Tumor';
                dims = [1 70];
                definput = {'10'};
                answer = inputdlg(prompt,dlgtitle,dims,definput);
                Random_N=str2num(answer{1});
                if (Random_N<length(NCID))
                %Random_N = input('Enter number of random samples for both normal and tumor samples: ');
                % Random Normal Samples filtering out
                Normal_N = randperm(length(Paired_NCID));
                Normal_Idx=Normal_N(1:Random_N);
                Rand_Selected_NCID=Paired_NCID(Normal_Idx);
                Rand_Selected_FName=N_FName(Normal_Idx);
                S_Normal=[Rand_Selected_NCID,Rand_Selected_FName];
                
                % Random selection of tumor samples
                Tumor_T= randperm(length(TCID));
                Tumor_Idx=Tumor_T(1:Random_N);
                Rand_Selected_TCID=TCID(Tumor_Idx);
                Rand_Selected_TFName=TFName(Tumor_Idx);
                S_Tumor=[Rand_Selected_TCID,Rand_Selected_TFName];
                Random_Selected_Paired=[Rand_Selected_NCID,Rand_Selected_TCID,Rand_Selected_FName,Rand_Selected_TFName];
                % Get out of while loop
                tryagain=0;
                else
                uiwait(msgbox('Your random sample size exceeds the number of normal patients; reselect a smaller sample'));
                tryagain=1; 
                end
            case 2
                Random_Selected_Paired=[Paired_NCID,Paired_TCID,N_FName,T_FName];
                % Get out of while loop
                tryagain=0;   
                
            case 3
           %display('Enter correct option for random or whole sample selection! '); 
           uiwait(msgbox('Enter correct option for random or whole sample selection!'));
                tryagain=1;
        end
       end
           
        PairedSheet=[{'Normal_CID','Tumor_CID','Normal_FName', 'Tumor_FName'};Random_Selected_Paired];
        % Write results data file %%
        Results_File_Path=cellstr(Results_File(1));
        UUID=cellstr(Results_File(2));
        ResultsFilePath=strcat(char(Results_File_Path),'Paired_Sample_Sheet_',char(UUID),'.xlsx');
        xlswrite(ResultsFilePath,PairedSheet);
        
        Reselect_var=0;
    else
        %display("Your expression dataset does not contain paired tumor samples, select any other option");
        uiwait(msgbox('Your samples dataset does not contain paired normal and tumor patients, select any other option'));
        Reselect_var=2;
    end

end
toc;
end
```
