## Supplementary material for "CanSeer: A Method for Development and Clinical Translation of Personalized Cancer Therapeutics": Filtering_CIDs_UnPaired.m

```
% This function takes 'Path_Sample_Sheet' and 'Results_File' as the input argument
% Return the output arguments 'Sheet_N', 'Sheet_T' and 'Reselect_var'

function [Sheet_N,Sheet_T,Reselect_var]=Filtering_CIDs_UnPaired(Path_Sample_Sheet,Results_File)
% Time calculator
tic;
Path_Sheet=Path_Sample_Sheet;
if("Empty"==Path_Sheet)
    display("File not found; reset the path");
else
    
    Sample_Sheet=readtable(Path_Sample_Sheet);
    Sample_Sheet_T=sortrows(Sample_Sheet,7);
    fieldnames(Sample_Sheet_T);
    Sample_Type=Sample_Sheet_T.('SampleType');
    Case_ID=Sample_Sheet_T.('CaseID');
    File_Name=Sample_Sheet_T.('FileName');
    PairedSheet=[];
    NCID=[];
    TCID=[];
    NFName=[];
    TFName=[];
    NSType=[];
    TSType=[];
    Sheet_N=[];
    Sheet_T=[];
    Reselect_var=0;
    [Unique_Sample_Type]=unique(Sample_Type);
    % This block of code will filter the normal
    for i=1:length(Unique_Sample_Type)
        % to check if any of these sample types present in the list
        Normal=strfind(Unique_Sample_Type(i),'Normal');
        Primary=strfind(Unique_Sample_Type(i),'Primary');
        Tumor=strfind(Unique_Sample_Type(i),'Tumor');
        Metastatic=strfind(Unique_Sample_Type(i),'Metastatic');
        % Would not cater recurrent so this should be empty
        Recurrent=strfind(Unique_Sample_Type(i),'Recurrent');
        
        if(logical(1)== (((~cellfun('isempty',Normal)) | (~cellfun('isempty',Primary)) | (~cellfun('isempty',Tumor)) |(~cellfun('isempty',Metastatic))) & (cellfun('isempty',Recurrent))))
            if (logical(1)==(~cellfun('isempty',Normal)))
                LA=ismember(Sample_Type,Unique_Sample_Type(i));
                Normal_CID=Case_ID(find(LA));
                Normal_FName=File_Name(LA);
                NCID=[NCID;Normal_CID];
                NFName=[NFName;Normal_FName];
                N_Sample_Type=Sample_Type(find(LA));
                NSType=[NSType;N_Sample_Type];
            else
                LA=ismember(Sample_Type,Unique_Sample_Type(i));
                Tumor_CID=Case_ID(find(LA));
                TCID=[TCID;Tumor_CID];
                Tumor_FName=File_Name(LA);
                TFName=[TFName;Tumor_FName];
                T_Sample_Type=Sample_Type(find(LA));
                TSType=[TSType;T_Sample_Type];
            end
        end
    end
    
    % This condition will be true when NCID is present
    if (logical(1) == (~cellfun('isempty',{NCID})))
        
        tryagain=1;
        while tryagain>0
            
           prompt = {'Enter 1 for random sample selection and 2 for all samples from filtered unpaired tumor samples:'};
           dlgtitle = 'Random sampling';
           dims = [1 70];
           definput = {'1 or 2'};
           answer = inputdlg(prompt,dlgtitle,dims,definput);
           Rand_All=str2num(answer{1});   
 
            
            % Giving user a choice for random or whole cohort selection
            %Rand_All = input('Enter 1 for random sample selection and 2 for all samples from filtered unpaired tumor samples: ');
            
            switch  Rand_All
                case 1
                    
                    prompt = {'Enter size of random samples for normal samples:','Enter size of random samples for tumor samples: '};
                    dlgtitle = 'Random Sample';
                    dims = [1 70];
                    definput = {'10','20'};
                    answer = inputdlg(prompt,dlgtitle,dims,definput);
                    Random_N=str2num(answer{1});
                    Random_T=str2num(answer{2});
                    %Random_N = input('Enter number of random samples for normal samples: ');
                    %Random_T = input('Enter number of random samples for tumor samples: ');
                    if (Random_N<length(NCID) & Random_T<length(TCID))
                    % Random Normal Samples filtering out
                    Normal_N = randperm(length(NCID));
                    Normal_Idx=Normal_N(1:Random_N);
                    Rand_Selected_NCID=NCID(Normal_Idx);
                    Rand_Selected_FName=NFName(Normal_Idx);
                    S_Normal=[Rand_Selected_NCID,Rand_Selected_FName];
                    
                    % Random selection of tumor samples
                    Tumor_T= randperm(length(TCID));
                    Tumor_Idx=Tumor_T(1:Random_T);
                    Rand_Selected_TCID=TCID(Tumor_Idx);
                    Rand_Selected_TFName=TFName(Tumor_Idx);
                    S_Tumor=[Rand_Selected_TCID,Rand_Selected_TFName];
                    
                    tryagain=0;
                    else
                     uiwait(msgbox('Your random sample size exceeds the number of normal and tumor patients; reselect a smaller sample'));
                     tryagain=1; 
                    end
                case 2
                    S_Normal=[NCID,NFName];
                    S_Tumor=[TCID,TFName];
                    
                    tryagain=0;
                case 3
                    %display('Enter correct option for random or whole sample selection! ');
                    uiwait(msgbox('Enter correct option for random or whole sample selection!'));
                    tryagain=1;
            end
        end
        
        % Return Normal and Tumor CIDs and FNames separately
        Header_N={'Normal_CaseID','Normal_FileName'};
        Header_T={'Tumor_CaseID','Tumor_FileName'};
        Sheet_N=[Header_N;S_Normal];
        Sheet_T=[Header_T;S_Tumor];
        
        % Write results data file %%
        Results_File_Path=cellstr(Results_File(1));
        UUID=cellstr(Results_File(2));
        % Normal sample sheet
        ResultsFilePath_N=strcat(char(Results_File_Path),'UnPaired_Normal_Sample_Sheet_',char(UUID),'.xlsx');
        xlswrite(ResultsFilePath_N,Sheet_N);
        % Tumor sample sheet
        ResultsFilePath_T=strcat(char(Results_File_Path),'UnPaired_Tumor_Sample_Sheet_',char(UUID),'.xlsx');
        xlswrite(ResultsFilePath_T,Sheet_T);
        
        Reselect_var=0;
    else
        % ONLY TUMOR
        %display("Your expression dataset does not contain normal samples; Reselect the onlytumor option"); 
        uiwait(msgbox('Your samples dataset does not contain normal samples; Reselect the onlytumor option'));
        Reselect_var=3;
    end
       
end
toc;
end
```
