## Supplementary material for "CanSeer: A Method for Development and Clinical Translation of Personalized Cancer Therapeutics": Main.m

```
%%%%%%%%%%%%%%%%%%%%%%%%%%%%                   Date:01-05-21        %%%%%%%%%%%%%%%%%%%%%%%%%%%%%%%%%%%%%%%%%%%%
%%%%%%%%%%%%%%%%%%%%%%%%%%%%%%%%%%%%%%%%%%%%%%%%%%%%%%%%%%%%%%%%%%%%%%%%%%%%%%%%%%%%%%%%%%%%%%%%%%%%%%%%%%%%%%%%
% Note:
% 1. Need path of the sample sheet that contains the information of FPKM expression files downloaded from https://portal.gdc.cancer.gov/
% 2. Need path of the network genes for which raw expression will be extracted
% 3. Need path of the copy number variations data file
% 4. Need path of the somatic mutations data file
% 5. Need path of the structural variants data file
% 6. Need path of the folder where FPKM text files are present(downloaded from https://portal.gdc.cancer.gov/ )
% Incase of missing path (data from 4 to 6 (CNV, Somatic Mutations or Structural variants)), set the path empty
% Path_SomaticMutations="";
% Path_StructuralVariants="";
% Path_Expression_Data="";
%%%%%%%%%%%%%%%%%%%%%%%%%%%%%%%%%%%%%%%%%%%%%%%%%%%%%%%%%%%%%%%%%%%%%%%%%%%%%%%%%%%%%%%%%%%%%%%%%%%%%%%%%%%%%%%%

clear all;
clc;
%% Time calculator %%
tic;

%% Set Path PC for lungs
% Path_Sample_Sheet="C:\Users\BIRL\Desktop\Pipeline_Normalization\Data\Lungs\Sample_Sheet_551.xlsx";
% Path_Network_Genes="C:\Users\BIRL\Desktop\Pipeline_Normalization\Data\Lungs\Network_Proteins_Genes_List.xlsx";
% Path_CNV="C:\Users\BIRL\Desktop\Pipeline_Normalization\Data\Lungs\lung_cancer_mutational_data\data_CNA_lung_cancer.txt";
% Path_SomaticMutations="C:\Users\BIRL\Desktop\Pipeline_Normalization\Data\Lungs\lung_cancer_mutational_data\data_somatic_mutations_lung_cancer.txt";
% Path_StructuralVariants="C:\Users\BIRL\Desktop\Pipeline_Normalization\Data\Lungs\lung_cancer_mutational_data\data_SV_lung_cancer.txt";
% Path_Expression_Data="C:\Users\BIRL\Desktop\Pipeline_Normalization\Data\Lungs\Expression_Data\";
% Results_File="C:\Users\BIRL\Desktop\Pipeline_Normalization\Results\Lung\";

% %% Set Path PC for breast  (When user wants to run Case Study I then please uncomment the below lines)
% Path_Sample_Sheet="C:\Users\rida\Desktop\brca_rb_4_3_22\Breast_Sample_Sheet.xlsx";
% Path_Network_Genes="C:\Users\rida\Desktop\brca_rb_4_3_22\Network_Proteins_Genes_List.xlsx";
% Path_CNV="C:\Users\rida\Desktop\brca_rb_4_3_22\Breast_Cancer_Mutational_Data\data_CNA.txt";
% Path_SomaticMutations="C:\Users\rida\Desktop\brca_rb_4_3_22\Breast_Cancer_Mutational_Data\data_mutations_extended.txt";
% Path_StructuralVariants="C:\Users\rida\Desktop\brca_rb_4_3_22\Breast_Cancer_Mutational_Data\data_fusions.txt";
% Path_Expression_Data="C:\Users\rida\Desktop\brca_rb_4_3_22\Expression_Data\";
% Results_File="C:\Users\rida\Desktop\brca_rb_4_3_22\Results\";

%% Set Path PC for Ovary
Path_Sample_Sheet="C:\Users\rida\Desktop\OV_NEW\Ovary_Sample_Sheet.xlsx";
Path_Network_Genes="C:\Users\rida\Desktop\OV_NEW\Network_Proteins_Genes_List.xlsx";
Path_CNV="C:\Users\rida\Desktop\OV_NEW\Ovarian_Cancer_Mutational_Data\data_CNA.txt";
Path_SomaticMutations="C:\Users\rida\Desktop\OV_NEW\Ovarian_Cancer_Mutational_Data\data_mutations_extended.txt";
Path_StructuralVariants="C:\Users\rida\Desktop\OV_NEW\Ovarian_Cancer_Mutational_Data\data_fusions.txt";
Path_Expression_Data="C:\Users\rida\Desktop\OV_NEW\Expression_Data\";
Results_File="C:\Users\rida\Desktop\OV_NEW\Results\";

%% User can opt for 'Paired', 'Unpaired' or 'Only Tumor' sample analysis %%
% If user select incorrect option, then will be asked to enter the correct option
test=1;
while test>0
    
    U_ID =  java.util.UUID.randomUUID;
    U_UID = char(U_ID.toString);
    Results_File_Path={Results_File,U_UID};
    
    prompt = {'Enter 1 for Paired, 2 for Unpaired, 3 for Only Tumor:'};
    dlgtitle = 'Input';
    dims = [1 70];
    definput = {'1 or 2 or 3'};
    answer = inputdlg(prompt,dlgtitle,dims,definput);
    NT_NAT_T=str2num(answer{1});
    
   %%% Note: Select 1 for paired samples, 2 for unpaired samples or 3 for onlytumor sample analysis %%%
   % NT_NAT_T = input('Enter 1 for Paired, 2 for Unpaired, 3 for Only Tumor: ');
    
    switch NT_NAT_T
        case 1
            %% Paired Sample Analysis %%
            %display('You have selected paired option');
            uiwait(msgbox('You have selected paired option'));
            % Step1: Filter sample_Sheet for paired samples
            [PairedSheet,a]=Filtering_CIDs_Paired(Path_Sample_Sheet,Results_File_Path);
            %[PairedSheet,a]=Filtering_CIDs_Paired(Path_Sample_Sheet);
            
            if(logical(1)== (~(isempty(PairedSheet))))
                % Step2: Extract expression for paired samples
                [NormalRawExpressionFile,TumorRawExpressionFile]=Extraction_of_Raw_Expression_Paired(Path_Network_Genes,PairedSheet,Path_Expression_Data,Results_File_Path);
              
                
                tryagain=1;
                while tryagain>0
                    
                    % Step3: Outliers detection
                    %%% Note: Select 1 for median absolute deviation(MAD) and 2 for inter quartile range (IQR) to detect outliers %%%
                    
                    prompt1 = {'Enter 1 for median absolute deviation (MAD) or 2 for inter quartile range(IQR) to detect outliers:'};
                    dlgtitle1 = 'Choice for outliers detection';
                    dims1 = [1 70];
                    definput1 = {'1 or 2'};
                    answer1 = inputdlg(prompt1,dlgtitle1,dims1,definput1);
                    MAD_OR_IQR=str2num(answer1{1});
        
                    %MAD_OR_IQR = input('Enter 1 for median absolute deviation (MAD) or 2 for inter quartile range(IQR) to detect outliers: ');
                    
                    switch MAD_OR_IQR
                        case 1
                            %display('You have selected median absolute deviation(MAD)');
                            uiwait(msgbox('You have selected median absolute deviation(MAD)'));
                            % Step3a: Detect outliers by applying median absolute deviation
                            [N_Outliers,T_Outliers,N_GenesOutliersInd,C_GenesOutliersInd]=Outliers_Detection_Using_Median_Absolute_Deviation(NormalRawExpressionFile,TumorRawExpressionFile,Results_File_Path);
                            
                            % Step4a: Filter copy number variations for network genes in paired samples
                            [CNV_LArray]= Filter_Copy_Number_Variations(Path_Network_Genes,PairedSheet,Path_CNV,Results_File_Path);
                            
                            % Step4b: Filter somatic mutations for network genes in paired samples
                            [SomaticMutation_LA]=Filter_SomaticMutations(Path_Network_Genes,PairedSheet,Path_SomaticMutations,Results_File_Path);
                            
                            % Step4c: Filter structural variations for network genes in paired samples
                            [StructuralVariants_LA]=Filter_Structural_Variants(Path_Network_Genes,PairedSheet,Path_StructuralVariants,Results_File_Path);
                            
                            % Step5: Normalize expression for network genes in paired samples
                            
                            [Median_Normalized_N,Median_Normalized_T]=Normalization_and_Median_Computation_Paired_Unpaired(NormalRawExpressionFile,TumorRawExpressionFile,N_Outliers,T_Outliers,CNV_LArray,SomaticMutation_LA,StructuralVariants_LA,Results_File_Path);
                            
                            tryagain=0;
                            
                        case 2
                            %display('You have selected inter quartile range(IQR)');
                            uiwait(msgbox('You have selected inter quartile range(IQR)'));
                            % Step3b: Detect outliers by applying inter-quartile range
                            [N_Outliers_Ind,N_Outliers,Normal_IQR,T_Outliers_Ind,T_Outliers,Tumor_IQR]=Outliers_Detection_Using_Inter_Quartile_Range_Paired_Unpaired(NormalRawExpressionFile,TumorRawExpressionFile,Results_File_Path);
                            
                            % Step4a: Filter copy number variations for network genes in paired samples
                            [CNV_LArray]= Filter_Copy_Number_Variations(Path_Network_Genes,PairedSheet,Path_CNV,Results_File_Path);
                            
                            % Step4b: Filter somatic mutations for network genes in paired samples
                            [SomaticMutation_LA]=Filter_SomaticMutations(Path_Network_Genes,PairedSheet,Path_SomaticMutations,Results_File_Path);
                            
                            % Step4c: Filter structural variations for network genes in paired samples
                            [StructuralVariants_LA]=Filter_Structural_Variants(Path_Network_Genes,PairedSheet,Path_StructuralVariants,Results_File_Path);
                            
                            % Step5: Normalize expression for network genes in paired samples
                            [Median_Normalized_N,Median_Normalized_T]=Normalization_and_Median_Computation_Paired_Unpaired(NormalRawExpressionFile,TumorRawExpressionFile,N_Outliers,T_Outliers,CNV_LArray,SomaticMutation_LA,StructuralVariants_LA,Results_File_Path);
                            
                            tryagain=0;
                            
                        case 3
                            display('Enter correct option');
                            tryagain=1;
                    end
                end
            else
                display('Your sample sheet is empty');
            end
            test=a;
            
        case 2
            %% UnPaired Sample Analysis %%
            display('You have selected unpaired option');
            
            % Step1: Filter sample_Sheet for unpaired samples
            [Sheet_Normal,Sheet_Tumor,a]=Filtering_CIDs_UnPaired(Path_Sample_Sheet,Results_File_Path);
            
            if ( logical(1)==(~(isempty(Sheet_Normal)) & (~(isempty(Sheet_Tumor)))))
                % Step2: Extract expression for unpaired samples
                [NormalRawExpressionFile,TumorRawExpressionFile]= Extraction_of_Raw_Expression_Unpaired(Path_Network_Genes,Sheet_Normal,Sheet_Tumor,Path_Expression_Data,Results_File_Path);
                
                
                tryagain=1;
                while tryagain>0
                    % Step3: Outliers detection
                    %%% Note: Select 1 for median absolute deviation(MAD) and 2 for inter quartile range (IQR) to detect outliers %%%
                    prompt1 = {'Enter 1 for median absolute deviation (MAD) or 2 for inter quartile range(IQR) to detect outliers:'};
                    dlgtitle1 = 'Choice for outliers detection';
                    dims1 = [1 70];
                    definput1 = {'1 or 2'};
                    answer1 = inputdlg(prompt1,dlgtitle1,dims1,definput1);
                    MAD_OR_IQR=str2num(answer1{1});
                    
                    %MAD_OR_IQR = input('Enter 1 for MAD or 2 for IQR to detect outliers: ');
                    
                    switch MAD_OR_IQR
                        case 1
                            %display('You have selected median absolute deviation(MAD)');
                            uiwait(msgbox('You have selected median absolute deviation(MAD)'));
                            % Step3a: Detect outliers by applying median absolute deviation
                            [N_Outliers,T_Outliers,N_GenesOutliersInd,C_GenesOutliersInd]=Outliers_Detection_Using_Median_Absolute_Deviation(NormalRawExpressionFile,TumorRawExpressionFile,Results_File_Path);
                            
                            % Step4a: Filter copy number variations for network genes in unpaired samples
                            [CNV_LArray]= Filter_Copy_Number_Variations(Path_Network_Genes,Sheet_Tumor,Path_CNV,Results_File_Path);
                            
                            % Step4b: Filter somatic mutations for network genes in unpaired samples
                            [SomaticMutation_LA]=Filter_SomaticMutations(Path_Network_Genes,Sheet_Tumor,Path_SomaticMutations,Results_File_Path);
                            
                            % Step4c: Filter structural variations for network genes in unpaired samples
                            [StructuralVariants_LA]=Filter_Structural_Variants(Path_Network_Genes,Sheet_Tumor,Path_StructuralVariants,Results_File_Path);
                            
                            % Step5: Normalize expression for network genes in unpaired samples
                            [Median_Normalized_N,Median_Normalized_T]=Normalization_and_Median_Computation_Paired_Unpaired(NormalRawExpressionFile,TumorRawExpressionFile,N_Outliers,T_Outliers,CNV_LArray,SomaticMutation_LA,StructuralVariants_LA,Results_File_Path);
                            
                            tryagain=0;
                            
                        case 2
                            %display('You have selected inter quartile range(IQR)')
                            uiwait(msgbox('You have selected inter quartile range(IQR)'));
                            % Step3b: Detect outliers by applying inter-quartile range
                            [N_Outliers_Ind,N_Outliers,Normal_IQR,T_Outliers_Ind,T_Outliers,Tumor_IQR]=Outliers_Detection_Using_Inter_Quartile_Range_Paired_Unpaired(NormalRawExpressionFile,TumorRawExpressionFile,Results_File_Path);
                            
                            % Step4a: Filter copy number variations for network genes in unpaired samples
                            [CNV_LArray]= Filter_Copy_Number_Variations(Path_Network_Genes,Sheet_Tumor,Path_CNV,Results_File_Path);
                            
                            % Step4b: Filter somatic mutations for network genes in unpaired samples
                            [SomaticMutation_LA]=Filter_SomaticMutations(Path_Network_Genes,Sheet_Tumor,Path_SomaticMutations,Results_File_Path);
                            
                            % Step4c: Filter structural variations for network genes in unpaired samples
                            [StructuralVariants_LA]=Filter_Structural_Variants(Path_Network_Genes,Sheet_Tumor,Path_StructuralVariants,Results_File_Path);
                            
                            % Step5: Normalize expression for network genes in unpaired samples
                            [Median_Normalized_N,Median_Normalized_T]=Normalization_and_Median_Computation_Paired_Unpaired(NormalRawExpressionFile,TumorRawExpressionFile,N_Outliers,T_Outliers,CNV_LArray,SomaticMutation_LA,StructuralVariants_LA,Results_File_Path);
                            
                            tryagain=0;
                            
                        case 3
                            display('Enter correct option');
                            tryagain=1;
                    end
                end
            else
                display('Your sample sheets are empty');
            end
            test=a;
            
        case 3
            %% OnlyTumor Sample Analysis %%
            display('You have selected onlytumor option');
            
            % Step1: Filter sample_Sheet for onlytumors
            [Sheet_Tumor,a]=Filtering_CIDs_Only_Tumor(Path_Sample_Sheet,Results_File_Path);
            
            if(logical(1)==(~(isempty(Sheet_Tumor))))
                % Step2: Extract expression data for onlytumor samples
                [TumorRawExpressionFile]= Extraction_of_Raw_Expression_Only_Tumor(Path_Network_Genes,Sheet_Tumor,Path_Expression_Data,Results_File_Path);

                 
                tryagain=1;
                while tryagain>0
                    
                    % Step3: Outlier detection
                    %%% Note: Select 1 for median absolute deviation(MAD) and 2 for inter quartile range (IQR) to detect outliers %%%
                    prompt1 = {'Enter 1 for median absolute deviation (MAD) or 2 for inter quartile range(IQR) to detect outliers:'};
                    dlgtitle1 = 'Choice for outliers detection';
                    dims1 = [1 70];
                    definput1 = {'1 or 2'};
                    answer1 = inputdlg(prompt1,dlgtitle1,dims1,definput1);
                    MAD_OR_IQR=str2num(answer1{1});
          
                    %MAD_OR_IQR = input('Enter 1 for MAD or 2 for IQR to detect outliers: ');
                    
                    switch MAD_OR_IQR
                        case 1
                            %display('You have selected median absolute deviation(MAD)');
                            uiwait(msgbox('You have selected median absolute deviation(MAD)'));
                            % Step3a: Detect outliers by applying median absolute deviation
                            [T_Outliers,T_GenesOutliersInd]=Outliers_Detection_Using_Median_Absolute_Deviation_Only_Tumor(TumorRawExpressionFile,Results_File_Path);
                            
                            % Step4a: Filter copy number variations for network genes in onlytumors samples
                            [CNV_LArray]= Filter_Copy_Number_Variations(Path_Network_Genes,Sheet_Tumor,Path_CNV,Results_File_Path);
                            
                            % Step4b: Filter somatic mutations for network genes in onlytumors samples
                            [SomaticMutation_LA]=Filter_SomaticMutations(Path_Network_Genes,Sheet_Tumor,Path_SomaticMutations,Results_File_Path);
                            
                            % Step4c: Filter structural variations for network genes in onlytumors samples
                            [StructuralVariants_LA]=Filter_Structural_Variants(Path_Network_Genes,Sheet_Tumor,Path_StructuralVariants,Results_File_Path);
                            
                            % Step5: Normalize expression for network genes in onlytumors samples
                            [Median_Normalized_T]=Normalization_and_Median_Computation_Only_Tumor(TumorRawExpressionFile,T_Outliers,CNV_LArray,SomaticMutation_LA,StructuralVariants_LA,Results_File_Path);
                            
                            tryagain=0;
                            
                        case 2
                            %display('You have selected inter quartile range(IQR)');
                            uiwait(msgbox('You have selected inter quartile range(IQR)'));
                            % Step3b: Detect outliers by applying inter-quartile range
                            [T_Outliers,T_Outliers_Ind,Tumor_IQR]=Outliers_Detection_Using_Inter_Quartile_Range_Only_Tumor(TumorRawExpressionFile,Results_File_Path);
                            
                            % Step4a: Filter copy number variations for network genes in onlytumors samples
                            [CNV_LArray]= Filter_Copy_Number_Variations(Path_Network_Genes,Sheet_Tumor,Path_CNV,Results_File_Path);
                            
                            % Step4b: Filter somatic mutations for network genes in onlytumors samples
                            [SomaticMutation_LA]=Filter_SomaticMutations(Path_Network_Genes,Sheet_Tumor,Path_SomaticMutations,Results_File_Path);
                            
                            % Step4c: Filter structural variations for network genes in onlytumors samples
                            [StructuralVariants_LA]=Filter_Structural_Variants(Path_Network_Genes,Sheet_Tumor,Path_StructuralVariants,Results_File_Path);
                            
                            % Step5: Normalize expression for network genes in onlytumors samples
                            [Median_Normalized_T]=Normalization_and_Median_Computation_Only_Tumor(TumorRawExpressionFile,T_Outliers,CNV_LArray,SomaticMutation_LA,StructuralVariants_LA,Results_File_Path);
                            
                            tryagain=0;
                            
                        case 3
                            %display('Enter correct option');
                            uiwait(msgbox('Enter correct option'));
                            tryagain=1;
                    end
                end
            else
                %display('Your sample sheet is empty');
                uiwait(msgbox('Your sample sheet is empty'));
            end
            test=a;
            
            %% Please select the right option
        case 4
            %disp('Reselect the right option');
            uiwait(msgbox('Reselect the right option'));
            test=4;
    end
    continue;
end

toc
%
```
