## Supplementary material for "CanSeer: A Method for Development and Clinical Translation of Personalized Cancer Therapeutics": Normalization_and_Median_Computation_Only_Tumor.m

```
% This function takes input parameters 
% 'TumorRawExpressionFile', 'T_Outliers', 'CNV_LArray', 'SomaticMutation_LA,'StructuralVariants_LA' and 'Results_File'
% Returns the median of normal abstracted from tumor samples and normalized data for tumor samples


% Steps:1 Detect outliers in tumor sampples
% Step2:  In tumor samples, check if outliers comes as a result of
%       1a) CNV    ab) SomaticMutation  2c) Structural Variation
%        Remove outliers that does not represent any of these three
%        mutations types and retain outliers if they contain any of the
%        three variations
% Step3: Now normalize the gene expression by looking at the max exp across
% all samples after removing outliers from 'T'
% Step4: Abstract Normal from tumor samples by checking one condition
% 1.Samples without these three mutations will be abstracted as
% Normal for every gene and take the median
% ~(CNV | SM |SV)
% NOTE: 'NaN' represents outliers in the Normalized data to keep cell array size consistent

function [Median_Normalized_Tumor]=Normalization_and_Median_Computation_Only_Tumor(TumorRawExpressionFile,T_Outliers,CNV_LArray,SomaticMutation_LA,StructuralVariants_LA,Results_File)
tic;

Median_N=[];

% Tumor Variable initialization
Tumor_Raw_Expression_File=TumorRawExpressionFile;
T_Outliers=T_Outliers;
CNV_LA=CNV_LArray;
SomaticMut_LA=SomaticMutation_LA;
Structural_Var_LA=StructuralVariants_LA;
[Row1_G, Col1_CID]=size(Tumor_Raw_Expression_File);
TumorRawExpression=Tumor_Raw_Expression_File(2:end,2:end);
CaseIDs_T=Tumor_Raw_Expression_File(1,2:end);
Sum_T_Outliers=[];
Normalized_T=[];
Median_T=[];

% Genes Cell Array
Net_Genes=TumorRawExpressionFile(2:end,1);

% Max will be the Max from both 'N' and 'T'
Max=[];
Count_Normal_Abstracted=[];
% Maximum number of samples in abstracted Normal genes
counter=0;
for j=1:length(Net_Genes)
    GeneExp_Vec_T=cell2mat(TumorRawExpression(j,:));
    CNV_LA_Cols=CNV_LA(j,:);
    SomaticMut_LA_Cols=SomaticMut_LA(j,:);
    SV_LA_Cols=Structural_Var_LA(j,:);
    Mutations=(CNV_LA_Cols | SomaticMut_LA_Cols | SV_LA_Cols);
    Index_Without_Mutations=find(~Mutations);
    Count_Normal_Abstracted=[Count_Normal_Abstracted;num2cell(length(Index_Without_Mutations))];
end
MaxsizeofNormal_Index=max(cell2mat(Count_Normal_Abstracted));
% Initialize a matrix to store the normalized expression of abstracted
% normal
Abstracted_Normal=zeros(length(Net_Genes),MaxsizeofNormal_Index);

% Execute number of genes times
for i=1:length(Net_Genes)
    % 01.b: Remove Outliers from Tumor Expression data & find max
    GeneExp_Vec_T=cell2mat(TumorRawExpression(i,:));
    T_Outliers_Cols= T_Outliers(i,:);
    CNV_LA_Cols=CNV_LA(i,:);
    SomaticMut_LA_Cols=SomaticMut_LA(i,:);
    SV_LA_Cols=Structural_Var_LA(i,:);
    FilterOutliersByCNV= T_Outliers_Cols & ~CNV_LA_Cols;
    FilterOutiersByMut= T_Outliers_Cols & ~SomaticMut_LA_Cols;
    FilterOutiersBySV= T_Outliers_Cols & ~SV_LA_Cols;
    RemoveOutliersT= FilterOutliersByCNV & FilterOutiersByMut & FilterOutiersBySV;
    % True outliers is an outlier without CNV or SM or SV and put []/NAn for
    % representation in the normalized data
    GeneExp_Vec_T(find(RemoveOutliersT))=[];
    MaxGeneExp_T = (max(GeneExp_Vec_T));
    Sum_Ind=find(RemoveOutliersT);
    Sum_T_Outliers=[Sum_T_Outliers;num2cell(length(Sum_Ind))];
    
    % 02: Compare both max values in 'N' and 'T' to normalize expression data by the largest
    % Max
    
    
    GeneExp_Vec_T=cell2mat(TumorRawExpression(i,:));
    % Normalized Tumor data by Tumor Max
    NormVec_T=GeneExp_Vec_T / MaxGeneExp_T;
    NormVec_T(find(RemoveOutliersT))=nan;
    Normalized_T=[Normalized_T;num2cell(NormVec_T)];
    
    % Find Median of Tumor data
    Median_Vec_T=median(NormVec_T,'omitnan');
    Median_T = [Median_T;num2cell(Median_Vec_T)];
    
    % Store Max in cell Array 'Max'
    Max_Temp=MaxGeneExp_T;
    Max=[Max;num2cell(Max_Temp)];
    
    
end


% Common header in T and Abs_N
Header1={'Network_Genes'};

%Header2=CaseIDs_N;
Header3={'Outliers_Count','Max','Median'};
Header4=CaseIDs_T;
Header_T=[Header1,Header4,Header3];
Median_Normalized_T=[Net_Genes,Normalized_T,Sum_T_Outliers,Max,Median_T];
%Median_Normalized_T=horzcat(horzcat(Net_Genes,Normalized_T),horzcat(Sum_T_Outliers,horzcat(Max,Median_T)));
Median_Normalized_Tumor=[Header_T;Median_Normalized_T];

%% Write results data file %%
Results_File_Path=cellstr(Results_File(1));
UUID=cellstr(Results_File(2));

% Tumor Raw Expression
ResultsFilePath_T=strcat(char(Results_File_Path),'Normalized_Expression_Tumor_',char(UUID),'.xlsx');
xlswrite(ResultsFilePath_T,Median_Normalized_Tumor);

toc;
end
```
