## Supplementary material for "CanSeer: A Method for Development and Clinical Translation of Personalized Cancer Therapeutics": Normalization_and_Median_Computation_Paired_Unpaired.m

```
% This function takes input parameters
% 'NormalRawExpressionFile','TumorRawExpressionFile', 'N_Outliers', 'T_Outliers', 'CNV_LArray ', 'SomaticMutation_LA' and 'Results_File'
% Returns the median and normalized data for Both Normal and Tumor samples

% Steps:
% Step1: In 'Normal Samples', remove outliers
% Step2: In Tumor Samples, Check if outliers comes as a result of 
%       2a) CNV    2b) SomaticMutation  2c) Structural Variation 
% Step3: Now normalize the gene expression by looking at the max exp across
% all samples after removing outliers from both 'N' and 'T' separately
% Step4: Take the median of Normal Samples
% NOTE: 'NaN' represents outliers in the Normalized data and they are kept
% to keep cell arrays consistent

function [Median_Normalized_Normal,Median_Normalized_Tumor]=Normalization_and_Median_Computation_Paired_Unpaired(NormalRawExpressionFile,TumorRawExpressionFile,N_Outliers,T_Outliers,CNV_LArray,SomaticMutation_LA,StructuralVariants_LA,Results_File)
tic;
% Normal variable initialization
NormalRawExpressionFile=NormalRawExpressionFile;
N_Outliers=N_Outliers;
[Row_G, Col_CID]=size(NormalRawExpressionFile);
NormalRawExpression=NormalRawExpressionFile(2:end,2:end);
CaseIDs_N=NormalRawExpressionFile(1,2:end);
Sum_N_Outliers=[];
Normalized_N=[];
Median_N=[];

% Tumor variable initialization
TumorRawExpressionFile=TumorRawExpressionFile;
T_Outliers=T_Outliers;
CNV_LA=CNV_LArray;
SomaticMut_LA=SomaticMutation_LA;
Structural_Var_LA=StructuralVariants_LA;
[Row1_G, Col1_CID]=size(TumorRawExpressionFile);
TumorRawExpression=TumorRawExpressionFile(2:end,2:end);
CaseIDs_T=TumorRawExpressionFile(1,2:end);
Sum_T_Outliers=[];
Normalized_T=[];
Median_T=[];

% Genes Cell Array
Net_Genes=NormalRawExpressionFile(2:end,1);

% Max will be the Max from both 'N' and 'T'
Max=[];

% Execute over the number of genes
for i=1:length(Net_Genes)
    % 01.a: Remove outliers from normal expression data & find max
    GeneExp_Vec_N=cell2mat(NormalRawExpression(i,:));
    N_Outliers_Cols= N_Outliers(i,:);
    GeneExp_Vec_N(find(N_Outliers(i,:)))=[];
    MaxGeneExp_N = (max(GeneExp_Vec_N));
    Sum_N_Outliers=[Sum_N_Outliers;num2cell(sum(N_Outliers(i,:)))];
    
    % 01.b: Remove Outliers from tumor expression data & find max
    GeneExp_Vec_T=cell2mat(TumorRawExpression(i,:));
    T_Outliers_Cols= T_Outliers(i,:);
    CNV_LA_Cols=CNV_LA(i,:);
    SomaticMut_LA_Cols=SomaticMut_LA(i,:);
    SV_LA_Cols=Structural_Var_LA(i,:);
    
    %%% Filter true outliers %%%
    % 1. outlier without CNV
    FilterOutliersByCNV= T_Outliers_Cols & ~ CNV_LA_Cols;
    % 2. outlier without somatic_mutations
    FilterOutiersByMut= T_Outliers_Cols & ~SomaticMut_LA_Cols;
    % 3. outlier without SV
    FilterOutiersBySV= T_Outliers_Cols & ~SV_LA_Cols;
    %%% Get the true outliers here  %%%
    RemoveOutliersT= FilterOutliersByCNV & FilterOutiersByMut & FilterOutiersBySV;
    % place []/Nan in the true outliers index to keep size of matrix
    % consistent
    GeneExp_Vec_T(find(RemoveOutliersT))=[];
    % Find the max gene expression across all patients for each gene
    MaxGeneExp_T = (max(GeneExp_Vec_T));
    Sum_Ind=find(RemoveOutliersT);
    % Store the number of outliers for each gene 
    Sum_T_Outliers=[Sum_T_Outliers;num2cell(length(Sum_Ind))];
    
    % 02: Compare both max values in 'N' and 'T' to normalize expression data by the largest
    % Max   
    Sum_N_Out=length(Sum_Ind);
    Total_CID=Col_CID - 1;
    if(MaxGeneExp_N > MaxGeneExp_T)

        GeneExp_Vec_N=cell2mat(NormalRawExpression(i,:));
        % Normalized Normal data by Normal Max
        NormVec_N=GeneExp_Vec_N / MaxGeneExp_N;
        NormVec_N(find(N_Outliers(i,:)))=nan;
        Normalized_N=[Normalized_N;num2cell(NormVec_N)];
        
        % Find Median of Normal data
        Median_Vec_N=median(NormVec_N,'omitnan');
        Median_N = [Median_N;num2cell(Median_Vec_N)];
        
        GeneExp_Vec_T=cell2mat(TumorRawExpression(i,:));
        % Normalized Tumor data by Normal Max
        NormVec_T=GeneExp_Vec_T / MaxGeneExp_N;
        NormVec_T(find(RemoveOutliersT))=nan;
        Normalized_T=[Normalized_T;num2cell(NormVec_T)];
        
        % Find Median of Tumor data
        Median_Vec_T=median(NormVec_T,'omitnan');
        Median_T = [Median_T;num2cell(Median_Vec_T)];
        
        % Store Max in cell Array 'Max'
        Max_Temp=MaxGeneExp_N;
        Max=[Max;num2cell(Max_Temp)];
    else
        % This is again intialized here just to enGeneExp_Vec_N
        GeneExp_Vec_N=cell2mat(NormalRawExpression(i,:)); 
        % Normalized Normal data by Tumor Max
        NormVec_N=GeneExp_Vec_N / MaxGeneExp_T;
        NormVec_N(find(N_Outliers(i,:)))=nan;
        Normalized_N=[Normalized_N;num2cell(NormVec_N)];
        
        % Find Median of Normal data
        Median_Vec_N=median(NormVec_N,'omitnan');
        Median_N = [Median_N;num2cell(Median_Vec_N)];
        
        GeneExp_Vec_T=cell2mat(TumorRawExpression(i,:));
        % Normalized Tumor data by Tumor Max
        NormVec_T=GeneExp_Vec_T / MaxGeneExp_T;
        NormVec_T(find(RemoveOutliersT))=nan;
        Normalized_T=[Normalized_T;num2cell(NormVec_T)];
        
        % Find Median of Tumor data
        Median_Vec_T=median(NormVec_T,'omitnan');
        Median_T = [Median_T;num2cell(Median_Vec_T)];      
        
        % Store Max in cell Array 'Max'
        Max_Temp=MaxGeneExp_T;
        Max=[Max;num2cell(Max_Temp)];
    end   
end
Header1={'Network_Genes'};
Header2=CaseIDs_N;
Header3={'Outliers_Count','Max','Median'};
Header4=CaseIDs_T;
Header_N=[Header1,Header2,Header3];
Header_T=[Header1,Header4,Header3];
Median_Normalized_N=horzcat(horzcat(Net_Genes,Normalized_N),horzcat(Sum_N_Outliers,horzcat(Max,Median_N)));
Median_Normalized_T=horzcat(horzcat(Net_Genes,Normalized_T),horzcat(Sum_T_Outliers,horzcat(Max,Median_T)));
Median_Normalized_Normal=[Header_N;Median_Normalized_N];
Median_Normalized_Tumor=[Header_T;Median_Normalized_T];

%% Write results data file %%
Results_File_Path=cellstr(Results_File(1));
UUID=cellstr(Results_File(2));
% Normal Raw Expression
ResultsFilePath_N=strcat(char(Results_File_Path),'Normalized_Expression_Normal_',char(UUID),'.xlsx');
xlswrite(ResultsFilePath_N,Median_Normalized_Normal);
% Tumor Raw Expression
ResultsFilePath_T=strcat(char(Results_File_Path),'Normalized_Expression_Tumor_',char(UUID),'.xlsx');
xlswrite(ResultsFilePath_T,Median_Normalized_Tumor);
    
end
```
