## Supplementary material for "CanSeer: A Method for Development and Clinical Translation of Personalized Cancer Therapeutics": Outliers_Detection_Using_Inter_Quartile_Range_Only_Tumor.m

```
%% This function takes parameters 'TumorRawExpressionFile' and 'Results_File'  
%% Returns the logical array of outliers for Tumor samples [Rows represent sorted list of Network Genes and Columns represents the order of CID in paired sample file]
%% Returns detailed IQR computation file for Tumor samples

function [T_Outliers,T_Outliers_Ind,Tumor_IQR]=Outliers_Detection_Using_Inter_Quartile_Range_Only_Tumor(TumorRawExpressionFile,Results_File)
tic;
% Get Tumor Raw expression as the input parameter
TumorRawExpression=TumorRawExpressionFile;
TRawExp=TumorRawExpression(2:end,2:end);
NUM1=cell2mat(TRawExp(:,:));
CCID=TumorRawExpression(1,2:end);
[TROWS,TCOLS]=size(NUM1);
T_Outliers=zeros(TROWS,TCOLS);

% Get Network Gene Names from the expression data file 
Net_Genes=TumorRawExpression(:,1);

% Variable Initialization for Tumor IQR computation
T_Q1=[];
T_Q3=[];
T_IQR=[];
T_LowerBound=[];
T_UpperBound=[];
T_OutliersSum=[];
C_=struct;
T_Count_Outliers=0;

% Compute this loop for number of network genes 'NROWS'
for i=1:TROWS
    % Get expression values from all the from each gene row 'i' and all patients columns ':'
    Out_temp1=NUM1(i,:);
    temp1=sort(Out_temp1);
    % Compute Q1, Q3 and IQR to find lower and upper bound to filter
    % outliers of every  gene across tumor samples
    TT_Q1= median(temp1(find(temp1<median(temp1))));
    TT_Q3= median(temp1(find(temp1>median(temp1))));
    TT_IQR= TT_Q3 - TT_Q1;
    TTemp= Out_temp1 < TT_Q1 - (1.5 * TT_IQR);
    TTemp1= Out_temp1 > TT_Q3 + (1.5 * TT_IQR);
    yy= TTemp + TTemp1;
    T_Outliers(i,:)= yy;
    Tsum=length(find(yy));
    
    % IQR Values stored in cell arrays
    T_Q1=[T_Q1,num2cell(TT_Q1)];
    T_Q3=[T_Q3,num2cell(TT_Q3)];
    T_IQR=[T_IQR,num2cell(TT_IQR)];
    T_LowerBound=[T_LowerBound,num2cell(TT_Q1 - (1.5 * TT_IQR))];
    T_UpperBound=[T_UpperBound,num2cell(TT_Q3 + (1.5 * TT_IQR))];
    T_OutliersSum=[T_OutliersSum,num2cell(Tsum)];
    
    % Find index of outliers of gene across Cancer Samples
    T_tempo=yy;
    Ind=find(T_tempo);
    Normal(i).GeneInd=num2cell(Ind);
    C_(i).Gene=Ind;
    [R,C]=size(C_(i).Gene);
    T_tempo=C;
    if(T_tempo>T_Count_Outliers)
        T_Count_Outliers=T_tempo;
    end   
end
% Concatenate IQR values with Expression file
Header={'Quartile1', 'Quartile3', 'IQR', 'LowerBound', 'UpperBound','SumOfOutliers'};
Tumor=[T_Q1;T_Q3; T_IQR;T_LowerBound; T_UpperBound;T_OutliersSum];
Tumor_IQR=horzcat(TumorRawExpressionFile,(vertcat(Header,transpose(Tumor))));

% Index of the outliers stored in matrix and '0' index in the matrix is not
% associated with any patient so ignore '0' values in the matrix
%Ignore zeros in this table
T_Outliers_Ind=zeros(TROWS,T_Count_Outliers);
for i=1: TROWS
    [R1,C1]=size(C_(i).Gene);
    T_Outliers_Ind(i,1:C1)=C_(i).Gene;
end

% Include Headers in the Outliers logical array
Header_Tumor=[CCID;num2cell(T_Outliers)];
Tumor_Outliers_LA=[Net_Genes,Header_Tumor];

%% Write results data file %%
Results_File_Path=cellstr(Results_File(1));
UUID=cellstr(Results_File(2));

% Tumor IQR and logical array
ResultsFilePath_T=strcat(char(Results_File_Path),'Tumor_Outliers_Logical_Array_IQR_',char(UUID),'.xlsx');
xlswrite(ResultsFilePath_T,Tumor_Outliers_LA);
ResultsFilePath_TT=strcat(char(Results_File_Path),'Tumor_IQR_',char(UUID),'.xlsx');
xlswrite(ResultsFilePath_TT,Tumor_IQR);

toc;
end
```
