## Supplementary material for "CanSeer: A Method for Development and Clinical Translation of Personalized Cancer Therapeutics": Outliers_Detection_Using_Median_Absolute_Deviation.m

```
%% This function takes parameters 'NormalRawExpressionFile','TumorRawExpressionFile' and 'Results_File'  
%% Returns the logical array of outliers for both Normal and Tumor samples [Rows represent sorted list of Network Genes and Columns represents the order of CID in paired sample file]
function [N_Outliers,T_Outliers,N_Outliers_Ind,T_Outliers_Ind]=Outliers_Detection_Using_Median_Absolute_Deviation(NormalRawExpressionFile,TumorRawExpressionFile,Results_File)
tic;
% Get Normal Raw expression and initialize variables
NormalRawExpression=NormalRawExpressionFile;
[Rows,Cols]=size(NormalRawExpression);
NRawExp=NormalRawExpression(2:end,2:end);
Temp=NRawExp(:,:);
NUM=cell2mat(Temp);
NCID=NormalRawExpression(1,2:end);
[NROWS,NCOLS]=size(NUM);
Normal=struct;
N_=struct;
N_Count_Outliers=0;

% Get Tumor Raw expression and initialize variables
TumorRawExpression=TumorRawExpressionFile;
TRawExp=TumorRawExpression(2:end,2:end);
Temp1=TRawExp(:,:);
NUM1=cell2mat(Temp1);
CCID=TumorRawExpression(1,2:end);
[NROWS1,NCOLS1]=size(NUM1);
Cancer=struct;
C_=struct;
T_Count_Outliers=0;

% Assigned network genes from the expression data file'Column1'
NetGenes=NormalRawExpression(2:end,1);

% isoutlier function by specifing '2' returns outliers across all columns (CID)
N_Outliers = isoutlier(NUM,2);
%NSumOutliers = sum(N_Outliers,2);

T_Outliers = isoutlier(NUM1,2);
%TSumOutliers = sum(T_Outliers,2);

for i=1: NROWS
    % Find index of outliers of gene across Normal Samples
    % returns logical array of outliers with '1' and non-outliers with '0'
    % for every gene(i)
    temp=N_Outliers(i,:);
    % find returns the index of '1' in the logical array
    Ind=find(temp);
    % storing indexes for every gene(i) in the struct 
    Normal(i).GeneInd=num2cell(Ind);
    N_(i).Gene=Ind;
    [R,C]=size(N_(i).Gene);
    Temp=C;
    if(Temp>N_Count_Outliers)
        N_Count_Outliers=Temp;
    end
    % Find index of outliers of gene across Cancer Samples
    temp1=T_Outliers(i,:);
    Ind1=find(temp1);
    Cancer(i).GeneInd=num2cell(Ind1);
    C_(i).Gene=Ind1;
    [R1,C1]=size(C_(i).Gene);
    Temp1=C1;
    if(Temp1>T_Count_Outliers)
        T_Count_Outliers=Temp1;
    end
    
end
%Ignore zeros in both the tables just for makiing matrix consistent interms
%of size
N_Outliers_Ind=zeros(NROWS,N_Count_Outliers);
T_Outliers_Ind=zeros(NROWS,T_Count_Outliers);
for i=1: NROWS
    [R,C]=size(N_(i).Gene);
    N_Outliers_Ind(i,1:C)=N_(i).Gene;
    [R1,C1]=size(C_(i).Gene);
    T_Outliers_Ind(i,1:C1)=C_(i).Gene;
end

% Include Headers in the outliers logical array 
Normal_Outliers= [NormalRawExpression(:,1),[NCID;num2cell(N_Outliers)]]; 
Tumor_Outliers= [TumorRawExpression(:,1),[CCID;num2cell(T_Outliers)]]; 

%% Write results data file %%
Results_File_Path=cellstr(Results_File(1));
UUID=cellstr(Results_File(2));
% Normal MAD logical array
ResultsFilePath_N=strcat(char(Results_File_Path),'Normal_Outliers_Logical_Array_MAD_',char(UUID),'.xlsx');
xlswrite(ResultsFilePath_N,Normal_Outliers);

% Tumor MAD logical array
ResultsFilePath_T=strcat(char(Results_File_Path),'Tumor_Outliers_Logical_Array_MAD_',char(UUID),'.xlsx');
xlswrite(ResultsFilePath_T,Tumor_Outliers);

toc;
end
```
