## Supplementary material for "CanSeer: A Method for Development and Clinical Translation of Personalized Cancer Therapeutics": Outliers_Detection_Using_Median_Absolute_Deviation_Only_Tumor.m

```
%% This function takes parameters 'TumorRawExpressionFile' and 'Results_File'
%% Returns the logical array of outliers in Tumor samples [Rows represent sorted list of Network Genes and Columns represents the CASEIDs
function [T_Outliers,T_Outliers_Ind]=Outliers_Detection_Using_Median_Absolute_Deviation_Only_Tumor(TumorRawExpressionFile,Results_File)
tic;
% Get Tumor Raw expression and initialize variables
TumorRawExpression=TumorRawExpressionFile;
TRawExp=TumorRawExpression(2:end,2:end);
Temp1=TRawExp(:,:);
NUM1=cell2mat(Temp1);
CCID=TumorRawExpression(1,2:end);
[NROWS1,NCOLS1]=size(NUM1);
Cancer=struct;
C_=struct;
T_Count_Outliers=0;

% Assigned network genes from the expression data file'Column1'
NetGenes=TumorRawExpression(2:end,1);
% isoutlier function with 2 returns outliers across caseid for all the genes
T_Outliers = isoutlier(NUM1,2);

for i=1:length(NetGenes)
    % Find index of outliers of gene across tumor samples
    temp1=T_Outliers(i,:);
    % Find returns the index of outliers from the logical array 'temp1'
    Ind1=find(temp1);
    Cancer(i).GeneInd=num2cell(Ind1);
    C_(i).Gene=Ind1;
    [R1,C1]=size(C_(i).Gene);
    Temp1=C1;
    if(Temp1>T_Count_Outliers)
        T_Count_Outliers=Temp1;
    end
    
end
% Ignore zeros in both the tables just for makiing matrix consistent interms of size
T_Outliers_Ind=zeros(length(NetGenes),T_Count_Outliers);
for i=1: length(NetGenes)
    [R1,C1]=size(C_(i).Gene);
    T_Outliers_Ind(i,1:C1)=C_(i).Gene;
end

% Include Headers in the outliers logical array
Tumor_Outliers= [TumorRawExpression(:,1),[CCID;num2cell(T_Outliers)]];


%% Write results data file %%
Results_File_Path=cellstr(Results_File(1));
UUID=cellstr(Results_File(2));

% Only Tumor MAD logical array
ResultsFilePath_T=strcat(char(Results_File_Path),'Tumor_Outliers_Logical_Array_MAD_',char(UUID),'.xlsx');
xlswrite(ResultsFilePath_T,Tumor_Outliers);

toc;
end
```
