## Supplementary material for "CanSeer: A Method for Development and Clinical Translation of Personalized Cancer Therapeutics": progressbar.m

```
function progressbar(varargin)

%{

        Copyright (c) 2005, Steve Hoelzer
        All rights reserved.

        Redistribution and use in source and binary forms, with or without
        modification, are permitted provided that the following conditions are
        met:

            * Redistributions of source code must retain the above copyright
              notice, this list of conditions and the following disclaimer.
            * Redistributions in binary form must reproduce the above copyrights
              notice, this list of conditions and the following disclaimer in
              the documentation and/or other materials provided with the distribution

        THIS SOFTWARE IS PROVIDED BY THE COPYRIGHT HOLDERS AND CONTRIBUTORS "AS IS"
        AND ANY EXPRESS OR IMPLIED WARRANTIES, INCLUDING, BUT NOT LIMITED TO, THE
        IMPLIED WARRANTIES OF MERCHANTABILITY AND FITNESS FOR A PARTICULAR PURPOSE
        ARE DISCLAIMED. IN NO EVENT SHALL THE COPYRIGHT OWNER OR CONTRIBUTORS BE
        LIABLE FOR ANY DIRECT, INDIRECT, INCIDENTAL, SPECIAL, EXEMPLARY, OR
        CONSEQUENTIAL DAMAGES (INCLUDING, BUT NOT LIMITED TO, PROCUREMENT OF
        SUBSTITUTE GOODS OR SERVICES; LOSS OF USE, DATA, OR PROFITS; OR BUSINESS
        INTERRUPTION) HOWEVER CAUSED AND ON ANY THEORY OF LIABILITY, WHETHER IN
        CONTRACT, STRICT LIABILITY, OR TORT (INCLUDING NEGLIGENCE OR OTHERWISE)
        ARISING IN ANY WAY OUT OF THE USE OF THIS SOFTWARE, EVEN IF ADVISED OF THE
        POSSIBILITY OF SUCH DAMAGE.

%}

persistent progfig progdata lastupdate

% Get inputs
if nargin > 0
    input = varargin;
    ninput = nargin;
else
    % If no inputs, init with a single bar
    input = {0};
    ninput = 1;
end

% If task completed, close figure and clear vars, then exit
if input{1} == 1
    if ishandle(progfig)
        delete(progfig) % Close progress bar
    end
    clear progfig progdata lastupdate % Clear persistent vars
    drawnow;
    return
end

% Init reset flag
resetflag = false;

% Set reset flag if first input is a string
if ischar(input{1})
    resetflag = true;
end

% Set reset flag if all inputs are zero
if input{1} == 0
    % If the quick check above passes, need to check all inputs
    if all([input{:}] == 0) && (length([input{:}]) == ninput)
        resetflag = true;
    end
end

% Set reset flag if more inputs than bars
if ninput > length(progdata)
    resetflag = true;
end

% If reset needed, close figure and forget old data
if resetflag
    if ishandle(progfig)
        delete(progfig) % Close progress bar
    end
    progfig = [];
    progdata = []; % Forget obsolete data
end

% Create new progress bar if needed
if ishandle(progfig)
else % This strange if-else works when progfig is empty (~ishandle() does not)
    
    % Define figure size and axes padding for the single bar case
    height = 0.035;
    width = height * 8;
    hpad = 0.02;
    vpad = 0.15;
    
    % Figure out how many bars to draw
    nbars = max(ninput, length(progdata));
    
    % Adjust figure size and axes padding for number of bars
    heightfactor = (1 - vpad) * nbars + vpad;
    height = height * heightfactor;
    vpad = vpad / heightfactor;
    
    % Initialize progress bar figure
    left = (1 - width) / 2;
    bottom = (1 - height) / 2;
    progfig = figure(...
        'Units', 'normalized',...
        'Position', [left bottom width height],...
        'NumberTitle', 'off',...
        'Resize', 'off',...
        'MenuBar', 'none' );
    
    % Initialize axes, patch, and text for each bar
    left = hpad;
    width = 1 - 2*hpad;
    vpadtotal = vpad * (nbars + 1);
    height = (1 - vpadtotal) / nbars;
    for ndx = 1:nbars
        % Create axes, patch, and text
        bottom = vpad + (vpad + height) * (nbars - ndx);
        progdata(ndx).progaxes = axes( ...
            'Position', [left bottom width height], ...
            'XLim', [0 1], ...
            'YLim', [0 1], ...
            'Box', 'on', ...
            'ytick', [], ...
            'xtick', [] );
        progdata(ndx).progpatch = patch( ...
            'XData', [0 0 0 0], ...
            'YData', [0 0 1 1] );
        progdata(ndx).progtext = text(0.99, 0.5, '', ...
            'HorizontalAlignment', 'Right', ...
            'FontUnits', 'Normalized', ...
            'FontSize', 0.7 );
        progdata(ndx).proglabel = text(0.01, 0.5, '', ...
            'HorizontalAlignment', 'left', ...
            'FontUnits', 'Normalized', ...
            'FontSize', 0.7 );
        if ischar(input{ndx})
            set(progdata(ndx).proglabel, 'String', input{ndx})
            input{ndx} = 0;
        end
        
        % Set callbacks to change color on mouse click
        set(progdata(ndx).progaxes, 'ButtonDownFcn', {@changecolor, progdata(ndx).progpatch})
        set(progdata(ndx).progpatch, 'ButtonDownFcn', {@changecolor, progdata(ndx).progpatch})
        set(progdata(ndx).progtext, 'ButtonDownFcn', {@changecolor, progdata(ndx).progpatch})
        set(progdata(ndx).proglabel, 'ButtonDownFcn', {@changecolor, progdata(ndx).progpatch})
        
        % Pick a random color for this patch
        changecolor([], [], progdata(ndx).progpatch)
        
        % Set starting time reference        
        if ~isfield(progdata(ndx), 'starttime') || isempty(progdata(ndx).starttime)
            progdata(ndx).starttime = clock;
        end
        
    end
    % Set time of last update to ensure a redraw
    lastupdate = clock - 1;
    
end

% Process inputs and update state of progdata
for ndx = 1:ninput
    if ~isempty(input{ndx})
        progdata(ndx).fractiondone = input{ndx};
        progdata(ndx).clock = clock;
    end
end

% Enforce a minimum time interval between graphics updates
myclock = clock;
if abs(myclock(6) - lastupdate(6)) < 0.01 % Could use etime() but this is faster
    return
end

% Update progress patch
for ndx = 1:length(progdata)
    set(progdata(ndx).progpatch, 'XData', ...
        [0, progdata(ndx).fractiondone, progdata(ndx).fractiondone, 0])
end

% Update progress text if there is more than one bar
if length(progdata) > 1
    for ndx = 1:length(progdata)
        set(progdata(ndx).progtext, 'String', ...
            sprintf('%1d%%', floor(100*progdata(ndx).fractiondone)))
    end
end

% Update progress figure title bar
if progdata(1).fractiondone > 0
    runtime = etime(progdata(1).clock, progdata(1).starttime);
    time_elapsed = runtime * progdata(1).fractiondone + runtime; 
    time_elapsed_str = sec2timestr(time_elapsed);
    titlebarstr = sprintf('%2d%%    Elapsed Time :  %s ', ...
        floor(100*progdata(1).fractiondone), time_elapsed_str);
else
    titlebarstr = ' 0%';
end
set(progfig, 'Name', titlebarstr)

% Force redraw to show changes
drawnow;

% Record time of this update
lastupdate = clock;


% ------------------------------------------------------------------------------
    function changecolor(h, e, progpatch) %#ok<INUSL>
        % Change the color of the progress bar patch
        
        % Prevent color from being too dark or too light
        colormin = 1.5;
        colormax = 2.8;
        
        thiscolor = rand(1, 3);
        while (sum(thiscolor) < colormin) || (sum(thiscolor) > colormax)
            thiscolor = rand(1, 3);
        end
        
        set(progpatch, 'FaceColor', thiscolor)
        
    end


% ------------------------------------------------------------------------------
    function timestr = sec2timestr(sec)
        % Convert a time measurement from seconds into a human readable string.
        
        % Convert seconds to other units
        w = floor(sec/604800); % Weeks
        sec = sec - w*604800;
        d = floor(sec/86400); % Days
        sec = sec - d*86400;
        h = floor(sec/3600); % Hours
        sec = sec - h*3600;
        m = floor(sec/60); % Minutes
        sec = sec - m*60;
        s = floor(sec); % Seconds
        
        % Create time string
        if w > 0
            if w > 9
                timestr = sprintf('%d week', w);
            else
                timestr = sprintf('%d week, %d day', w, d);
            end
        elseif d > 0
            if d > 9
                timestr = sprintf('%d day', d);
            else
                timestr = sprintf('%d day, %d hr', d, h);
            end
        elseif h > 0
            if h > 9
                timestr = sprintf('%d hr', h);
            else
                timestr = sprintf('%d hr, %d min', h, m);
            end
        elseif m > 0
            if m > 9
                timestr = sprintf('%d min', m);
            else
                timestr = sprintf('%d min, %d sec', m, s);
            end
        else
            timestr = sprintf('%d sec', s);
        end
        
    end

end
```
